# Linked *OXTR* Variants Are Associated With Social Behavior Differences in Bonobos (*Pan paniscus*)

**DOI:** 10.1101/2023.12.22.573122

**Authors:** Sara A. Skiba, Ryan McCall, Azeeza Byers, Sarah Waldron, Amanda J. Epping, Jared P. Taglialatela

## Abstract

Single-nucleotide polymorphisms (SNPs) in forkhead box protein P2 (*FOXP2*), oxytocin receptor (*OXTR*), and arginine vasopressin receptor gene 1A (*AVPR1A*) have been associated with linguistic and social development in humans, as well as symptom severity in autism. Studying biobehavioral mechanisms in the species most closely related to humans can provide insights into the origins of human communication, and the impact of genetic variation on complex behavioral phenotypes. Here, we tested the hypothesis that similar genetic factors underlie social communication differences in both bonobos (*Pan paniscus*) and humans. We analyzed Sanger sequencing results to determine if bonobos exhibit individual variation at 10 loci across *FOXP2*, *OXTR*, and *AVPR1A* that have been implicated in human social development and behavior. We identified a novel variant in bonobo *FOXP2*, as well as three novel variants in bonobo *OXTR* that were 19-184 base pairs away from the target human SNPs. We also found a linked SNP combination (TGA) across the 3 novel bonobo *OXTR* sites at high frequency (65%) in the study population, including 6 homozygous bonobos. When comparing the combined *OXTR* genotypes, we found significant group differences in social behavior; bonobos with two copies of the TGA combination were more social than bonobos with one copy and bonobos with zero copies of the TGA combination. Taken together, our findings suggest that these *OXTR* variants may influence individual-level social behavior in bonobos and support the notion that linked genetic variants are promising biomarkers for differences in human social communication.

**Revision Summary:** The original manuscript version underwent peer-review at Frontiers in Behavioral Neuroscience. As a part of this process, a reviewer requested that the sequence files be included in the manuscript. While preparing these files, Dr. Hudson identified several errors in his work regarding the individual bonobo genotypes reported in the original version of the manuscript. Upon discovery of the errors, Dr. Skiba and Dr. Taglialatela conducted a new genetic analysis using the raw files and subsequently corrected the misreported information - updating the editorial team and including the raw sequences with the revised manuscript. In addition to requested edits for improvement and clarity by the peer-reviewers, this revised manuscript corrects several errors in the human and bonobo reference genomes, the SNP locations, and the individual genotypes reported in the original manuscript. Information in the figures, main text, and supplementary materials has been corrected in this revised version. Dr. Hudson and his lab member, Mr. Hansen, were unable to identify how the errors in data reporting occurred, and neither party contributed to the revised version of this manuscript. The co-author list has been adjusted to reflect contributions to the corrected analyses and revised manuscript. We are deeply grateful to the editorial team and reviewers at Frontiers in Behavioral Neuroscience for their thorough peer-review of this work - improving the overall quality of the manuscript and leading to the discovery and correction of substantial errors in the original version.

## 1 Introduction

Autism spectrum disorder (ASD) is characterized by social communication deficits and restricted, repetitive behaviors (RRBs) that impact daily functioning and can persist into adulthood. Determining the genetic factors that impact individual-level social communication is critical to our understanding of ASD and other neurodevelopmental disorders and may aid in identifying children at risk of developing social and linguistic impairments. The Simons Foundation created a database of genes associated with aspects of the ASD behavioral phenotype – SFARI Gene [1] – providing a systematic assessment of evidence for individual ASD-related genes [2]. This growing database highlights the complexity of human neurodevelopmental disorders, like autism, and how challenging it can be to identify biomarkers in humans. To help address the potential for false positives in large genetic datasets, the SFARI gene scoring system ranks genes as 1, 2, or 3 (high confidence, strong candidate, or suggestive evidence) based on the strength of the evidence supporting the gene’s link to autism risk, as well as the frequency of replicated results [1,2].

A potential biological factor underlying individual differences in social communication is the forkhead box protein P2 (*FOXP2*; SFARI Gene Score 1). *FOXP2* is one of the first genes to be associated with human language disorders and fine orofacial motor control [3–5]. Most notably, researchers have determined that *FOXP2* is critical to the developmental processes underlying speech and language [3,4]. In addition, Haghighatfard and colleagues [6] found that lower *FOXP2* expression was associated with executive dysfunction in children diagnosed with ASD. Thus, it is possible that polymorphisms in *FOXP2* impact gene regulation and protein expression, and therefore may underlie individual-level differences in social communication [7].

Much like in the case of *FOXP2*, several studies have demonstrated the important role that the oxytocin receptor gene (*OXTR*; SFARI Gene Score 2) plays in social bond formation and social motivation [8–11]. In particular, researchers have documented relations between OXTR variation and social affiliation [12], vocal symptoms [13], as well as social communication impairments associated with ASD [14]. There is also considerable evidence that *OXTR* SNPs are related to empathy, prosocial temperament, social sensitivity, and stress reactivity in individuals diagnosed with ASD [15]. Variations in the arginine vasopressin receptor gene 1A (*AVPR1A*; SFARI Gene Score 2) have also been linked to differences in social communication in both humans and nonhuman mammals [16–22]. Over the course of hominid evolution, there is evidence indicating a tandem duplication of a non-coding, 5’ flanking region of *AVPR1A* (DupB) [17] in the five great apes (orangutans, gorillas, bonobos, and humans). The DupB region (∼350BP) contains a complex (CT)4-(TT)-(CT)8-(GT)24 polymorphic variable motif, known as the RS3 microsatellite [17]. In their 2008 study [17], Donaldson and colleagues found that among hominids (orangutans, gorillas, chimpanzees, bonobos, and humans), chimpanzees are unique in that individuals either have two copies, one copy, or zero copies of the DupB microsatellite-containing element in the 5’ flanking region of *AVPR1A*. Hopkins and colleagues found evidence that chimpanzees lacking the DupB region perform significantly poorer on social cognition tasks [19] and responded less to joint attention cues [23] than chimpanzees with at least one copy of the DupB region.

In contrast to the three DupB genotypes observed in chimpanzees, bonobos and humans are homozygous for the microsatellite (DupB+). While all bonobos and humans exhibit both copies of the DupB region, individuals vary in the length of the RS3 microsatellite [17]. In humans, allelic variations in *AVPR1A* RS3 microsatellite length have been associated with social behavior and pair-bonding, as well as individual differences in aspects of social cognition [14,22,24,25]. Specifically, individuals with a shorter RS3 microsatellite were less likely to act altruistically and reported higher levels of social conflict with their siblings, than did individuals with a longer RS3 microsatellite length [16,26]. All told, converging data across multiple disciplines have demonstrated that *AVPR1A* variations are associated with individual variability in social behavior, communication, and social cognition in humans and nonhuman great apes [16–17,21,27].

Collectively, these studies suggest that variation in *FOXP2*, *OXTR*, and *AVPR1A* influence social communication development and social functioning in humans. To better understand the impact of genetic variants on complex behavioral phenotypes, burgeoning evidence supports the study of these biobehavioral mechanisms in nonhuman animals [28–30]. Indeed, variation in *FOXP2*, *OXTR*, and *AVPR1A* have been associated with differences in social behavior and/or social communication in rodents [31,32], zebrafish [33], great apes [17,20,21,34,35], and zebra finches [36]. Seminal work in rodents, including knock-out experiments, account for much of what we know about ASD-associated variants and other social communication related genes [37–41]. However, recent evidence suggests that common animal models of neurodevelopmental disorders (e.g., rodents and invertebrates) are limited in their comparability to human social communication [42,43] and may be too phylogenetically distant from humans to advance early identification and intervention techniques [44].

Bonobos, along with chimpanzees, are the closest living relatives to humans [45] and are regarded as having one of the most complex social communication systems in the animal kingdom [46,47]. Although bonobos cannot be diagnosed with human neurodevelopmental disorders, they do show significant individual-level variability in social engagement [48–50], communicative production [51,52], and repetitive behaviors [50,53]. Several studies have identified links between ASD-associated genes and social communication in bonobos [9,54,55], as well as in chimpanzees [56–58]. There is also evidence that oxytocin is related to socio-sexual behavior in female bonobos [59]. Thus, bonobos are a promising model species for investigating the impact of genetic variants on human neurodevelopment and social communication.

Identifying biological factors that underlie individual-level social communication is critical to our understanding of autism and other neurodevelopmental disorders. To this end, we aimed to determine if bonobos - our closest living relatives - exhibit single nucleotide variation in *FOXP2, OXTR*, and *AVPR1A*, which are implicated in autism or differences in social communication. Given that bonobos live in large, dynamic social groups and that they produce complex communicative signals of various types (vocalizations, facial expressions, manual gestures, and multi-source signals), we hypothesized that bonobos may exhibit allelic variation in *FOXP2*, *OXTR*, and *AVPR1A*.

## 2 Materials and methods

### 2.1 Genetic analyses

Biological samples were collected from 29 bonobos (7 whole blood samples and 22 buccal samples) living at the Ape Cognition and Conservation Initiative (n=7; IACUC protocol #210305-01), the Columbus Zoo and Aquarium (n=6), and the Milwaukee County Zoo (n=16). Whole blood samples were collected under anesthesia during the bonobo’s routine physical exam (n=6), except for one blood sample that was collected from an awake bonobo via voluntary presentation (n=1). Buccal samples were collected by swabbing the inner cheek or lower lip for 10-15 seconds with a QIAGEN OmniSwab (n=22). All buccal samples were taken from awake bonobos that presented voluntarily for sample collection. Of the 22 buccal samples, 16 yielded DNA of a quality that was too low for Polymeric Chain Reaction (PCR) amplification. This was likely due to the level of bacteria in the buccal cavity of the bonobos, as well as the collection method; samples from Ape Initiative were repeated following the first attempt at DNA extraction and were collected from the bonobos’ lower lips after rinsing their mouths with water (as opposed to collecting the samples from the bonobos’ inner cheeks without rinsing with water beforehand). These repeated samples were sufficient for DNA extraction, resulting in a total of 13 biological samples for the genetic analyses (whole blood n=7, buccal swab n=6).

Autism-associated genes included in this study were *FOXP2*, *OXTR*, and *AVPR1A*. Primer pairs were designed using NCBI Primer-BLAST [60] and ApE [61] to flank each SNP by ∼250 base pairs (giving approximately 500bp amplicons) and ordered from Thermo Fisher Scientific. Primer pairs that were included in the study met the following criteria: 1) the PCR product was at least 250 base pairs long, 2) the melting temperature (Tm) of the primer was 60°C and the GC% was approximately 50%, and 3) the primer did not have hair-pin formations or self-annealing sites and was not 3’ prime complimentary. This resulted in the inclusion of 10 human loci: *FOXP2* rs6980093, *OXTR* rs2270463, rs237877, rs237878, rs35062132, rs2254295, rs237894, rs237895, and rs237900, and *AVPR1A* rs3803107 (Table 1). See Supplementary Material 1 for a list of human SNPs that did not meet the criteria.

**Table 1:**
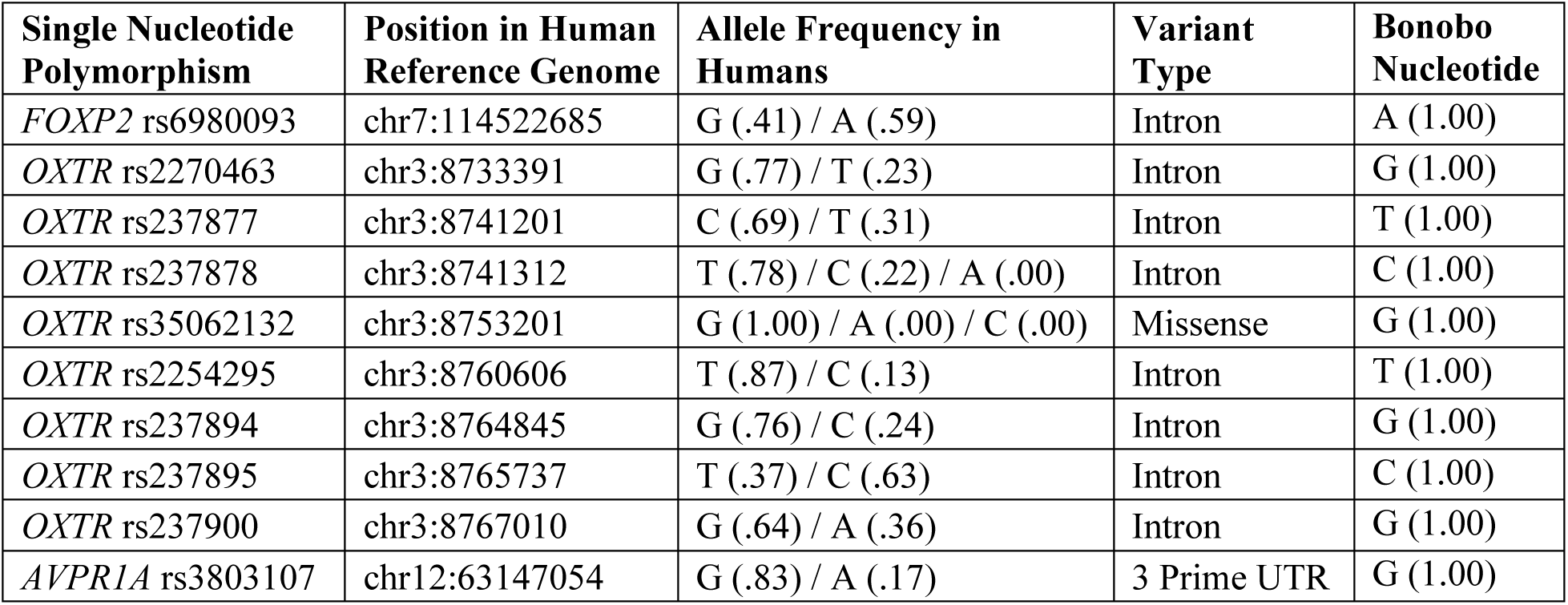
Selected Human SNPs: Name, position, relative allele frequency, and variant type for the selected human single nucleotide polymorphisms. Human alleles that were only found in a small subset of studies have rounded frequences of zero (.00).

DNA was extracted from bonobo whole blood and buccal samples, amplified by PCR, resolved by gel electrophoresis, gel purified (Zymo Gel DNA Recovery Kit) and sent to Genewiz for Sanger sequencing (see Supplementary Material 2 for details). Sequences for the bonobo reference genome (May 2020 (Muhdiblu_PPA_v0/panPan3)) and the human reference genome (Dec. 2013 (GRCh38/hg38)) were obtained from the UCSC Genome Browser [62]. To determine SNP presence, individual bonobo Sanger sequences were aligned to the other bonobos from our sample, the bonobo reference genome, and the human reference genome, using Clustal Omega [63]. See Supplementary Material 3 for the novel bonobo SNP visualizations in Clustal Omega, and Supplementary Material 4 for the raw bonobo sequences and reference sequences used in the alignments. Heterozygotes were identified by visual inspection of the Sanger chromatograms in 4Peaks [64] and confirmed via peak height ratios >33% to minimize the likelihood of false positives. Peak height ratios were calculated by dividing the height of the smaller peak by the height of the larger peak. Individual genotypes were cross-checked based on known pedigrees. See Supplementary Material 5 for the relatedness between each subject.

### 2.2 Behavioral data

To determine if any observed genetic variation was linked to social behavior, we utilized previously collected observational data that were available for 12 of the 13 subjects [50,65]. Focal observations were not available for subject M15 because the individual was under the age threshold of 2 years at the time of data collection. Bonobos included in the behavioral analyses ranged in age from 6 to 40 years old, with a mean age of 20.88 years (female n=6, female mean age=21.58 years; male n=6, male mean age=20.17 years). The bonobos were housed in their indoor and outdoor enclosures, which featured environmental enrichment (e.g., artificial vines and sleeping platforms) that promoted species-typical behaviors (e.g., climbing and nest building). Facility care staff were responsible for monitoring the health and well-being of the bonobos, and any behavioral concerns or signs of distress noted by the researchers were reported immediately to the care staff. No modifications were made to the animals’ housing or social groupings for this study, and data collection did not interfere in any way with the bonobo/care staff’s daily schedule.

For this study, we utilized eight 10-minute focal observations (i.e., observing the behavior of a single individual) from each subject to assess group differences in communicative production and in social proximity – an established method for measuring social relationships in nonhuman primates that encapsulates both social interest and engagement [66–68]. Social proximity was recorded instantaneously at 1-minute increments starting at time zero, which resulted in 11 proximity data points per observation. Communicative production (i.e., the total number of gestures, facial expressions, and multi-source signals, including vocalizations + gestures, gestures + facial expressions, and facial expressions + vocalizations, produced by the focal individual) was recorded on an all-occurrence basis. See Table 2 for social proximity measures.

**Table 2:** Social Proximity Measures.

| Proximity Measure | Description |
| --- | --- |
| Close-Touching | The focal individual is <1.5 meters from the nearest conspecific. |
| Socially Close | The focal individual is 1.5-3 meters from the nearest conspecific. |
| Socially Peripheral | The focal individual is 3-5 meters from the nearest conspecific. |
| Isolated | The focal individual is >5 meters from the nearest conspecific. |

### 2.3 Statistical analyses

For each observation, the total number of communicative signals produced was summed, and a social proximity score (ranging from 0-3), was calculated using the following formula, where 11 is the total number of proximity data points per focal follow:

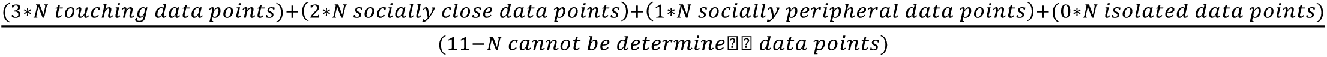

Kruskal-Wallis H tests and Wilcoxon rank tests were utilized to assess group differences in social behavior and communicative production. To rule out rearing influences on any observed differences in bonobo social behavior, we grouped bonobos based on whether they were raised primarily by their mother (typical rearing, n=7), or were raised primarily by humans and/or captured from the wild (atypical rearing, n=5). To rule out age influences on any observed differences in bonobo social behavior, we grouped bonobos based on whether they were a subadult (<14 years old, n=4), adult (14-34 years old, n=6), or elder (>34 years old, n=2), at the time the behavioral observations were collected.

## 3 Results

### 3.1 Genetic variation

Analyses revealed a novel SNP in bonobo *FOXP2*, 78bp downstream of *FOXP2* rs6980093 (human chr7:114522685; A/G). Five bonobos were heterozygous at this location (G/A), one bonobo was homozygous (G/G), and the rest were homozygous (A/A; bonobo 5’ flanking sequence CACTCGTATCACATTATAAT A/G; Figure 1). Analyses did not reveal individual variation in *AVPR1A*; all of the bonobos were homozygous for the same allele at this position (G/G).

**Figure 1.**
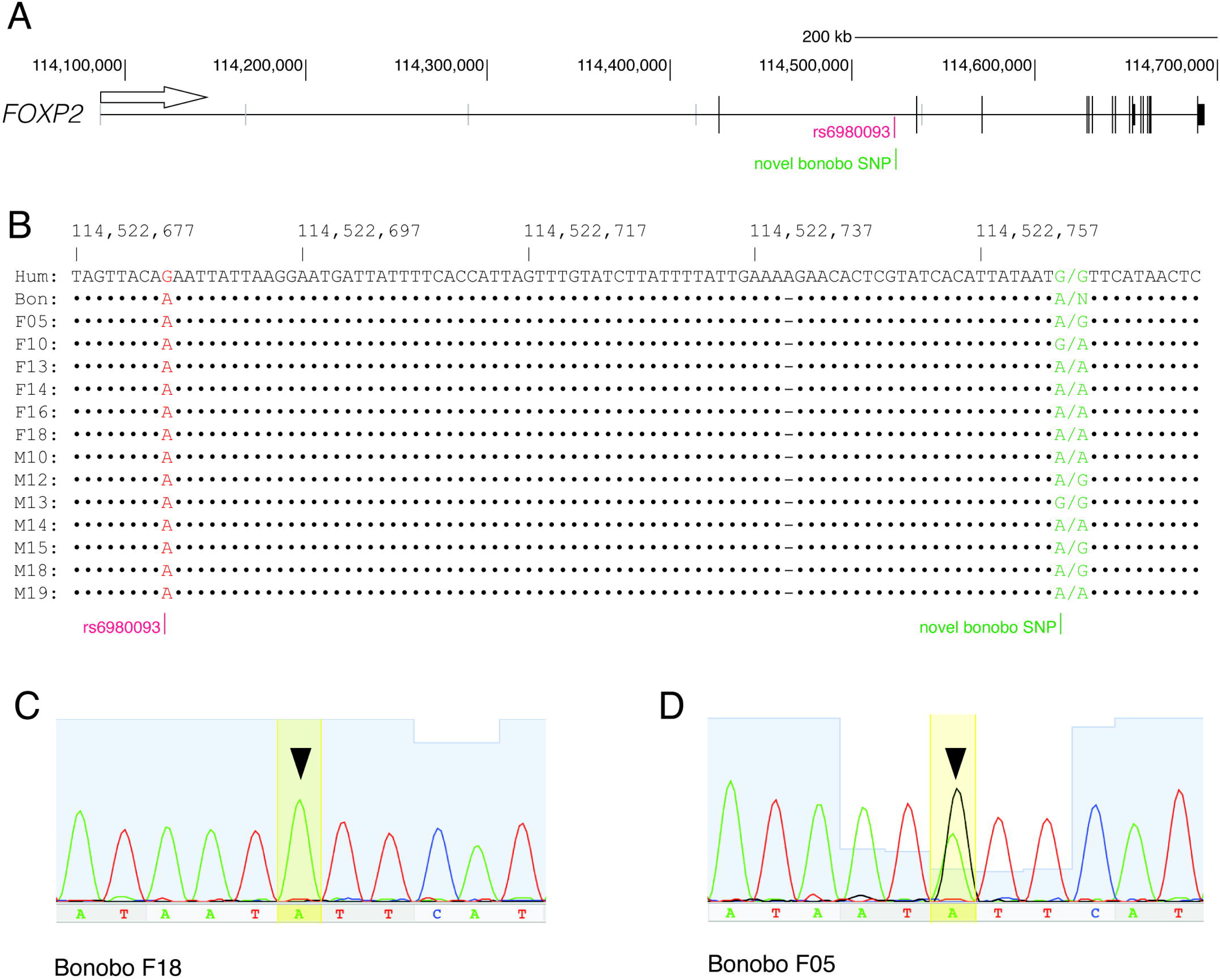
SNP in Bonobo *FOXP2*. Identification of a single-nucleotide polymorphism in bonobo *FOXP2* (A) Diagram of the human *FOXP2* gene showing relative location of the SNP rs6980093. Arrow shows direction of gene transcription. Short vertical bars (grey) show transcriptional start sites. Black vertical bars show exons. (B) Alignment of the human and bonobo references genomes, along with individual sequencing data from subjects in this study. Numbers refer to the human reference genome location on chromosome seven. (C, D) Representative Sanger sequencing chromatograms across the SNP, showing (C) a homozygous sample (F18), and (D) a heterozygous sample (F05).

Our analyses revealed genetic variation in bonobo *OXTR* at three novel positions (Figure 2A). Specifically, we identified a novel SNP 73bp upstream of rs237877 (chr3:8741201; C/T) and 184bp upstream of rs237878 (chr3:8741312; T/A/C); 11 of the 13 bonobos were homozygous (T/T) and 2 bonobos were heterozygous (T/G; 5’ flanking sequence TTGCAGCTATCACCTCATTT T/G; Novel *OXTR* SNP #1). We also identified a novel SNP 19bp downstream of human SNP rs35062132 (chr3:8753201; G/A/C). In our sample, 11 bonobos were homozygous (G/G), and 2 bonobos were heterozygous (G/A; 5’ flanking sequence CGATGGCTCAGGACAAAGGA G/A; Novel *OXTR* SNP #2). In addition, a novel SNP was observed 19bp upstream of rs237900 (chr3:8767010; G/A). Analyses revealed 3 genotypes at this locus; 8 bonobos were homozygous (A/A), 3 bonobos were heterozygous (A/C), and 2 bonobos were homozygous (C/C; 5’ flanking sequence GCCCAAGGACTGTGCTAAGG A/C; Novel *OXTR* SNP #3). Collectively, we observed a linked SNP combination (TGA) across the 3 novel bonobo *OXTR* sites at high frequency (65%) in the study population, including 6 homozygous bonobos (Figure 3). Observed genotypes across the novel *OXTR* sites included TGA (65%), TGC (27%), and GAA (8%). See Table 3 for a complete list of genotypes and Table 4 for the novel bonobo SNP frequencies.

**Figure 2.**
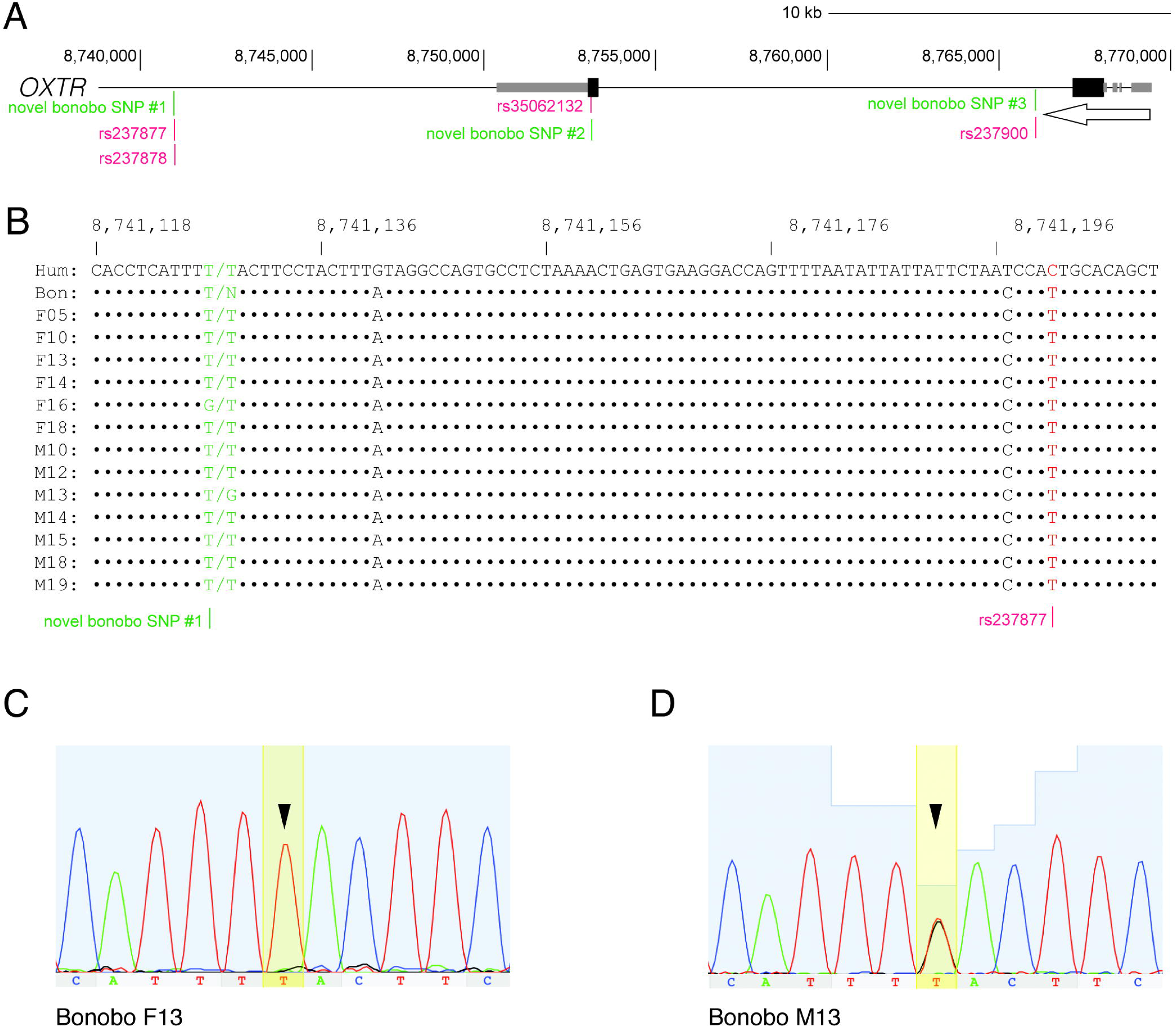
SNPs in Bonobo *OXTR*: Identification of single-nucleotide polymorphisms in bonobo *OXTR* (A) Diagram of the human *OXTR* gene showing relative locations of SNPs rs237877, rs237878, rs35062132, and rs237900, as well as the novel bonobo SNPs. Arrow shows direction of gene transcription. Untranslated regions are shown in grey. Black oblongs show exons. (B) Alignment of the human and bonobo references genomes, along with individual sequencing data from subjects in this study. Numbers refer to the human reference genome location on chromosome three. (C, D) Representative Sanger sequencing chromatograms across the novel SNP, showing (C) a homozygous sample (F13), and (D) a heterozygous sample (M13).

**Figure 3.**
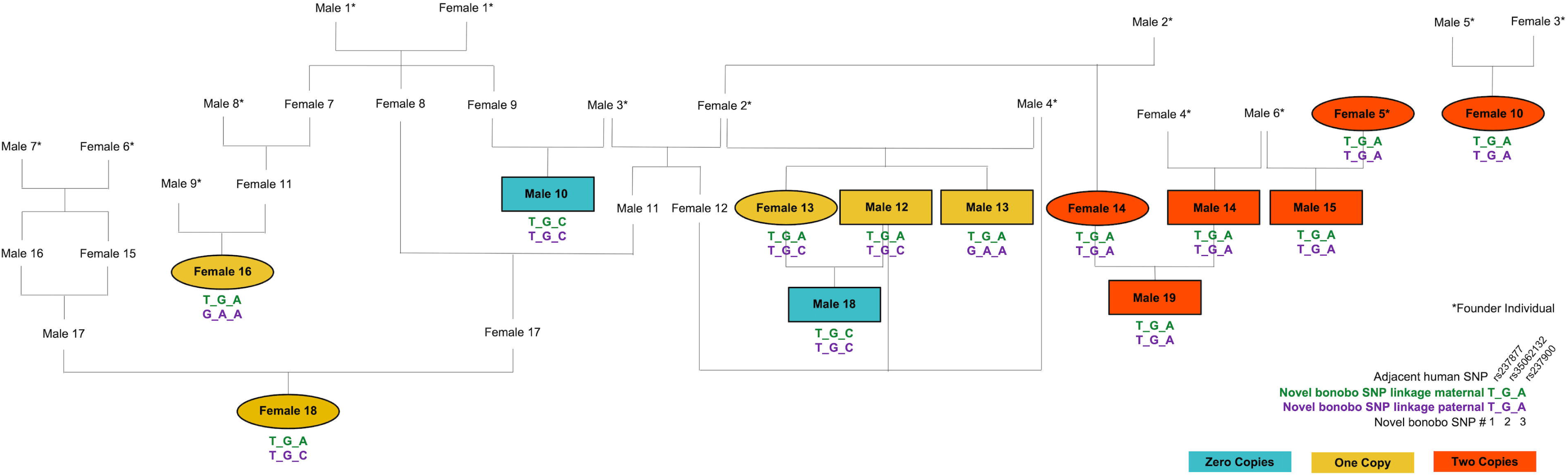
Bonobo Pedigree: Pedigree of the 13 subjects, with corresponding *OXTR* genotypes at the three novel SNPs in bonobo *OXTR*. Bonobos with zero copies are in blue, bonobos with one copy are in yellow, and bonobos with two copies of the TGA combination are in red.

**Table 3:** Individual Sequencing Data: Individual sequencing data for the novel SNPs identified in bonobo *FOXP2* and *OXTR* and for the selected human loci.

| Bonobo ID | F14 | F16 | M14 | F13 | M10 | F05 | M13 | F18 | M15 | M12 | M18 | F10 | M19 |
| --- | --- | --- | --- | --- | --- | --- | --- | --- | --- | --- | --- | --- | --- |
| Novel SNP *FOXP2* | A/A | A/A | A/A | A/A | A/A | A/G | G/G | A/A | A/G | A/G | A/G | A/G | A/A |
| Novel SNP *OXTR* #1 | T/T | T/G | T/T | T/T | T/T | T/T | T/G | T/T | T/T | T/T | T/T | T/T | T/T |
| Novel SNP *OXTR* #2 | G/G | G/A | G/G | G/G | G/G | G/G | G/A | G/G | G/G | G/G | G/G | G/G | G/G |
| Novel SNP *OXTR* #3 | A/A | A/A | A/A | A/C | C/C | A/A | A/A | A/C | A/A | A/C | C/C | A/A | A/A |
| *FOXP2* rs6980093 | A/A | A/A | A/A | A/A | A/A | A/A | A/A | A/A | A/A | A/A | A/A | A/A | A/A |
| *OXTR* rs2270463 | G/G | G/G | G/G | G/G | G/G | G/G | G/G | G/G | G/G | G/G | G/G | G/G | G/G |
| *OXTR* rs237877 | T/T | T/T | T/T | T/T | T/T | T/T | T/T | T/T | T/T | T/T | T/T | T/T | T/T |
| *OXTR* rs237878 | C/C | C/C | C/C | C/C | C/C | C/C | C/C | C/C | C/C | C/C | C/C | C/C | C/C |
| *OXTR* rs35062132 | G/G | G/G | G/G | G/G | G/G | G/G | G/G | G/G | G/G | G/G | G/G | G/G | G/G |
| *OXTR* rs2254295 | T/T | T/T | T/T | T/T | T/T | T/T | T/T | T/T | T/T | T/T | T/T | T/T | T/T |
| *OXTR* rs237894 | G/G | G/G | G/G | G/G | G/G | G/G | G/G | G/G | G/G | G/G | G/G | G/G | G/G |
| *OXTR* rs237895 | C/C | C/C | C/C | C/C | C/C | C/C | C/C | C/C | C/C | C/C | C/C | C/C | C/C |
| *OXTR* rs237900 | G/G | G/G | G/G | G/G | G/G | G/G | G/G | G/G | G/G | G/G | G/G | G/G | G/G |
| *AVPR1A* rs3803107 | G/G | G/G | G/G | G/G | G/G | G/G | G/G | G/G | G/G | G/G | G/G | G/G | G/G |

**Table 4:** Allele Frequencies in Bonobos:

| Novel Bonobo SNP | Nearest Human SNP(s) | Allele Frequency in Bonobos |
| --- | --- | --- |
| Novel SNP *FOXP2* | rs6980093 | A (.73) / G (.27) |
| Novel SNP *OXTR* #1 | rs237877, rs237878 | T (.92) / G (.08) |
| Novel SNP *OXTR* #2 | rs35062132 | G (.92) / A (.08) |
| Novel SNP *OXTR* #3 | rs237900 | A (.73) / C (.27) |

### 3.2 Behavioral differences

The behavioral analyses excluded one participant for whom behavioral data were not available – subject M15 (two copies of *OXTR* TGA and heterozygous at the novel *FOXP2* SNP). For *OXTR*, individual bonobos were grouped based on the number of TGA copies for the novel *OXTR* SNPs (Figure 3) – zero (n=2), one (n=5), or two (n=5; Table 5). Kruskal-Wallis test results indicated a significant *OXTR* group difference in social proximity score (*H* (2) = 15.461, *p* < 0.001; Figure 4; Table 6). Bonobos with two copies of TGA were more social (*Mdn* = 2.36) than individuals with one copy (*Mdn* = 1.77) and individuals with zero copies of TGA (*Mdn* = 1.55). A post-hoc Wilcoxon rank test with Benjamini-Hochberg adjustment revealed a significant difference in social proximity scores between bonobos with zero TGA copies and bonobos with two TGA copies (*p* < 0.001), and between bonobos with two copies and one copy (*p* = 0.007), but not between one copy and zero copies (*p* = 0.156). There was not a significant *OXTR* group difference in communicative production (*H* (2) = 2.888, *p* = 0.236; Figure 5, Table 7). For *FOXP2*, individuals were grouped based on whether they were homozygous (A/A, n=7), heterozygous (G/A, n=4), or homozygous (G/G, n=1) at the *FOXP2* SNP locus. Kruskal-Wallis test results indicated that there was not a significant *FOXP2* group difference in social proximity score (*H* (2) = 3.896, *p* = 0.143, Figure 6A), nor in communicative production (*H* (2) = 0.942, *p* = 0.625, Figure 6B). Post-hoc analyses indicated that rearing (atypical or typical) and age (subadult, adult, or elder) did not influence the observed differences in bonobo social behavior; there were no significant differences in social proximity score based on rearing (*W* = 1071, *p* = 0.717; Figure 7) or based on age (*H* (2) = 1.345, *p* = 0.510; Figure 8).

**Figure 4.**
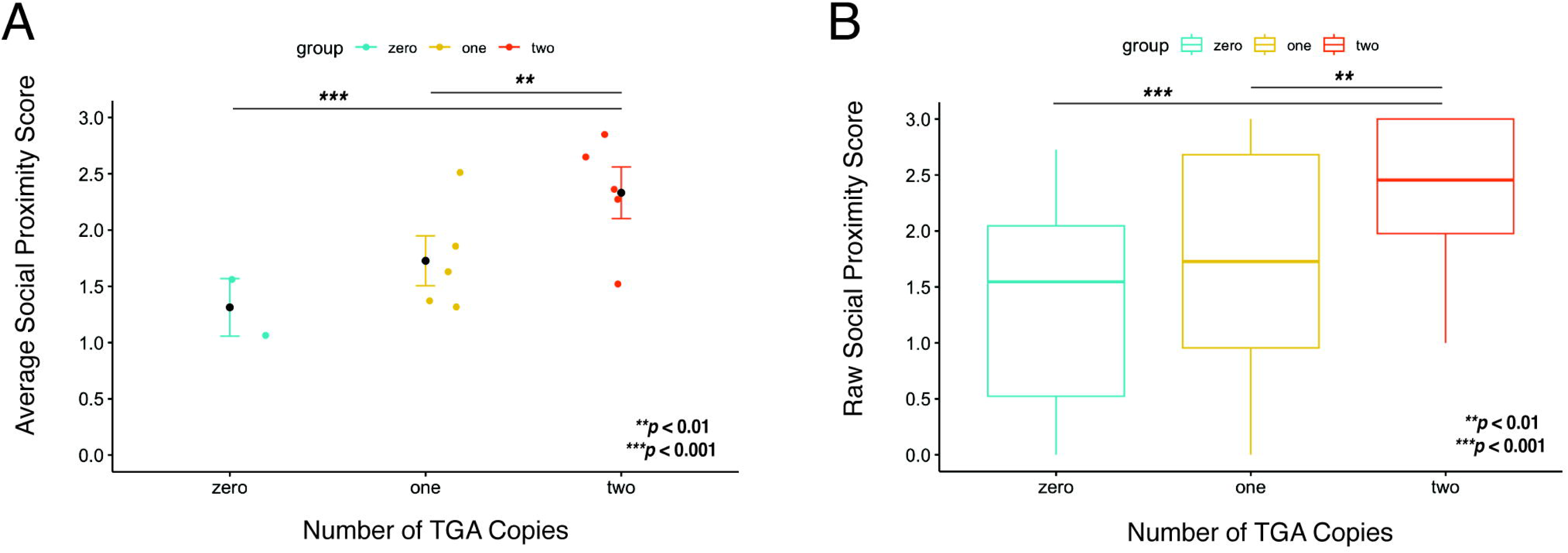
*OXTR* Group Differences in Bonobo Social Behavior: Observed group differences in social behavior based on the number of *OXTR* TGA copies. Average social proximity scores with standard error bars (A) and raw social proximity scores (B) for bonobos with zero, one, and two copies of the TGA combination.

**Figure 5.**
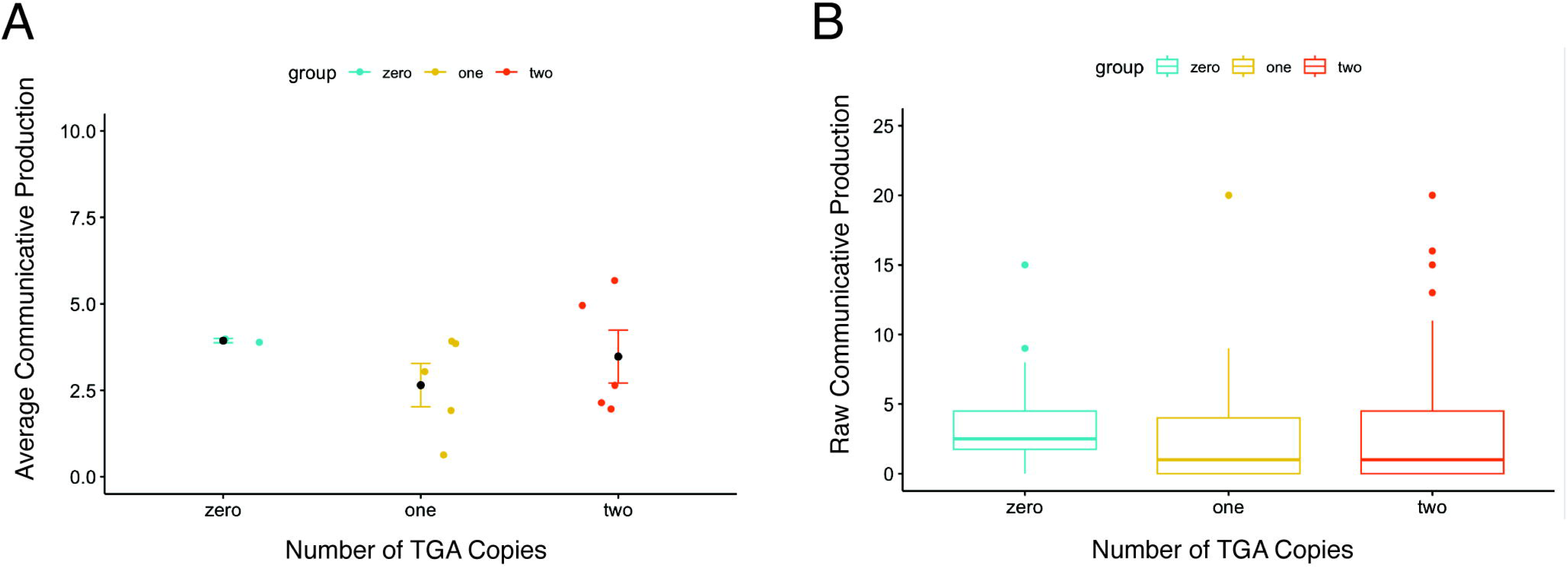
*OXTR* Groups and Bonobo Communicative Production: Communicative production based on the number of *OXTR* TGA copies. Average communicative production with standard error bars (A) and raw communicative production totals (B) for bonobos with zero, one, and two copies of the TGA combination.

**Figure 6.**
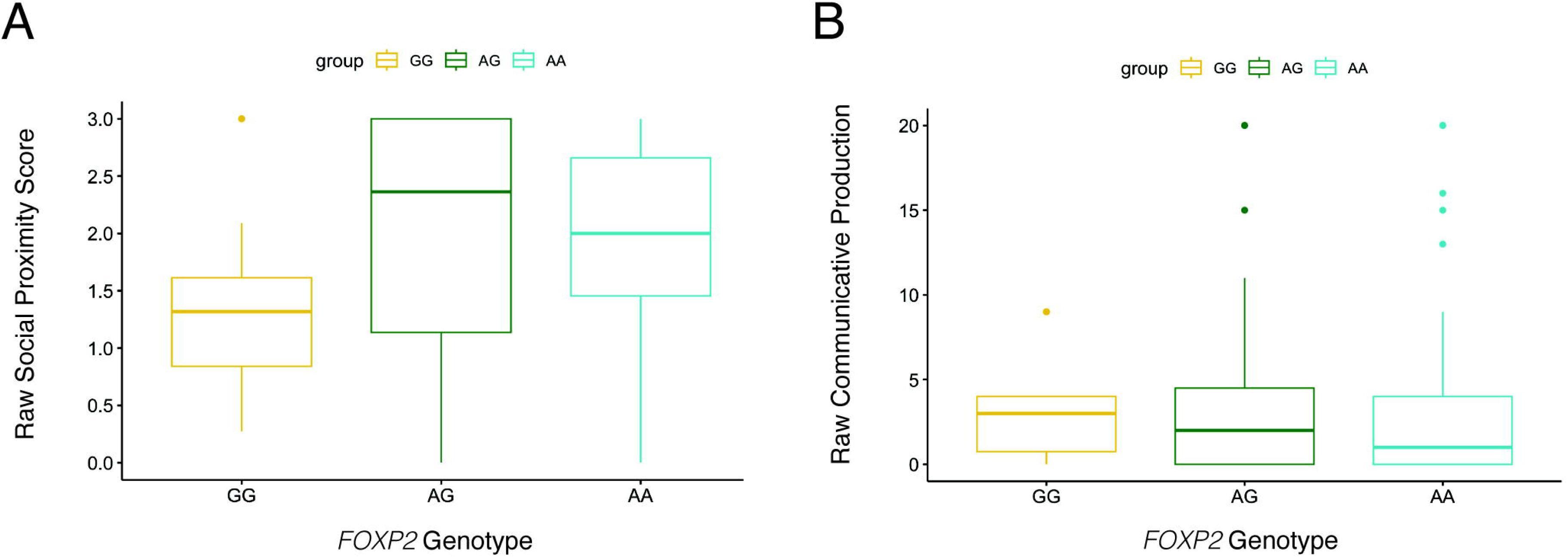
*FOXP2* Groups and Bonobo Social Communication: Social proximity and communicative production based on *FOXP2* genotype. Raw social proximity scores (A) and raw communicative production totals (B) for bonobos that were homozygous (G/G), heterozygous (A/G), and homozygous (A/A) at this position.

**Figure 7.**
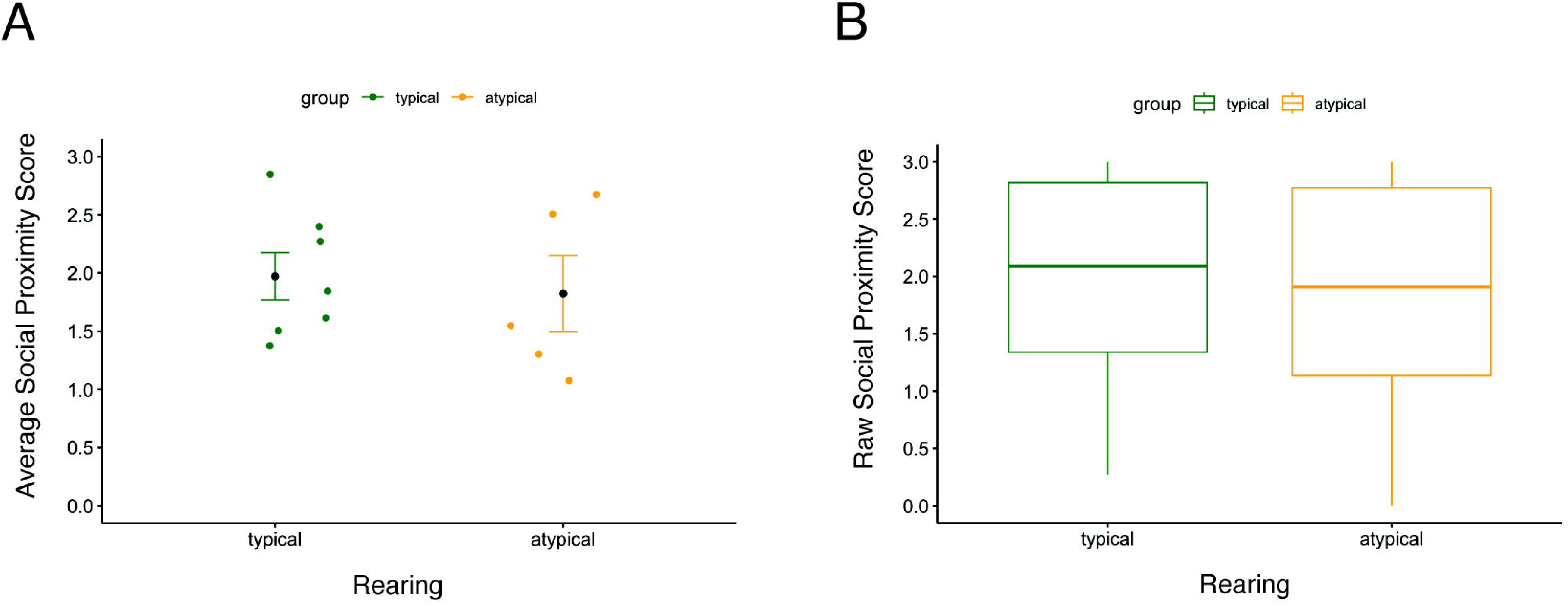
Rearing and Bonobo Social Behavior: Social proximity scores based on rearing. Average social proximity scores with standard error bars (A) and raw social proximity scores (B) for bonobos that were raised primarily by their mothers (typical) and bonobos that were raised primarily by humans or taken from the wild (atypical).

**Figure 8.**
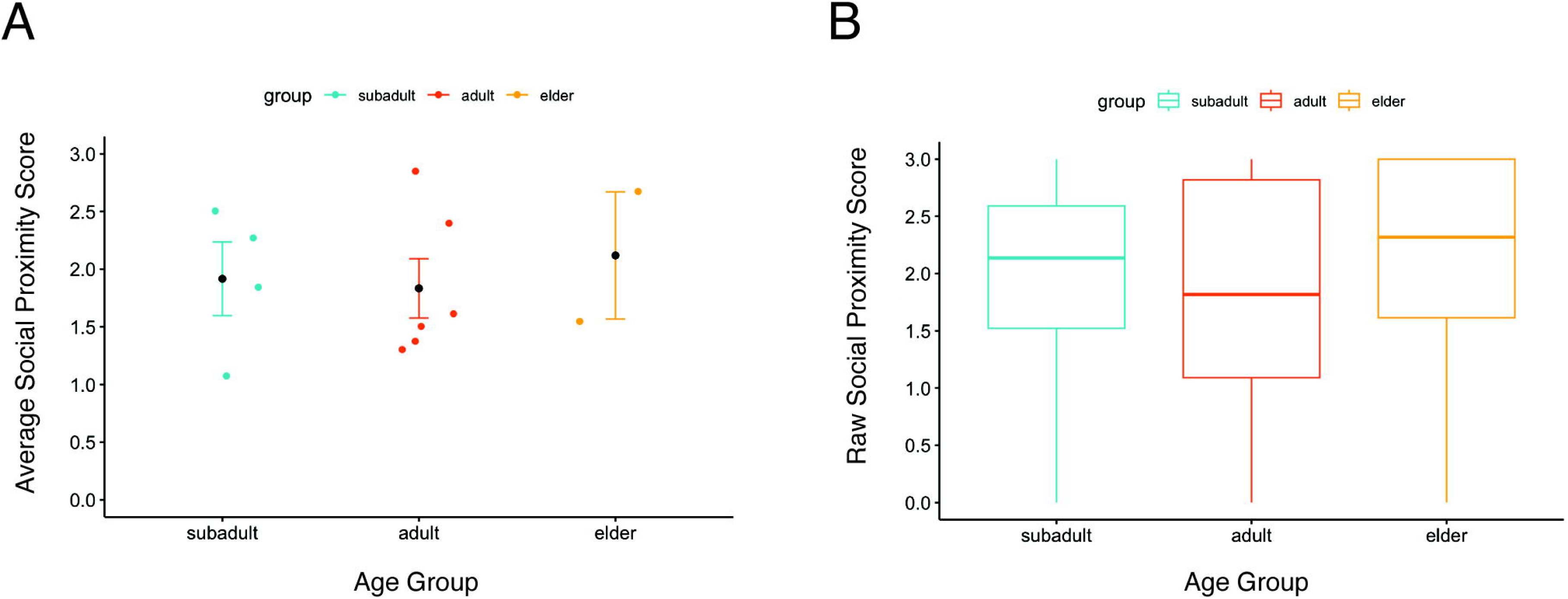
Age and Bonobo Social Behavior: Social proximity scores based on age. Average social proximity scores with standard error bars (A) and raw social proximity scores (B) for subadult (<14 years old), adult (14-34 years old), and elder (>34 years old) bonobos.

**Table 5:** *OXTR* Combined Genotypes: Individual identifications and corresponding ages for bonobos with zero, one, and two copies of the *OXTR* TGA combination.

| TGA Copies | Zero | One | Two |
| --- | --- | --- | --- |
| **Bonobo IDs** | M10, M18 | F13, F16, F18, M12, M13 | F05, F10, F14, M14, M19 |
| **Ages in Years** | 40, 10 | 23, 10, 13, 22, 20 | 35, 24, 24, 23, 6 |

**Table 6:** Social Proximity Scores: Social proximity scores for each of the bonobo’s focal observations.

| Bonobo ID | F14 | F16 | M14 | F13 | M10 | F05 | M13 | F18 | M12 | M18 | F10 | M19 |
| --- | --- | --- | --- | --- | --- | --- | --- | --- | --- | --- | --- | --- |
| Proximity Score 1 | 2.73 | 1.70 | 1.18 | 0.64 | 1.82 | 3.00 | 3.00 | 1.64 | 0.00 | 0.55 | 3.00 | 2.36 |
| Proximity Score 2 | 2.27 | 0.82 | 1.00 | 0.55 | 2.18 | 3.00 | 1.36 | 3.00 | 1.27 | 1.00 | 3.00 | 2.09 |
| Proximity Score 3 | 3.00 | 1.45 | 1.09 | 0.82 | 0.64 | 2.73 | 2.09 | 3.00 | 0.00 | 0.00 | 3.00 | 2.00 |
| Proximity Score 4 | 2.36 | 3.00 | 1.09 | 3.00 | 2.73 | 3.00 | 1.45 | 2.27 | 3.00 | 0.00 | 3.00 | 2.46 |
| Proximity Score 5 | 1.45 | 2.18 | 1.91 | 2.82 | 1.64 | 3.00 | 0.27 | 3.00 | 0.45 | 0.45 | 3.00 | 2.18 |
| Proximity Score 6 | 1.50 | 2.82 | 2.36 | 2.64 | 2.00 | 2.45 | 1.27 | 1.82 | 1.73 | 1.55 | 3.00 | 1.55 |
| Proximity Score 7 | 2.90 | 1.00 | 2.00 | 0.82 | 0.00 | 3.00 | 0.36 | 2.91 | 2.18 | 2.18 | 2.45 | 2.55 |
| Proximity Score 8 | 2.82 | 1.73 | 1.45 | 1.73 | 1.55 | 1.18 | 1.00 | 2.55 | 1.73 | 2.73 | 2.27 | 2.82 |

**Table 7:**
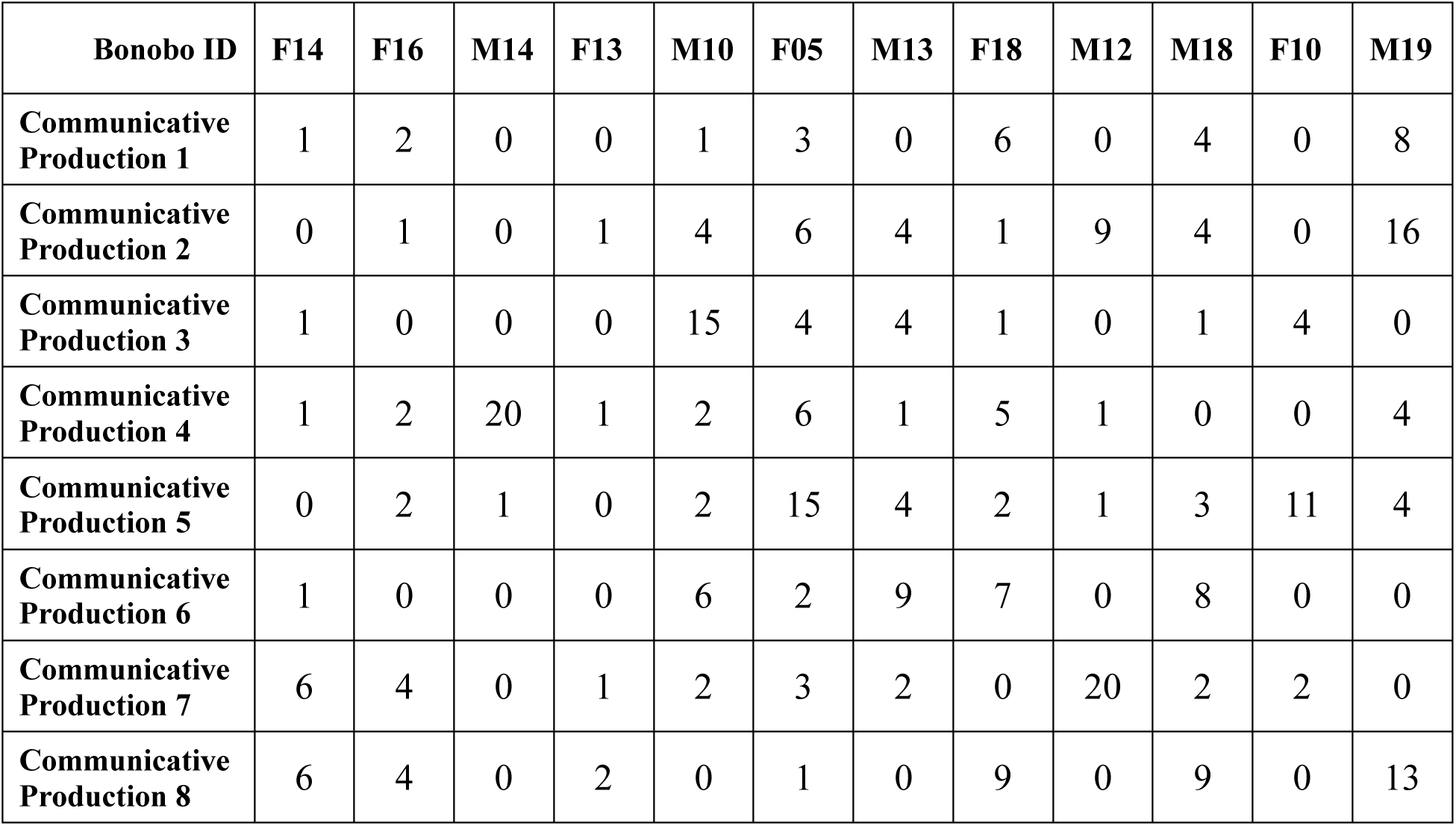
Communicative Production: Number of communicative signals produced during each of the bonobo’s focal observations.

## 4 Discussion

Determining the biological factors underlying individual-level social communication is key to understanding autism and other neurodevelopmental disorders and may aid in identifying children at risk of developing social communication impairments. Studying biobehavioral mechanisms in captive, nonhuman animals, permits researchers to elucidate the genetic contributions to complex behavioral phenotypes, while minimizing confounding rearing and environmental factors. Thus, we aimed to determine if one of the species most closely related to humans, the bonobo, exhibits SNPs in *FOXP2, OXTR*, and *AVPR1A* at known human loci.

Despite whole genomic investigations in all four nonhuman species of great ape, no SNPs have been identified in bonobo *FOXP2* to date. We are the first to report a SNP in bonobo *FOXP2* (near rs6980093) and we identified two novel genotypes in our sample. Although we did not find behavioral differences based on the bonobos’ *FOXP2* genotypes, we are encouraged by the identification of variation in bonobo *FOXP2* (75bp from the ASD-linked human SNP) and believe that further investigation into the bonobo *FOXP2* gene and measures of bonobo vocal communication might yield promising results.

The human *FOXP2* SNP rs6980093 is an intronic polymorphism, suggesting it may be involved in regulating *FOXP2* gene expression [7,69]. Several studies have identified relations between *FOXP2* rs6980093 variation and speech production [68], cortical activation in language-related regions [71], as well as speech and language learning abilities [7,72]. The identification of variation in bonobo *FOXP2* is a foundational step in understanding the impact of *FOXP2* on social communication in humans and our closest living relatives. Bonobos have the largest vocal repertoire size of the nonhuman great apes [73] and modify their communicative signals depending on social context [74–76]. In addition, multiple reports in a single bonobo support the notion that bonobos are able to understand spoken English words and sentences [77–79], as well as degraded and computer-generated speech [80]. Thus, it is possible that *FOXP2* alleles modulate *FOXP2* expression in bonobos and that differential *FOXP2* regulation impacts individual variability in bonobo vocal communication.

Specific *OXTR* variants have been linked to social functioning [81] and symptom severity [14,15] in autism. In our sample of bonobos, we identified a novel SNP between human *OXTR* rs237877 and rs237878. Variants at these positions have been linked to human levels of extraversion [8] and reward responsiveness [82] in typically developed adults, as well as neurological responses to oxytocin in autistic adults [83]. In addition, we identified a novel SNP in bonobo *OXTR* close to the human SNP rs35062132. Eleven of the 13 bonobos in our sample exhibited the G/G genotype. In humans, the C/G genotype of *OXTR* rs35062132 was associated with an increased risk of ASD and proposed to be a biological basis for individual differences in social behavior [84]. Although they identified three genotypes in their sample, Egawa and colleagues did not find a relation between these genetic variants and ASD [85]. Hence, further investigations are needed to understand the role rs35062132 variation plays in social communication development and to determine if the rs35062132 G allele is a risk factor for autism [84,85].

We also found a novel SNP in bonobo adjacent to rs237900. In humans, rs237900 variants were associated with reaction times for nonverbal-reaction-based-judgements, but not with nonverbal communication scores [83]. When controlling for sex, researchers found a significant rs237900 variant difference in harm avoidance. Female participants with the alternate (T) allele scored significantly higher on harm avoidance than all other participants [86]. Interestingly, rs237900 variants have also been linked to differences in DNA methylation of the *OXTR* gene [87]. Findings linking social functioning to another SNP in *OXTR,* rs2254298, are also well-established [88–91]. Yang and colleagues found differences in serum oxytocin levels based on the rs2254298 genotype [11]. However, the role that rs2254298 variants play in autism remains unclear. Specifically, results differ among human populations; the “A” allele was associated with autism in Japanese [92] and Chinese [93] populations, whereas the “G” allele was considered a risk factor in a population of autistic children and adolescents of European ancestry [94]. Parker and colleagues also identified links between rs2254298 variants and social impairments in children with and without ASD [95]. In both groups, individuals with the “A” allele exhibited greater social impairments than individuals without the “A” allele. Collectively, these findings suggest that specific *OXTR* variants might be promising biomarkers for social communication dysfunction in humans and highlight the need for alternative approaches to assessing the impact *OXTR* variants have on complex behavioral phenotypes, like those observed in ASD [95–97].

In addition to finding 3 novel *OXTR* sites, we found a linked SNP combination (TGA) across the *OXTR* sites at high frequency (65%) in the study population, including 6 homozygous bonobos. Linked *OXTR* variants have also been found in children, adolescents, and young adults diagnosed with ASD [12]. In addition, Wu and colleagues identified linkage among two *OXTR* variants in autistic people from the Chinese Han population [93]. Thus, we encourage researchers interested in the biomarkers of human social communication disorders to consider the relative influence of individual SNPs and their collective contribution to complex behavioral phenotypes.

If the observed TGA combination positively impacts bonobo sociality, we would expect to see behavioral differences based on these genotypes. In our sample, bonobos with two copies of the TGA combination had higher social proximity scores (i.e., they spent more time in close proximity to conspecific social partners) than bonobos with zero copies and bonobos with one copy of the TGA combination. This result supports previous conclusions that *OXTR* plays a pivotal role in bonobo social behavior [9,35,59] and is consistent with findings in humans that indicate that linked *OXTR* variants are associated with differences on the social responsiveness and repetitive behavior scales in autistic children [98]. In chimpanzees, researchers did not identify a link between *OXTR* variation and social behavior [99].

Our study complements several studies with rhesus macaques, including an investigation where an *OXTR* haplotype was associated with cognitive and behavioral differences [100]. Our results are also similar to data collected from children, adolescents, and young adults diagnosed with ASD; a haplotype comprised of four *OXTR* loci was associated with greater impairments in social interaction and communication in autistic individuals [12]. Thus, our findings suggest that the *OXTR* TGA combination may have been selected for in bonobos. It is also possible that the high frequency of the TGA combination is the product of a founder effect. Our collective results highlight the importance of considering multiple genetic variants in a given study and the benefits of multi-loci investigations in nonhuman great apes.

Given the documented relations between *OXTR* and social functioning, researchers have proposed that oxytocin can help facilitate social information processing in individuals with ASD. Indeed, evidence exists that oxytocin treatment can improve social abilities in children diagnosed with ASD [101] and that oxytocin infusions can reduce repetitive behaviors in autistic adults [83,102]. Researchers have also determined that oxytocin treatment efficacy differs between people, with the greatest improvements to social behavior occurring in individuals with the lowest pretreatment oxytocin blood concentration levels [101]. Along with previous evidence of a low social, high repetitive behavior phenotype in bonobos [50], our findings suggest that bonobos are an exemplary species for evaluating the efficacy of oxytocin interventions in the treatment of social communication dysfunction. Notwithstanding, further studies are needed to determine the specific characteristics that impact oxytocin efficacy and to identify biomarkers that predict which individuals will benefit the most from oxytocin treatments [82,101].

Unlike our findings in *FOXP2* and *OXTR*, we did not identify single-nucleotide variability in bonobo *AVPR1A*. With that said, we only targeted one human *AVPR1A* SNP (rs3803107) - sequencing ∼250 bp surrounding this locus. Other primer sets designed to amplify additional ASD-related SNPs in *AVPR1A* failed to work. Associations between *AVPR1A* and social behavior are well-documented in chimpanzees [57,99,103], and to a lesser extent bonobos [20]. However, the nonhuman great ape literature has centered around the RS3 microsatellite, given that chimpanzees are polymorphic for a secondary deletion of the DupB microsatellite-containing element in the 5’ flanking region of *AVPR1A* [17,20,103]. Many of these studies in chimpanzees and bonobos utilized personality questionnaire data, where human caregivers made a subjective estimate of each ape’s social behavior [20,56,99,103]. Additional objective assessments of individual-level social communication in nonhuman great apes are needed.

Seminal work in rodents accounts for much of what we know about *OXTR* and other ASD-related genes. However, many of these studies require substantial modification of the gene and/or the receptors or are limited in their translatability to human development and the complex phenotype of ASD [38,40,42]. In this study, we identified naturally occurring linkage among three novel *OXTR* variants and documented differences in bonobo social behavior based on this combined *OXTR* genotype. We also demonstrated that it is possible to detect genetic variability, variant linkage, and behavioral differences in even small samples of nonhuman great apes. If researchers incorporate our modified technique of rinsing the animal’s buccal cavity beforehand and swabbing the individual’s lower lip, they should be able to obtain voluntary biological samples that allow for sufficient DNA extraction from any nonhuman great ape living in human care. We encourage the inclusion of bonobos in future studies and suggest incorporating additional objective assessments of individual-level social communication (e.g., social attention, joint engagement, communicative proficiency). Singular investigations into Sanger sequencing data may underestimate individual variability across the target genes. Thus, we emphasize the need for a publicly accessible database to report SNPs identified in nonhuman primates and to contribute to larger scale genomic studies in nonhuman great apes with future research.

### 4.1 Conclusions

Here, we are the first to report a SNP in bonobo *FOXP2* – a gene necessary for typical linguistic development in humans. We also identified three novel SNPs in bonobo *OXTR* and demonstrated linkage among these *OXTR* variants. Our results indicate that individuals with two copies of the *OXTR* TGA combination are more social than individuals with one copy or zero copies of the TGA combination. Our collective findings suggest that these *OXTR* variants influence individual-level social communication in bonobos and support the notion that linked *OXTR* variants could be promising biological factors for identifying humans at risk of developing social communication deficits. This study also highlights the advantages of studying biobehavioral mechanisms in the species most closely related to humans and indicates that bonobos are a suitable model for testing hypotheses about the etiology of ASD and other human neurodevelopmental disorders.

## 7 Funding

This work was supported in part by NIH grant R25GM111565 (AB; PI Jonathan McMurry), Kennesaw State University Department of Ecology, Evolution, and Organismal Biology, and Autism Speaks Postdoctoral Fellowship 13439 (SAS). The funders had no role in study design, data collection and analysis, decision to publish, or preparation of the manuscript.

## 8 Acknowledgements

Thank you to Ape Initiative, the Columbus Zoo and Aquarium, and the Milwaukee County Zoo for providing biological samples, and to Dr. Hudson and Mr. Hansen for assisting in DNA extraction and purification. We would like to acknowledge that bonobos are an endangered species, with human activity as the greatest threat to their survival. Content from this manuscript appeared in a previous version on bioRxiv (doi: 10.1101/2023.12.22.573122).

## Notes

### Competing Interest Statement

The authors have declared no competing interest.

### Summary of Updates

The original manuscript version underwent peer-review at Frontiers in Behavioral Neuroscience. As a part of this process, a reviewer requested that the sequence files be included in the manuscript. While preparing these files, Dr. Hudson identified several errors in his work regarding the individual bonobo genotypes reported in the original version of the manuscript. Upon discovery of the errors, Dr. Skiba and Dr. Taglialatela conducted a new genetic analysis using the raw files and subsequently corrected the misreported information - updating the editorial team and including the raw sequences with the revised manuscript. In addition to requested edits for improvement and clarity by the peer-reviewers, this revised manuscript corrects several errors in the human and bonobo reference genomes, the SNP locations, and the individual genotypes reported in the original manuscript. Information in the figures, main text, and supplementary materials has been corrected in this revised version. Dr. Hudson and his lab member, Mr. Hansen, were unable to identify how the errors in data reporting occurred, and neither party contributed to the revised version of this manuscript. The co-author list has been adjusted to reflect contributions to the corrected analyses and revised manuscript. We are deeply grateful to the editorial team and reviewers at Frontiers in Behavioral Neuroscience for their thorough peer-review of this work - improving the overall quality of the manuscript and leading to the discovery and correction of substantial errors in the original version. We have also included this revision summary in the main manuscript file.

## References

1. Human Gene module. SFARI Gene. [Internet]. Available from: https://gene.sfari.org/database/human-gene/

2. Abrahams BS, Arking DE, Campbell DB, Mefford HC, Morrow EM, Weiss LA, et al. SFARI Gene 2.0: a community-driven knowledgebase for the autism spectrum disorders (ASDs). Molecular Autism. 2013 Dec;4(1):1–3.

3. Fisher SE, Scharff C. FOXP2 as a molecular window into speech and language. Trends in Genetics. 2009 Apr 1;25(4):166–77.

4. Lai CS, Fisher SE, Hurst JA, Vargha-Khadem F, Monaco AP. A forkhead-domain gene is mutated in a severe speech and language disorder. Nature. 2001 Oct 4;413(6855):519-23.

5. Shriberg LD, Ballard KJ, Tomblin JB, Duffy JR, Odell KH, Williams CA. Speech, prosody, and voice characteristics of a mother and daughter with a 7;13 translocation affecting *FOXP2*. J Speech Lang Hear Res. 2006;49(3):500–25.

6. Haghighatfard A, Yaghoubi asl E, Bahadori RA, Aliabadian R, Farhadi M, Mohammadpour F, et al. FOXP2 down expression is associated with executive dysfunctions and electrophysiological abnormalities of brain in Autism spectrum disorder; a neuroimaging genetic study. Autism & Developmental Language Impairments. 2022 Sep;7:23969415221126391.

7. Mozzi A, Riva V, Forni D, Sironi M, Marino C, Molteni M, et al. A common genetic variant in FOXP2 is associated with language-based learning (dis) abilities: Evidence from two Italian independent samples. American Journal of Medical Genetics Part B: Neuropsychiatric Genetics. 2017 Jul;174(5):578–86.

8. Haram M, Tesli M, Dieset I, Steen NE, Røssberg JI, Djurovic S, et al. An attempt to identify single nucleotide polymorphisms contributing to possible relationships between personality traits and oxytocin-related genes. Neuropsychobiology. 2014 Jan 22;69(1):25–30.

9. Theofanopoulou C, Andirkó A, Boeckx C, Jarvis ED. Oxytocin and vasotocin receptor variation and the evolution of human prosociality. Comprehensive Psychoneuroendocrinology. 2022 Aug 1;11:100139.

10. Wu N, Su Y. Variations in the oxytocin receptor gene and prosocial behavior: moderating effects of situational factors. Integrative Zoology. 2018 Nov;13(6):687–97.

11. Yang S, Dong X, Guo X, Han Y, Song H, Gao L, et al. Serum oxytocin levels and an oxytocin receptor gene polymorphism (rs2254298) indicate social deficits in children and adolescents with autism spectrum disorders. Frontiers in Neuroscience. 2017 Apr 21;11:221.

12. Wermter AK, Kamp-Becker I, Hesse P, Schulte-Körne G, Strauch K, Remschmidt H. Evidence for the involvement of genetic variation in the oxytocin receptor gene (OXTR) in the etiology of autistic disorders on high-functioning level. American Journal of Medical Genetics Part B: Neuropsychiatric Genetics. 2010 Mar;153(2):629–39.

13. Holmqvist Jämsen S, Johansson A, Westberg L, Santtila P, von der Pahlen B, Simberg S. Associations between vocal symptoms and genetic variants in the oxytocin receptor and arginine vasopressin 1A receptor gene. Journal of Speech, Language, and Hearing Research. 2017 Jul 12;60(7):1843–54.

14. Oztan O, Jackson LP, Libove RA, Sumiyoshi RD, Phillips JM, Garner JP, et al. Biomarker discovery for disease status and symptom severity in children with autism. Psychoneuroendocrinology. 2018 Mar 1;89:39–45.

15. Cataldo I, Azhari A, Esposito G. A review of oxytocin and arginine-vasopressin receptors and their modulation of autism spectrum disorder. Frontiers in Molecular Neuroscience. 2018 Feb 13;11:27.

16. Bachner-Melman R, Zohar AH, Bacon-Shnoor N, Elizur Y, Nemanov L, Gritsenko I, et al. Link between vasopressin receptor AVPR1A promoter region microsatellites and measures of social behavior in humans. Journal of Individual Differences. 2005 Jan;26(1):2–10.

17. Donaldson ZR, Kondrashov FA, Putnam A, Bai Y, Stoinski TL, Hammock EA, Young LJ. Evolution of a behavior-linked microsatellite-containing element in the 5’ flanking region of the primate AVPR1A gene. BMC Evol Biol. 2008 Jun 23;8:180.

18. Johnson ZV, Young LJ. Oxytocin and vasopressin neural networks: Implications for social behavioral diversity and translational neuroscience. Neuroscience & Biobehavioral Reviews. 2017 May 1;76:87–98.

19. Mahovetz LM, Young LJ, Hopkins WD. The influence of AVPR1A genotype on individual differences in behaviors during a mirror self-recognition task in chimpanzees (Pan troglodytes). Genes, Brain and Behavior. 2016 Jun;15(5):445–52.

20. Staes N, Weiss A, Helsen P, Korody M, Eens M, Stevens JM. Bonobo personality traits are heritable and associated with vasopressin receptor gene 1a variation. Scientific Reports. 2016 Dec 2;6(1):38193.

21. Staes N, Bradley BJ, Hopkins WD, Sherwood CC. Genetic signatures of socio-communicative abilities in primates. Current Opinion in Behavioral Sciences. 2018 Jun 1;21:33–8.

22. Wang J, Qin W, Liu F, Liu B, Zhou Y, Jiang T, et al. Sex-specific mediation effect of the right fusiform face area volume on the association between variants in repeat length of AVPR 1 A RS 3 and altruistic behavior in healthy adults. Human brain mapping. 2016 Jul;37(7):2700–9.

23. Hopkins WD, Keebaugh AC, Reamer LA, Schaeffer J, Schapiro SJ, Young LJ. Genetic influences on receptive joint attention in chimpanzees (Pan troglodytes). Scientific reports. 2014 Jan 20;4(1):3774.

24. Wade M, Hoffmann TJ, Jenkins JM. Association between the arginine vasopressin receptor 1A (AVPR1A) gene and preschoolers’ executive functioning. Brain and Cognition. 2014 Oct 1;90:116–23.

25. Walum H, Westberg L, Henningsson S, Neiderhiser JM, Reiss D, Igl W, et al. Genetic variation in the vasopressin receptor 1a gene (AVPR1A) associates with pair-bonding behavior in humans. Proceedings of the National Academy of Sciences. 2008 Sep 16;105(37):14153–6.

26. Knafo A, Israel S, Darvasi A, Bachner-Melman R, Uzefovsky F, Cohen L, et al. Individual differences in allocation of funds in the dictator game associated with length of the arginine vasopressin 1a receptor RS3 promoter region and correlation between RS3 length and hippocampal mRNA. Genes, brain and behavior. 2008 Apr;7(3):266–75.

27. Mulholland MM, Navabpour SV, Mareno MC, Schapiro SJ, Young LJ, Hopkins WD. AVPR1A variation is linked to gray matter covariation in the social brain network of chimpanzees. Genes, Brain and Behavior. 2020 Apr;19(4):e12631.

28. De Abreu MS, Genario R, Giacomini AC, Demin KA, Lakstygal AM, Amstislavskaya TG, et al. Zebrafish as a model of neurodevelopmental disorders. Neuroscience. 2020 Oct 1;445:3–11.

29. Crawley JN. Translational animal models of autism and neurodevelopmental disorders. Dialogues in Clinical Neuroscience. 2022 Apr 1.

30. Eaton SL, Wishart TM. Bridging the gap: large animal models in neurodegenerative research. Mammalian Genome. 2017 Aug;28:324–37.

31. Chabout J, Sarkar A, Patel SR, Radden T, Dunson DB, Fisher SE, et al. A Foxp2 mutation implicated in human speech deficits alters sequencing of ultrasonic vocalizations in adult male mice. Frontiers in Behavioral Neuroscience. 2016 Oct 20;10:197.

32. Medvedeva VP, Rieger MA, Vieth B, Mombereau C, Ziegenhain C, Ghosh T, et al. Altered social behavior in mice carrying a cortical Foxp2 deletion. Human Molecular Genetics. 2019 Mar 1;28(5):701–17.

33. Gemmer A, Mirkes K, Anneser L, Eilers T, Kibat C, Mathuru A, et al. Oxytocin receptors influence the development and maintenance of social behavior in zebrafish (*Danio rerio*). Scientific Reports. 2022 Mar 12;12(1):4322.

34. Staes N, Sherwood CC, Wright K, De Manuel M, Guevara EE, Marques-Bonet T, et al. FOXP2 variation in great ape populations offers insight into the evolution of communication skills. Scientific Reports. 2017 Dec 4;7(1):16866.

35. Staes N, Stevens JM, Helsen P, Hillyer M, Korody M, Eens M. Oxytocin and vasopressin receptor gene variation as a proximate base for inter-and intraspecific behavioral differences in bonobos and chimpanzees. PLoS One. 2014 Nov 18;9(11):e113364.

36. Heston JB, White SA. Behavior-linked FoxP2 regulation enables zebra finch vocal learning. Journal of Neuroscience. 2015 Feb 18;35(7):2885–94.

37. Donaldson ZR, Young LJ. The relative contribution of proximal 5′ flanking sequence and microsatellite variation on brain vasopressin 1a receptor (Avpr1a) gene expression and behavior. PLoS Genetics. 2013 Aug 29;9(8):e1003729.

38. Fischer J, Hammerschmidt K. Ultrasonic vocalizations in mouse models for speech and socio-cognitive disorders: insights into the evolution of vocal communication. Genes, Brain and Behavior. 2011 Feb;10(1):17–27.

39. Johnson ZV, Walum H, Xiao Y, Riefkohl PC, Young LJ. Oxytocin receptors modulate a social salience neural network in male prairie voles. Hormones and Behavior. 2017 Jan 1;87:16–24.

40. Pobbe RL, Pearson BL, Blanchard DC, Blanchard RJ. Oxytocin receptor and Mecp2308/Y knockout mice exhibit altered expression of autism-related social behaviors. Physiology & Behavior. 2012 Dec 5;107(5):641–8.

41. Shu W, Cho JY, Jiang Y, Zhang M, Weisz D, Elder GA, et al. Altered ultrasonic vocalization in mice with a disruption in the Foxp2 gene. Proceedings of the National Academy of Sciences. 2005 Jul 5;102(27):9643–8.

42. Bey AL, Jiang YH. Overview of mouse models of autism spectrum disorders. Current Protocols in Pharmacology. 2014 Sep;66(1):5–66.

43. Silverman JL, Yang M, Lord C, Crawley JN. Behavioural phenotyping assays for mouse models of autism. Nature Reviews Neuroscience. 2010 Jul;11(7):490–502.

44. Berendzen KM, Sharma R, Mandujano MA, Wei Y, Rogers FD, Simmons TC, et al. Oxytocin receptor is not required for social attachment in prairie voles. Neuron. 2023 Mar 15;111(6):787–96.

45. Prüfer K, Munch K, Hellmann I, Akagi K, Miller JR, Walenz B, Koren S, Sutton G, Kodira C, Winer R, Knight JR. The bonobo genome compared with the chimpanzee and human genomes. Nature. 2012 Jun 28;486(7404):527-31.

46. Clay Z, Genty E. Natural communication in bonobos: insights into social awareness and the evolution of language. Bonobos. Unique in Mind, Brain and Behaviour. Oxford University Press. 2017:105–22.

47. Clay Z, Ravaux L, de Waal F, Zuberbühler K. Bonobos (Pan paniscus) vocally protest against violations of social expectations. Journal of Comparative Psychology. 2016 Feb;130(1):44.

48. Mulholland MM, Mahovetz LM, Mareno MC, Reamer LA, Schapiro SJ, Hopkins WD. Differences in the mutual eye gaze of bonobos (Pan paniscus) and chimpanzees (Pan troglodytes). Journal of Comparative Psychology. 2020 Aug;134(3):318.

49. Surbeck M, Hohmann G. Social preferences influence the short-term exchange of social grooming among male bonobos. Animal Cognition. 2015 Mar;18:573–9.

50. Skiba, SA. Behavioral, cognitive, and genetic factors underlying socio-communicative development in bonobos. PhD. Dissertation, Georgia State University; 2021 Jul 23. Available from: https://scholarworks.gsu.edu/psych_diss/237/

51. Pika S, Liebal K, Tomasello M. Gestural communication in subadult bonobos (Pan paniscus): repertoire and use. American Journal of Primatology: Official Journal of the American Society of Primatologists. 2005 Jan;65(1):39–61.

52. Schamberg I, Cheney DL, Clay Z, Hohmann G, Seyfarth RM. Call combinations, vocal exchanges and interparty movement in wild bonobos. Animal Behaviour. 2016 Dec 1;122:109–16.

53. Laméris DW, Staes N, Salas M, Matthyssen S, Verspeek J, Stevens JM. The influence of sex, rearing history, and personality on abnormal behaviour in zoo-housed bonobos (Pan paniscus). Applied Animal Behaviour Science. 2021 Jan 1;234:105178.

54. Kovalaskas S, Rilling JK, Lindo J. Comparative analyses of the Pan lineage reveal selection on gene pathways associated with diet and sociality in bonobos. Genes, Brain and Behavior. 2021 Mar;20(3):e12715.

55. Staes N, Guevara EE, Helsen P, Eens M, Stevens JM. The Pan social brain: An evolutionary history of neurochemical receptor genes and their potential impact on sociocognitive differences. Journal of Human Evolution. 2021 Mar 1;152:102949.

56. Anestis SF, Webster TH, Kamilar JM, Fontenot MB, Watts DP, Bradley BJ. AVPR1A variation in chimpanzees (Pan troglodytes): Population differences and association with behavioral style. International Journal of Primatology. 2014 Feb;35:305–24.

57. Hopkins WD, Staes N, Guevara EE, Mulholland MM, Sherwood CC, Bradley BJ. Vasopressin, and not oxytocin, receptor gene methylation is associated with individual differences in receptive joint attention in chimpanzees (Pan troglodytes). Autism Research. 2023 Apr;16(4):713–22.

58. Schneider E, El Hajj N, Richter S, Roche-Santiago J, Nanda I, Schempp W, Riederer P, et al. Widespread differences in cortex DNA methylation of the “language gene” CNTNAP2 between humans and chimpanzees. Epigenetics. 2014 Apr 17;9(4):533–45.

59. Moscovice LR, Surbeck M, Fruth B, Hohmann G, Jaeggi AV, Deschner T. The cooperative sex: sexual interactions among female bonobos are linked to increases in oxytocin, proximity and coalitions. Hormones and Behavior. 2019 Nov 1;116:104581.

60. Primer designing tool. Nih.gov. [Internet]. Available from: https://www.ncbi.nlm.nih.gov/tools/primer-blast/

61. ApE-A plasmid Editor. Utah.edu. [Internet]. Available from: https://jorgensen.biology.utah.edu/wayned/ape/

62. UCSC Genome Browser. Genome.ucsc.edu [Internet]. Available from: https://genome.ucsc.edu/index.html

63. Madeira F, Madhusoodanan N, Lee J, et al. The EMBL-EBI Job Dispatcher sequence analysis tools framework in 2024. Nucleic Acids Research. 2024 Jul;52(W1):W521–W525.

64. 4Peaks. Nucleobytes.com. [Internet]. Available from: https://nucleobytes.com/4peaks/index.html

65. Skiba SA. The adaptive value of complex socio-communicative behavior. M.S. Thesis. Kennesaw State University; 2017 Jul 14. Available from: https://digitalcommons.kennesaw.edu/integrbiol_etd/23/

66. Furuichi T. Social interactions and the life history of female Pan paniscus in Wamba, Zaire. International journal of Primatology. 1989 Jun;10:173–97.

67. Gelardi V, Godard J, Paleressompoulle D, Claidiere N, Barrat A. Measuring social networks in primates: wearable sensors versus direct observations. Proceedings of the Royal Society A. 2020 Apr 29;476(2236):20190737.

68. Taglialatela JP, Skiba SA, Evans RE, Bogart S, Schwob NG. Social behavior and social tolerance in chimpanzees and bonobos. Chimpanzee in Context: A Comparative Perspective on Chimpanzee Behavior, Cognition, Conservation, and Welfare. 2020:95–114.

69. Cooper DN. Functional intronic polymorphisms: Buried treasure awaiting discovery within our genes. Human Genomics. 2010 Dec;4(5):1–5.

70. Zhang S, Zhao J, Guo Z, Jones JA, Liu P, Liu H. The association between genetic variation in FOXP2 and sensorimotor control of speech production. Frontiers in Neuroscience. 2018 Sep 20;12:666.

71. Pinel P, Fauchereau F, Moreno A, Barbot A, Lathrop M, Zelenika D, et al. Genetic variants of FOXP2 and KIAA0319/TTRAP/THEM2 locus are associated with altered brain activation in distinct language-related regions. Journal of Neuroscience. 2012 Jan 18;32(3):817–25.

72. Chandrasekaran B, Yi HG, Blanco NJ, McGeary JE, Maddox WT. Enhanced procedural learning of speech sound categories in a genetic variant of FOXP2. Journal of Neuroscience. 2015 May 20;35(20):7808–12.

73. McComb K, Semple S. Coevolution of vocal communication and sociality in primates. Biology Letters. 2005 Dec 22;1(4):381–5.

74. Clay Z, Archbold J, Zuberbühler K. Functional flexibility in wild bonobo vocal behaviour. PeerJ. 2015 Aug 4;3:e1124.

75. Clay Z, Zuberbühler K. Communication during sex among female bonobos: effects of dominance, solicitation and audience. Scientific Reports. 2012 Mar 1;2(1):291.

76. Hohmann G, Fruth B. Structure and use of distance calls in wild bonobos (Pan paniscus). International Journal of Primatology. 1994 Oct;15:767–82.

77. Margiotoudi K, Bohn M, Schwob N, Taglialatela J, Pulvermüller F, Epping A, et al. Bo-NO-bouba-kiki: picture-word mapping but no spontaneous sound symbolic speech-shape mapping in a language trained bonobo. Proceedings of the Royal Society B. 2022 Feb 9;289(1968):20211717.

78. Brakke KE, Savage-Rumbaugh ES. The development of language skills in bonobo and chimpanzee: I. Comprehension. Language & Communication. 1995 Apr.

79. Savage-Rumbaugh ES, Murphy J, Sevcik RA, Brakke KE, Williams SL, Rumbaugh DM, et al. Language comprehension in ape and child. Monographs of the Society for Research in Child Development. 1993 Jan 1:i-252.

80. Lahiff NJ, Slocombe KE, Taglialatela J, Dellwo V, Townsend SW. Degraded and computer-generated speech processing in a bonobo. Animal Cognition. 2022 Dec;25(6):1393–8.

81. Baribeau DA, Dupuis A, Paton TA, Scherer SW, Schachar RJ, Arnold PD, et al. Oxytocin receptor polymorphisms are differentially associated with social abilities across neurodevelopmental disorders. Scientific Reports. 2017 Sep 14;7(1):11618.

82. Davis C, Zai CC, Adams N, Bonder R, Kennedy JL. Oxytocin and its association with reward-based personality traits: A multilocus genetic profile (MLGP) approach. Personality and Individual Differences. 2019 Feb 1;138:231–6.

83. Watanabe T, Otowa T, Abe O, Kuwabara H, Aoki Y, Natsubori T, et al. Oxytocin receptor gene variations predict neural and behavioral response to oxytocin in autism. Social Cognitive and Affective Neuroscience. 2017 Mar;12(3):496–506.

84. Ma WJ, Hashii M, Munesue T, Hayashi K, Yagi K, Yamagishi M. Non-synonymous single-nucleotide variations of the human oxytocin receptor gene and autism spectrum disorders: a case–control study in a Japanese population and functional analysis. Molecular Autism. 2013 Dec;4(1):1–4.

85. Egawa J, Watanabe Y, Shibuya M, Endo T, Sugimoto A, Igeta H, et al. Resequencing and association analysis of OXTR with autism spectrum disorder in a Japanese population. Psychiatry and Clinical Neurosciences. 2015 Mar;69(3):131–5.

86. Stankova T, Eichhammer P, Langguth B, Sand PG. Sexually dimorphic effects of oxytocin receptor gene (OXTR) variants on Harm Avoidance. Biology of sex differences. 2012 Dec;3:1–5.

87. Milaniak I, Cecil CA, Barker ED, Relton CL, Gaunt TR, McArdle W, Jaffee SR. Variation in DNA methylation of the oxytocin receptor gene predicts children’s resilience to prenatal stress. Development and Psychopathology. 2017 Dec;29(5):1663–74.

88. Bozorgmehr A, Alizadeh F, Sadeghi B, Shahbazi A, Ofogh SN, Joghataei MT, et al. Oxytocin moderates risky decision-making during the Iowa gambling task: A new insight based on the role of oxytocin receptor gene polymorphisms and interventional cognitive study. Neuroscience Letters. 2019 Aug 24;708:134328.

89. Costa B, Pini S, Gabelloni P, Abelli M, Lari L, Cardini A, et al. Oxytocin receptor polymorphisms and adult attachment style in patients with depression. Psychoneuroendocrinology. 2009 Nov 1;34(10):1506–14.

90. Thompson RJ, Parker KJ, Hallmayer JF, Waugh CE, Gotlib IH. Oxytocin receptor gene polymorphism (rs2254298) interacts with familial risk for psychopathology to predict symptoms of depression and anxiety in adolescent girls. Psychoneuroendocrinology. 2011 Jan 1;36(1):144–7.

91. Wu N, Li Z, Su Y. The association between oxytocin receptor gene polymorphism (OXTR) and trait empathy. Journal of Affective Disorders. 2012 May 1;138(3):468–72.

92. Liu X, Kawamura Y, Shimada T, Otowa T, Koishi S, Sugiyama T. Association of the oxytocin receptor (OXTR) gene polymorphisms with autism spectrum disorder (ASD) in the Japanese population. Journal of Human Genetics. 2010 Mar;55(3):137–41.

93. Wu S, Jia M, Ruan Y, Liu J, Guo Y, Shuang M, et al. Positive association of the oxytocin receptor gene (OXTR) with autism in the Chinese Han population. Biological Psychiatry. 2005 Jul 1;58(1):74–7.

94. Jacob S, Brune CW, Carter CS, Leventhal BL, Lord C, Cook Jr EH. Association of the oxytocin receptor gene (OXTR) in Caucasian children and adolescents with autism. Neuroscience letters. 2007 Apr 24;417(1):6–9.

95. Parker KJ, Garner JP, Libove RA, Hyde SA, Hornbeak KB, Carson DS, et al. Plasma oxytocin concentrations and OXTR polymorphisms predict social impairments in children with and without autism spectrum disorder. Proceedings of the National Academy of Sciences. 2014 Aug 19;111(33):12258–63.

96. LoParo D, Waldman ID. The oxytocin receptor gene (OXTR) is associated with autism spectrum disorder: a meta-analysis. Molecular Psychiatry. 2015 May;20(5):640–6.

97. Ylisaukko-oja T, Alarcón M, Cantor RM, Auranen M, Vanhala R, Kempas E, et al. Search for autism loci by combined analysis of Autism Genetic Resource Exchange and Finnish families. Annals of Neurology. 2006 Jan;59(1):145–55.

98. Harrison AJ, Gamsiz ED, Berkowitz IC, Nagpal S, Jerskey BA. Genetic variation in the oxytocin receptor gene is associated with a social phenotype in autism spectrum disorders. American Journal of Medical Genetics Part B: Neuropsychiatric Genetics. 2015 Dec;168(8):720–9.

99. Staes N, Koski SE, Helsen P, Fransen E, Eens M, Stevens JM. Chimpanzee sociability is associated with vasopressin (Avpr1a) but not oxytocin receptor gene (OXTR) variation. Hormones and Behavior. 2015 Sep 1;75:84–90.

100. Howarth ER, Szott ID, Witham CL, Wilding CS, Bethell EJ. Genetic polymorphisms in the serotonin, dopamine and opioid pathways influence social attention in rhesus macaques (Macaca mulatta). PLoS One. 2023 Aug 2;18(8):e0288108.

101. Parker KJ, Oztan O, Libove RA, Sumiyoshi RD, Jackson LP, Karhson DS, et al. Intranasal oxytocin treatment for social deficits and biomarkers of response in children with autism. Proceedings of the National Academy of Sciences. 2017 Jul 25;114(30):8119–24.

102. Hollander E, Novotny S, Hanratty M, Yaffe R, DeCaria CM, Aronowitz BR, et al. Oxytocin infusion reduces repetitive behaviors in adults with autistic and Asperger’s disorders. Neuropsychopharmacology. 2003 Jan;28(1):193–8.

103. Wilson VA, Weiss A, Humle T, Morimura N, Udono T, Idani GI, et al. Chimpanzee personality and the arginine vasopressin receptor 1A genotype. Behavior Genetics. 2017 Mar;47:215–26

