## Supplementary Material 1 for "Linked *OXTR* Variants Are Associated With Social Behavior Differences in Bonobos (*Pan paniscus*)"

**Supplementary Material 1 – Excluded Human SNPs**

Primer pairs that were included in the study met the following criteria: 1) the PCR product was at least 250 base pairs long, 2) the melting temperature (Tm) of the primer was 60°C and the GC% was approximately 50%, and 3) the primer did not have hair-pin formations or self-annealing sites and was not 3' prime complimentary.

Human SNPs that did not meet these criteria:

***OXTR***: rs53576, rs237887, rs11914920, rs7632287, rs2268495, rs237922, rs2254298, rs10490801, rs2072582, rs4686300, rs2139184, rs11706648, rs2268494, rs11131149, rs17049528, rs401015

***FOXP2***: rs2396753, rs145603, rs1852469

***AVPR1A***: rs7294536, rs10877969, rs11174815, rs3741865, rs3759292
