## Supplementary Material 2 for "Linked *OXTR* Variants Are Associated With Social Behavior Differences in Bonobos (*Pan paniscus*)"

**Supplementary Material 2 – DNA Extraction and Purification**

Bonobo DNA extraction was completed using the PureLink® Genomic DNA Mini Kit. For whole blood samples, 3 ml of blood was collected from each bonobo and stored in a freezer at -20 degrees Celsius. The blood lysate started as a combination of 200 µl of frozen whole blood samples, 20 µl Proteinase K, and 20 µl RNase A in a 1.5 ml microcentrifuge tube. The tube was then vortexed briefly and incubated at room temperature for 2 minutes. Next, 200 µl PureLink® Genomic Lysis /Binding Buffer was added to the lysate, and the lysate was vortexed. Once the lysate was homogenous, it was incubated at 55 degrees Celsius for 10 minutes in a hot bead bath to promote protein digestion. Then 200 µl 96-100% ethanol was added to the lysate, and vortexed for 5 seconds to yield a homogenous solution. Next, the lysate prepared with the genomic lysis/binding buffer and ethanol (~ 640 µl) was transferred to a PureLink® Spin Column in a collection tube. The column was then centrifuged at 10,000 x g for 1 minute at room temperature. The collection tube was discarded, and the spin column was placed in a new sterile collection tube.

For buccal swab samples, the Qiagen Omni Swab tips were ejected and placed in their own 2 ml microcentrifuge tube, and then 600 µl of phosphate buffered saline (PBS) was added to the tube. Next, 20 µl of Proteinase K was added to a large 15 ml centrifuge tube, as well as 600 µl of swab lysate, which were mixed well via pipetting. Afterwards, 600 µl of PureLink® Genomic Lysis/Binding Buffer was added to the lysate and mixed by vortexing. The lysate was incubated at 55 degrees Celsius in a hot bead bath for 15 minutes. The tube was centrifuged to collect any leftover lysate that may be on the cap, and then 200 µl of 96-100% ethanol was added to the lysate and vortexed for 5 seconds to get a homogenous solution. This solution was then transferred PureLink® Spin Column in a collection tube and the column was then centrifuged at 10,000 x g for 1 minute at room temperature. The collection tube was discarded, and the spin column was placed in a new sterile collection tube.

For DNA washing, 500 µl of ethanol prepared wash buffer 1 was added to the spin column. Then the column was centrifuged at 10,000 x g at room temperature for 1 minute. The collection tube was discarded afterwards and replaced with a new sterile tube. The DNA was washed a second time with ethanol prepared wash buffer 2 and centrifuged at maximum speed, 15,000 x g, for 3 minutes at room temperature, and then the collection tube was discarded. To ensure DNA binding conditions were still met while removing salt and protein contaminants, 96-100% ethanol was added to PureLink® Genomic Wash Buffer 1 and 2. To elute the DNA, the spin column was placed in a 1.5 ml microcentrifuge tube. Then 100 µl of PureLink® Genomic Elution Buffer was added to the column and incubated at room temperature for 1 minute before centrifugation at maximum speed. After centrifugation, the 1.5 ml microcentrifuge tube contained purified genomic DNA. At the end, the DNA concentrations extracted from blood were checked with a Nanodrop.

Conventional polymerase chain reaction (PCR; PCR Master Mix (2X) from Thermo Scientific) was used to amplify the target DNA sequence. A combination of 25 µl of PCR Master Mix, 1 µM of forward primer, 1 µM of reverse primer, and 1 µg of template DNA was then added to PCR tube kept on ice. Next, 22 µL of water was added to the mix making it a total of 50 µL. The mix ratio was repeated for each bonobo DNA sample and placed into the thermal cycler upon completion. The thermocycler conditions were set as follows:

1.) Initial denaturation - temperature: 94 degrees Celsius, time: 1 minute
2.) Denaturation – temperature: 94 degrees Celsius, time: 20 seconds
3.) Annealing – temperature: 55 degrees Celsius, time: 20 seconds
4.) Extension – temperature: 72 degrees Celsius, time: 20 seconds
5.) Final extension – 72 degrees Celsius, time: hold

The initial denaturation and final extension phases ran for one cycle each while the annealing and extension phases ran for a total of 35 cycles. Once the thermocycler completed the protocol, the PCR products were loaded onto 1.3% mini agarose gels for visualization. The visualized images served as a check to be sure that the PCR reaction was successful and produced the target amplicon. The gel was made of 1.3 g of agarose, 100 ml of TAE buffer, and 2 µl of ethidium bromide. The TAE buffer was a combination of 20 ml TAE buffer, 980 ml of dH2O, and 2 µl of ethidium bromide. 250 ml of TAE buffer and 2.5 µl of ethidium bromide was added to the electrophoresis cell. For each well, the PCR product and purple loading dye was loaded with a ratio of 10:2 as suggested by the Quick-Load Purple 100 bp DNA ladder guide. The cells ran at 90 volts and 400 amps for 55 minutes. Afterwards the gels were placed in a Bio-Rad ChemiDoc XRS+ System.

The third objective, gel purification, was conducted using a Zymoclean™ Gel DNA Recovery kit. The gels were placed on a UV transilluminator, and then each band was excised using a scalpel and transferred to a 1.5 ml microcentrifuge tubes. The microcentrifuge tubes' mass was 1.00 gram each. The total mass of the gel was calculated by taking the mass of the gel piece inside the microcentrifuge tube and subtracting the microcentrifuge tubes' mass. The mass number was then multiplied by 3 to figure out how many volumes (µl) of agarose dissolving buffer (ADB) to add to the microcentrifuge tube. Once ADB was added to the microcentrifuge tubes, they were incubated in a 55 degrees Celsius hot bead bath until the gel piece was completely dissolved and the solution was homogenous. Afterwards, the melted agarose solution was transferred to a Zymo-Spin™ Column in a collection tube and centrifuged for 60 seconds at 15,000 x g. The flow through was discarded from the collection tube so that it could be used again. For DNA washing, 24 ml of 96-100% ethanol was added to the 6 ml DNA wash buffer and 96 ml of 96-100% ethanol was added to the 24 ml DNA wash buffer. Next, 200 µl of DNA wash buffer was added to the column, and then the column was spun again for 30 seconds 15,000 x g. The flow through was discarded and the washing step was repeated. The final steps included placing the spin column into a 1.5 ml microcentrifuge tube, adding 10 µl of DNA elution Buffer directly to the center of the spin column, and centrifuging for 60 seconds. The final mass of the purified DNA products was determined using both gel visualization (using 1 µl of DNA mixed with 2 µl dye and 3 µl water for clarity), and a nanodrop machine. The remaining extracted DNA from the gels were sent to GENEWIZ for Sanger sequencing.
