## Supplementary figures and images for "Linked *OXTR* Variants Are Associated With Social Behavior Differences in Bonobos (*Pan paniscus*)"

### Supplementary Material 3

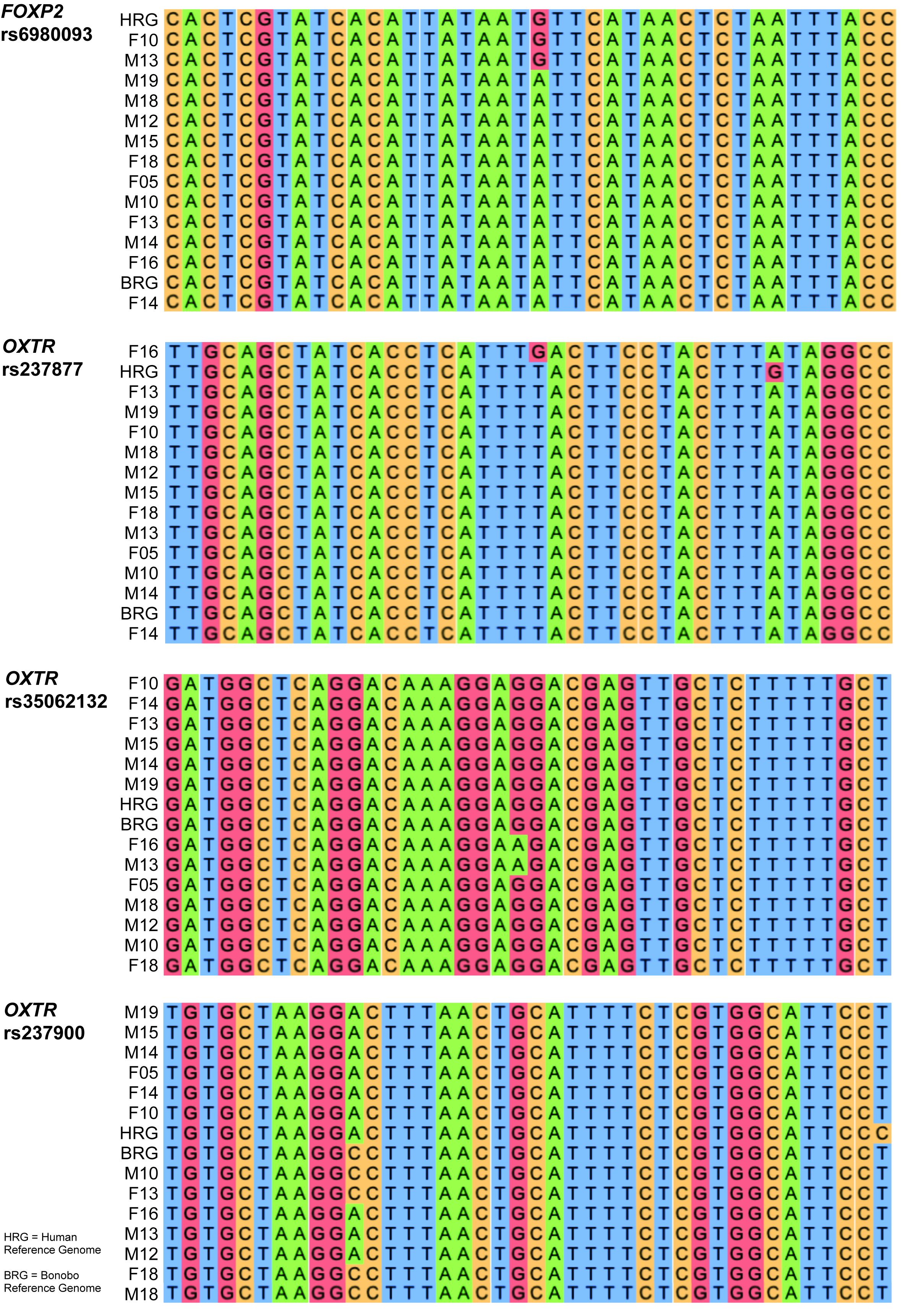
