## Supplementary Material 4 for "Linked *OXTR* Variants Are Associated With Social Behavior Differences in Bonobos (*Pan paniscus*)"

### Supplementary Material 4 – Individual Sequences and Reference Genomes for Alignments

**FOXP2_rs6980093**

>HumanReferenceGenome_UCSC_FOXP2_rs6980093
TATAAGAAATAATTTTGAAATCAGAAATATGTTTTAACTTCTTTGAGGTTACAGGACACTTGACAAGATTTTTAGGGCTTAA
GTCCAAAGGACACCGTAAAATTTACATGGATGTAGTTACAGAATTATTAAGGAATGATTATTTTCACCATTAGTTTGTATCT
TATTTTATTGAAAAGAACACTCGTATCACATTATAATGTTCATAACTCTAATTTACCACATAATATGCTAAAGATAATCT

>BonoboReferenceGenome_UCSC_FOXP2_rs6980093
TATAAGAAATAATTTTGAAATCAGAAATATGTTTTAACCTCTTTGAGGTTACAGGACACTTCACAAGATTTTTAGGGCTTAA
GTCCAAAGGACACCATAAAATTTACATGGATGTAGTTACAAAATTATTAAGGAATGATTATTTTCACCATTAGTTTGTATCT
TATTTTATTGAAAGAACACTCGTATCACATTATAATATTCATAACTCTAATTTACCACATAATATGCTAAAGATAATCTA

>F14_FOXP2_rs6980093
TTATAGCTACCAACAGGCAACCATTTTCTAAGTGGATATTTGTACATTTTTATTTATATACATATTTTCTACAATCATATTTTA
GATAATTCAATATAAGAAATAATTTTGAAATCAGAAATATGTTTTAACCTCTTTGAGGTTACAGGACACTTCACAAGATTTT
TAGGGCTTAAGTCCAAAGGACACCATAAAATTTACATGGATGTAGTTACAAAATTATTAAGGAATGATTATTTTCACCATTA
GTTTGTATCTTATTTTATTGAAAGAACACTCGTATCACATTATAATATTCATAACTCTAATTTACCACATAATATGCTAAAGAT
AATCTAACCTAAATTCTTAANGGCTAGNNNNNNNNNNNNNNNN

>F16_FOXP2_rs6980093
NTATAGCTANCCAACAGGCAACCATTTTCTAAGTGGATATTTGTACATTTTTATTTATATACATATTTTCTACAATCATATTTT
AGATAATTCAATATAAGAAATAATTTTGAAATCAGAAATATGTTTTAACCTCTTTGAGGTTACAGGACACTTCACAAGATTT
TTAGGGCTTAAGTCCAAAGGACACCATAAAATTTACATGGATGTAGTTACAAAATTATTAAGGAATGATTATTTTCACCATT
AGTTTGTATCTTATTTTATTGAAAGAACACTCGTATCACATTATAATATTCATAACTCTAATTTACCACATAATATGCTAAAGA
TAATCTAACCTAAATTCTTAANGGCTANNNNNNNNNNCNNNNNTNNN

>M14_FOXP2_rs6980093
TTATAGCTACCCAACAGGCAACCATTTTCTAAGTGGATATTTGTACATTTTTATTTATATACATATTTTCTACAATCATATTTT
AGATAATTCAATATAAGAAATAATTTTGAAATCAGAAATATGTTTTAACCTCTTTGAGGTTACAGGACACTTCACAAGATTT
TTAGGGCTTAAGTCCAAAGGACACCATAAAATTTACATGGATGTAGTTACAAAATTATTAAGGAATGATTATTTTCACCATT
AGTTTGTATCTTATTTTATTGAAAGAACACTCGTATCACATTATAATATTCATAACTCTAATTTACCACATAATATGCTAAAGA
TAATCTAACCTAAANTCTTAANGGCTAGANNNGNNNNNNNNNNNN

>F13_FOXP2_rs6980093
TATAGCTANNCACAGGCAACCATTTTCTAAGTGGATATTTGTACATTTTTATTTATATACATATTTTCTACAATCATATTTTAG
ATAATTCAATATAAGAAATAATTTTGAAATCAGAAATATGTTTTAACCTCTTTGAGGTTACAGGACACTTCACAAGATTTTTA
GGGCTTAAGTCCAAAGGACACCATAAAATTTACATGGATGTAGTTACAAAATTATTAAGGAATGATTATTTTCACCATTAGT
TTGTATCTTATTTTATTGAAAGAACACTCGTATCACATTATAATATTCATAACTCTAATTTACCACATAATATGCTAAAGATAA
TCTAACCTAAANTCTTAANGGCTANNNNNNNNNNNNNNNNNN

>M10_FOXP2_rs6980093
NNNNNNNNNNNNNNNNNNNTTATTTATATACATATTTTCTACAATCATATTTTAGATAATTCAATATAAGAAATAATTTTGAAAT
CAGAAATATGTTTTAACCTCTTTGAGGTTACAGGACACTTCACAAGATTTTTAGGGCTTAAGTCCAAAGGACACCATAAA
ATTTACATGGATGTAGTTACAAAATTATTAAGGAATGATTATTTTCACCATTAGTTTGTATCTTATTTTATTGAAAGAACACT
CGTATCACATTATAATATTCATAACTCTAATTTACCACATAATATGCTAAAGATAATCTAACCTAAATTTCTTAATGGCTAGAA
TAGAAAACAGATTTCTCCGTCAGGTGAGTTTGCCCAAGATCNNANN

>F05_FOXP2_rs6980093
NNNNNNNNNNNNGNACTTTTTATTTATATACATATTTTCTACAATCATATTTTAGATAATTCAATATAAGAAATAATTTTGAAAT
CAGAAATATGTTTTAACCTCTTTGAGGTTACAGGACACTTCACAAGATTTTTAGGGCTTAAGTCCAAAGGACACCATAAA
ATTTACATGGATGTAGTTACAAAATTATTAAGGAATGATTATTTTCACCATTAGTTTGTATCTTATTTTATTGAAAGAACACT
CGTATCACATTATAATATTCATAACTCTAATTTACCACATAATATGCTAAAGATAATCTAACCTAAATTTCTTAATGGCTAGAA
TAGAAAACAGATTTCTCCGTCAGGTGAGTNNGNNNCAAGATNNN

>M13_FOXP2_rs6980093
NNNNNNNNNNNNNNNNNACTTTTNNTTTATATACATATTTTCTACAATCATATTTTAGATAATTCAATATAAGAAATAATTTTGA
AATCAGAAATATGTTTTAACCTCTTTGAGGTTACAGGACACTTCACAAGATTTTTAGGGCTTAAGTCCAAAGGACACCAT
AAAATTTACATGGATGTAGTTACAAAATTATTAAGGAATGATTATTTTCACCATTAGTTTGTATCTTATTTTATTGAAAGAAC
ACTCGTATCACATTATAATGTTCATAACTCTAATTTACCACATAATATGCTAAAGATAATCTAACCTAAATTTCTTAATGGCTA
GAATAGAAAACAGATTTCTCCGTCAGGTGAGTTTGCCCAAGATCTAN

>F18_FOXP2_rs6980093
NNNNNNNNTNNACTTTTTATTTATATACATATTTTCTACAATCATATTTTAGATAATTCAATATAAGAAATAATTTTGAAATCA
GAAATATGTTTTAACCTCTTTGAGGTTACAGGACACTTCACAAGATTTTTAGGGCTTAAGTCCAAAGGACACCATAAAAT
TTACATGGATGTAGTTACAAAATTATTAAGGAATGATTATTTTCACCATTAGTTTGTATCTTATTTTATTGAAAGAACACTCG
TATCACATTATAATATTCATAACTCTAATTTACCACATAATATGCTAAAGATAATCTAACCTAAATTTCTTAATGGCTAGAATA
GAAAACAGATTTCTCCGTCAGGTGAGTTTGCCCAAGANNNNN

>M15_FOXP2_rs6980093
TTATAGCTACCCAACAGGCAACCATTTTCTAAGTGGATATTTGTACATTTTTATTTATATACATATTTTCTACAATCATATTTT
AGATAATTCAATATAAGAAATAATTTTGAAATCAGAAATATGTTTTAACCTCTTTGAGGTTACAGGACACTTCACAAGATTT
TTAGGGCTTAAGTCCAAAGGACACCATAAAATTTACATGGATGTAGTTACAAAATTATTAAGGAATGATTATTTTCACCATT
AGTTTGTATCTTATTTTATTGAAAGAACACTCGTATCACATTATAATATTCATAACTCTAATTTACCACATAATATGCTAAAGA
TAATCTAACCTAAATTCTTAANGGCTAGAANNGNNNNNNNNNNNNNN

>M12_FOXP2_rs6980093
TATAGCTANCCACAGGCAACCATTTTCTAAGTGGATATTTGTACATTTTTATTTATATACATATTTTCTACAATCATATTTTAG
ATAATTCAATATAAGAAATAATTTTGAAATCAGAAATATGTTTTAACCTCTTTGAGGTTACAGGACACTTCACAAGATTTTTA
GGGCTTAAGTCCAAAGGACACCATAAAATTTACATGGATGTAGTTACAAAATTATTAAGGAATGATTATTTTCACCATTAGT
TTGTATCTTATTTTATTGAAAGAACACTCGTATCACATTATAATATTCATAACTCTAATTTACCACATAATATGCTAAAGATAA
TCTAACCTAAATTCTTAATGGCTAGAANNNNNNNANNNNNNNNN

>M18_FOXP2_rs6980093
NNNNNNNNNNNACTTTTTATTTATATACATATTTTCTACAATCATATTTTAGATAATTCAATATAAGAAATAATTTTGAAATCA
GAAATATGTTTTAACCTCTTTGAGGTTACAGGACACTTCACAAGATTTTTAGGGCTTAAGTCCAAAGGACACCATAAAAT
TTACATGGATGTAGTTACAAAATTATTAAGGAATGATTATTTTCACCATTAGTTTGTATCTTATTTTATTGAAAGAACACTCG
TATCACATTATAATATTCATAACTCTAATTTACCACATAATATGCTAAAGATAATCTAACCTAAATTTCTTAATGGCTAGAATA
GAAAACAGATTTCTCCGTCAGGTGAGTTTGCCCCAAGATCTNN

>F10_FOXP2_rs6980093
TTATAGCTACCAACAGGCAACCATTTTCTAAGTGGATATTTGTACATTTTTATTTATATACATATTTTCTACAATCATATTTTA
GATAATTCAATATAAGAAATAATTTTGAAATCAGAAATATGTTTTAACCTCTTTGAGGTTACAGGACACTTCACAAGATTTT
TAGGGCTTAAGTCCAAAGGACACCATAAAATTTACATGGATGTAGTTACAAAATTATTAAGGAATGATTATTTTCACCATTA
GTTTGTATCTTATTTTATTGAAAGAACACTCGTATCACATTATAATGTTCATAACTCTAATTTACCACATAATATGCTAAAGAT
AATCTAACCTAAATTCTTAANGGCTANNNNNNNNNNNNNNNNNN

>M19_FOXP2_rs6980093
TTATAGCTACCAACAGGCAACCATTTTCTAAGTGGATATTTGTACATTTTTATTTATATACATATTTTCTACAATCATATTTTA
GATAATTCAATATAAGAAATAATTTTGAAATCAGAAATATGTTTTAACCTCTTTGAGGTTACAGGACACTTCACAAGATTTT
TAGGGCTTAAGTCCAAAGGACACCATAAAATTTACATGGATGTAGTTACAAAATTATTAAGGAATGATTATTTTCACCATTA
GTTTGTATCTTATTTTATTGAAAGAACACTCGTATCACATTATAATATTCATAACTCTAATTTACCACATAATATGCTAAAGAT
AATCTAACCTAAATTNNAATGGCTANNNNNNNNNNNNNNNNN

**OXTR_1_rs237877**

>HumanReferenceGenome_UCSC_OXTR_1_rs237877
GATGCAGAGCTTTGTCATTTTTACAGTTTTTGCAGCTATCACCTCATTTTACTTCCTACTTTGTAGGCCAGTGCCTCTAAAACT
GAGTGAAGGACCAGTTTTAATATTATTATTCTAATCCACTGCACAGCTATACTTTTGCAAAACGCAACAAAAATTAATCACTGG
AAAAATGAAATGAAAACAGCCGTACAGTTACAAGCACCAATGTTTATTGTTAGATTCAATAAACATAAAATTACTGC

>BonoboReference_UCSC_OXTR_1_rs237877
GATGCAGAGCTTTGTCATTTTTACAGTTTTTGCAGCTATCACCTCATTTTACTTCCTACTTTATAGGCCAGTGCCTCTAAAACTG
AGTGAAGGACCAGTTTTAATATTATTATTCTAACCCATTGCACAGCTATACTTTTGCAAAATACAACAAAAATTAATCACTGGA
AAAATGAAATGAAAACAGCCGTACAGTTACAAACACCAATGTTTATTGTTAGATTCAATAAACACAAAATTACTGC

>F14_OXTR_1_rs237877
NNNNNNNNCNNNTGGGTGTNNNNTGGTTATCCCAGATTGTACAGAATTCTATTCTTTTGTTGCTACTGTTTATAACNAAAG
ATGCAGAGCTTTGTCATTTTTACAGTTTTTGCAGCTATCACCTCATTTTACTTCCTACTTTATAGGCCAGTGCCTCTAAAACTGA

GTGAAGGACCAGTTTTAATATTATTATTCTAACCCATTGCACAGCTATACTTTTGCAAAATACAACAAAAATTAATCACTGGAA
AAATGAAATGAAAACAGCCGTACAGTTACAAACACCAATGTTTATTGTTAGATTCAATAAACACAAAATTACTGCATCAAATTT
CTATGAAAATTTCTAAGCACTCTCAACTTCTATAATTACCTTGTTGGGAGACAGGACACACTGATTGTGAACTGCCTTTTAAGT
AGCAAGGTAATAGGGGAGGAAGCTGGGATAN

>F16_OXTR_1_rs237877
NNNNNNNNNNNNNNNNNNNNNNNNGGGTGTTACCATGGTTATCCCAGATTGTACAGAATTCTATTCTTTTGTTGCTACTGT
TNATAANNAAAGATGCAGAGCTTTGTCATTTTTACAGTTTTTGCAGCTATCACCTCATTTGACTTCCTACTTTATAGGCCAGTG
CCTCTAAAACTGAGTGAAGGACCAGTTTTAATATTATTATTCTAACCCATTGCACAGCTATACTTTTGCAAAATACAACAAAAA
TTAATCACTGGAAAAANNATATTGNNNCANTN

>M14_OXTR_1_rs237877
NNNNNNNNNNCTNNNNGGGTGTTACCATGGTTATCCCAGATTGTACAGAATTCTATTCTTTTGTTGCTACTGTTTATAACAA
AAGATGCAGAGCTTTGTCATTTTTACAGTTTTTGCAGCTATCACCTCATTTTACTTCCTACTTTATAGGCCAGTGCCTCTAAAAC
TGAGTGAAGGACCAGTTTTAATATTATTATTCTAACCCATTGCACAGCTATACTTTTGCAAAATACAACAAAAATTAATCACTG
GAAAAATGAAATGAAAACAGCCGTACAGTTACAAACACCAATGTTTATTGTTAGATTCAATAAACACAAAATTACTGCATCAA
ATTTCTATGAATATTTCTAAGCACTCTCAACTTCTATAATTACCTTGTTGGGAGACAGGACACACTGATTGTGAACTGCCTTTTA
AGTAGCAAGGTAATAGGGGAGGAAGCTGGGATAN

>F13_OXTR_1_rs237877
NNNNNNNNNNNNNNNNCNNGNNGTGTTACCATGGTTATCCCAGATTGTACAGAATTCTATTCTTTTGTTGCTACTGTTTATA
ACAAAAGATGCAGAGCTTTGTCATTTTTACAGTTTTTGCAGCTATCACCTCATTTTACTTCCTACTTTATAGGCCAGTGCCTCTA
AAACTGAGTGAAGGACCAGTTTTAATATTATTATTCTAACCCATTGCACAGCTATACTTTTGCAAAATACAACAAAAATTAATC
ACTGGAAAAATGAAATGAAAACAGCCGTACAGTTACAAACACCAATGTTTATTGTTAGATTCAATAATCACAAAATTACTGC

>M10_OXTR_1_rs237877
NANNNNNNANNNNNTCNNNNNGGTGTTACCATGGTTATCCCAGATTGTACAGAATTCTATTCTTTTGTTGCTACTGTTTATA
ACAAAAGATGCAGAGCTTTGTCATTTTTACAGTTTTTGCAGCTATCACCTCATTTTACTTCCTACTTTATAGGCCAGTGCCTCTA
AAACTGAGTGAAGGACCAGTTTTAATATTATTATTCTAACCCATTGCACAGCTATACTTTTGCAAAATACAACAAAAATTAATC
ACTGGAAAAATGAAATGAAAACAGCCGTACAGTTACAAACACCAATGTTTATTGTTAGATTCAATAAACACAAAATTACTGCA
TCAAATTTCTATGAATATTTCTAAGCACTCTCAACTTCTATAATTACCTTGTTGGGAGACAGGACACACTGATTGTGAACTGCC
TTTTAAGTAGCAAGGTAATAGGGGAGGANNNTGGGATAA

>F05_OXTR_1_rs237877
NNNNNNNNCGTGGGTGTTACCATGGTTATCCCAGATTGTACAGAATTCTATTCTTTTGTTGCTACTGTTTATAACAAAAGATG
CAGAGCTTTGTCATTTTTACAGTTTTTGCAGCTATCACCTCATTTTACTTCCTACTTTATAGGCCAGTGCCTCTAAAACTGAGTG
AAGGACCAGTTTTAATATTATTATTCTAACCCATTGCACAGCTATACTTTTGCAAAATACAACAAAAATTAATCACTGGAAAAA
TGAAATGAAAACAGCCGTACAGTTACAAACACCAATGTTTATTGTTAGATTCAATAAACACAAAATTACTGCATCAAATTTCTA
TGAATATTTCTAAGCACTCTCAACTTCTATAATTACCTTGTTGGGAGACAGGACACACTGATTGTGAACTGCCTTTTAAGTAGC
AAGGTAATAGGGGAGGAAGCTGGGGATNAANNNNNCCATGGTAAACACCCANNATGTATTTGGGCCCTTTTATGNNTGT
NAGATNGGGGN

>M13_OXTR_1_rs237877
NNNNNNNNNNNNNNCNNTGGGTGTTACCATGGTTATCCCAGATTGTACAGAATTCTATTCTTTTGTTGCTACTGTTTATAACA
AAAGATGCAGAGCTTTGTCATTTTTACAGTTTTTGCAGCTATCACCTCATTTTACTTCCTACTTTATAGGCCAGTGCCTCTAAA
ACTGAGTGAAGGACCAGTTTTAATATTATTATTCTAACCCATTGCACAGCTATACTTTTGCAAAATACAACAAAAATTAATCACT
GGAAAAATGAAATGAAAACAGCCGTACAGTTACAAACACCAATGTTTATTGTTAGATTCAATAAACACAAAATTACTGCATCA
AATTTCTATGAATATTTCTAAGCACTCTCAACTTCTATAATTACCTTGTTGGGAGACAGGACACACTGATTGTGAACTGCCTTT
TAAGTAGCAAGGTAATAGGGGAGGANNNTTGGGATAA

>F18_OXTR_1_rs237877
NNNNNNNNNACNNCGNGGGTGTTACCNTGGTTATCCCAGATTGTACAGAATTCTATTCTTTTGTTGCTACTGTTTATAACAA
AAGATGCAGAGCTTTGTCATTTTTACAGTTTTTGCAGCTATCACCTCATTTTACTTCCTACTTTATAGGCCAGTGCCTCTAAAAC
TGAGTGAAGGACCAGTTTTAATATTATTATTCTAACCCATTGCACAGCTATACTTTTGCAAAATACAACAAAAATTAATCACTG
GAAAAATGAAATGAAAACAGCCGTACAGTTACAAACACCAATGTTTATTGTTAGATTCAATAAACACAAAATTACTGCATCAA
ATTTCTATGAATATTTCTAAGCACTCTCAACTTCTATAATTACCTTGTTGGGAGACAGGACACACTGATTGTGAACTGCCTTTTA
AGTAGCAAGGTAATAGGGGAGGANNCTGGGATAN

>M15_OXTR_1_rs237877
NNNNANNNNCNNNNNNGNNNGTTACCATGGTTATCCCAGATTGTACAGAATTCTATTCTTTTGTTGCTACTGTTTATAACA
AAAGATGCAGAGCTTTGTCATTTTTACAGTTTTTGCAGCTATCACCTCATTTTACTTCCTACTTTATAGGCCAGTGCCTCTAAA
ACTGAGTGAAGGACCAGTTTTAATATTATTATTCTAACCCATTGCACAGCTATACTTTTGCAAAATACAACAAAAATTAATCACT
GGAAAAATGAAATGAAAACAGCCGTACAGTTACAAACACCAATGTTTATTGTTAGATTCAATAAACACAAAATTACTGCATCA
AATTTCTATGAATATTTCTAAGCACTCTCAACTTCTATAATTACCTTGTTGGGAGACAGGACACACTGATTGTGAACTGCCTTT
TAAGTAGCAAGGTAATAGGGGAGGAAGCTGGGATAN

>M12_OXTR_1_rs237877
NNNNNNNNNNNTCNNGGGTGTTACCATGGTTATCCCAGATTGTACAGAATTCTATTCTTTTGTTGCTACTGTTTATAACAAAA
GATGCAGAGCTTTGTCATTTTTACAGTTTTTGCAGCTATCACCTCATTTTACTTCCTACTTTATAGGCCAGTGCCTCTAAAACTG
AGTGAAGGACCAGTTTTAATATTATTATTCTAACCCATTGCACAGCTATACTTTTGCAAAATACAACAAAAATTAATCACTGGA
AAAATGAAATGAAAACAGCCGTACAGTTACAAACACCAATGTTTATTGTTAGATTCAATAAACACAAAATTACTGCATCAAAT
TTCTATGAATATTTCTAAGCACTCTCAACTTCTATAATTACCTTGTTGGGAGACAGGACACACTGATTGTGAACTGCCTTTTAA
GTAGCAAGGTAATAGGGGAGNNNCCTTGGGATA

>M18_OXTR_1_rs237877
NNNNNNNNNATANTCGNTGGGTGTTACCATGGTTATCCCAGATTGTACAGAATTCTATTCTTTTGTTGCTACTGTTTATAACAA
AAGATGCAGAGCTTTGTCATTTTTACAGTTTTTGCAGCTATCACCTCATTTTACTTCCTACTTTATAGGCCAGTGCCTCTAAAAC
TGAGTGAAGGACCAGTTTTAATATTATTATTCTAACCCATTGCACAGCTATACTTTTGCAAAATACAACAAAAATTAATCACTG
GAAAAATGAAATGAAAACAGCCGTACAGTTACAAACACCAATGTTTATTGTTAGATTCAATAAACACAAAATTACTGCATCAA
ATTTCTATGAATATTTCTAAGCACTCTCAACTTCTATAATTACCTTGTTGGGAGACAGGACACACTGATTGTGAACTGCCTTTTA
AGTAGCAAGGTAATAGGGGAGGANNTGGGATAA

>F10_OXTR_1_rs237877
NNNNNNNNNCGTGGGTGTTACCATGGTTATCCCAGATTGTACAGAATTCTATTCTTTTGTTGCTACTGTTTATAACAAAAGATG
CAGAGCTTTGTCATTTTTACAGTTTTTGCAGCTATCACCTCATTTTACTTCCTACTTTATAGGCCAGTGCCTCTAAAACTGAGTG
AAGGACCAGTTTTAATATTATTATTCTAACCCATTGCACAGCTATACTTTTGCAAAATACAACAAAAATTAATCACTGGAAAAA
TGAAATGAAAACAGCCGTACAGTTACAAACACCAATGTTTATTGTTAGATTCAATAAACACAAAATTACTGCATCAAATTTCTA
TGAATATTTCTAAGCACTCTCAACTTCTATAATTACCTTGTTGGGAGACAGGACACACTGATTGTGAACTGCCTTTTAAGTAGC
AAGGTAATAGGGGAGGAAGCTGGGATAN

>M19_OXTR_1_rs237877
NNNNNNNNNNNTGGGTGTTACCATGGTTATCCCAGATTGTACAGAATTCTATTCTTTTGTTGCTACTGTTTATAACAAAAGATG
CAGAGCTTTGTCATTTTTACAGTTTTTGCAGCTATCACCTCATTTTACTTCCTACTTTATAGGCCAGTGCCTCTAAAACTGAGTG
AAGGACCAGTTTTAATATTATTATTCTAACCCATTGCACAGCTATACTTTTGCAAAATACAACAAAAATTAATCACTGGAAAAA
TGAAATGAAAACAGCCGTACAGTTACAAACACCAATGTTTATTGTTAGATTCAATAAACACAAAATTACTGCATCAAATTTCTA
TGAATATTTCTAAGCACTCTCAACTTCTATAATTACCTTGTTGGGAGACAGGACACACTGATTGTGAACTGCCTTTTAAGTAGC
AAGGTAATAGGGGAGGAANNTGGGNNNNAAAAGGGCAT

**OXTR_2_ rs35062132**

>HumanReferenceGenome_UCSC_OXTR_2_rs35062132
CATACGCCATCACCTAGGAGCAGAGCACTTATGCCAGCACAGCCTGAGCCTCAGGCTGCAGCCCTGGCCCTGGCTGGTGGG
TCACGCCGTGGATGGCTGGGAGCAGCTCCTCTGGCTGGAGCTGCGATGGCTCAGGACAAAGGAGGACGAGTTGCTCTTTT
TGCTGGCACTCGTCTCTCCCAGGCGTCTGCCCTTCAGGTAGCTGGCGGAGCAGCACAGGAAGCGCTGCACGAGTTCGTGG
AAGAGGTGG

>BonoboReference_UCSC_OXTR_2_rs35062132
CTTACGCCATCACCTAGGAGCAGAGCACTTATGCCAGCACAGCCTGAGCCTCAGGCTGCAGCCCTGGCCCTGGCTGGTGGG
TCACGCCGTGGATGGCTGGGAGCAGCTCCTCTGGCTGGAGCTGCGATGGCTCAGGACAAAGGAGGACGAGTTGCTCTTTT
TGCTGGCACTCGTCTCTCCCAGGCGTTTGCCCCTCAGGTAGCTGGCGGAGCAGCACAGGAAGCGCTGCACGAGTTCGTGG
AAGAGGTGG

>F14_OXTR_2_rs35062132
NNNNNNNNNNNNNNCNCNGCTGCAGCCCTGGCCCTGGCTGGTGGGTCACGCCGTGGATGGCTGGGAGCAGCTCCTCTG
GCTGGAGCTGCGATGGCTCAGGACAAAGGAGGACGAGTTGCTCTTTTTGCTGGCACTCGTCTCTCCCAGGCGTTTGCCCCT
CAGGTAGCTGGCGGAGCAGCACAGGAAGCGCTGCACGAGTTCGTGGAAGAGGTGGCCCGTGAACAGCATGTAGATCCAG
GGGTTGCAGCAGCTGTTGAGGCTGGCCAGGAGCATGACGATGATGAAGGCCGAGGCTGAGGGGGTGGGGGCAGGAGA
AAGGAGAAAAGGGCATA

>F16_OXTR_2_rs35062132
TCACCTAGGANCAGAGCACTTATGCCAGCACAGCCTGAGCCTCAGGCTGCAGCCCTGGCCCTGGCTGGTGGGTCACGCCG
TGGATGGCTGGGAGCAGCTCCTCTGGCTGGAGCTGCGATGGCTCAGGACAAAGGAAGACGAGTTGCTCTTTTTGCTGGCA
CTCGTCTCTCCCAGGCGTTTGCCCCTCAGGTAGCTGGCGGAGCAGCACAGGAAGCGCTGCACGAGTTCGTGGAAGAGGTG
GCCCGTGAACAGCATGTAGATCCAGGGGTTGCAGCAGCTGTTGAGGCTGGCCAGGAGCATGACGATGAGAANNNNNNN
NNNNNNNNNNN

>M14_OXTR_2_rs35062132
NNNNNNNNNNNNNNNNNNNNNNNTGCNGCCCTGGCCCTGGCTGGTGGGTCACGCCGTGGATGGCTGGGAGCAGCT
CCTCTGGCTGGAGCTGCGATGGCTCAGGACAAAGGAGGACGAGTTGCTCTTTTTGCTGGCACTCGTCTCTCCCAGGCGTTT
GCCCCTCAGGTAGCTGGCGGAGCAGCACAGGAAGCGCTGCACGAGTTCGTGGAAGAGGTGGCCCGTGAACAGCATGTAG
ATCCAGGGGTTGCAGCAGCTGTTGAGGCTGGCCAGGAGCATGACGATGATGAAGGCCGAGGCTGAGGGGGTGGGGGCA
GGAGAAAGGAGAAAAGGGCAT

>F13_OXTR_2_rs35062132
NNNNNNNNNNNNNNNTCNNGCTGCAGCCCTGGCCCTGGCTGGTGGGTCACGCCGTGGATGGCTGGGAGCAGCTCCTCT
GGCTGGAGCTGCGATGGCTCAGGACAAAGGAGGACGAGTTGCTCTTTTTGCTGGCACTCGTCTCTCCCAGGCGTTTGCCCC
TCAGGTAGCTGGCGGAGCAGCACAGGAAGCGCTGCACGAGTTCGTGGAAGAGGTGGCCCGTGAACAGCATGTAGATCCA
GGGGTTGCAGCAGCTGTTGAGGCTGGCCAGGAGCATGACGATGATGAAGGCCGAGGCTGAGGGGGTGGGGGCAGGAG
AAAGGAGA

>M10_OXTR_2_rs35062132
TCCAGCACAGCCTGAGCCTCAGGCTGCAGCCCTGGCCCTGGCTGGTGGGTCACGCCGTGGATGGCTGGGAGCAGCTCCTC
TGGCTGGAGCTGCGATGGCTCAGGACAAAGGAGGACGAGTTGCTCTTTTTGCTGGCACTCGTCTCTCCCAGGCGTTTGCCC
CTCAGGTAGCTGGCGGAGCAGCACAGGAAGCGCTGCACGAGTTCGTGGAAGAGGTGGCCCGTGAACAGCATGTAGATCC
NGGNNTNNNNNNNNNNNN

>F05_OXTR_2_rs35062132

TCACCTAGGANCAGAGCACTTATGCCAGCACAGCCTGAGCCTCAGGCTGCAGCCCTGGCCCTGGCTGGTGGGTCACGCCG
TGGATGGCTGGGAGCAGCTCCTCTGGCTGGAGCTGCGATGGCTCAGGACAAAGGAGGACGAGTTGCTCTTTTTGCTGGCA
CTCGTCTCTCCCAGGCGTTTGCCCCTCAGGTAGCTGGCGGAGCAGCACAGGAAGCGCTGCACGAGTTCGTGGAAGAGGTG
GCCCGTGAACAGCATGTAGATCCAGGGGTTGCAGCAGCTGTTGAGGCTGGCCAGGAGCATGACGATGATGAAGNCCGAN
NNNNNNNNNN

>M13_OXTR_2_rs35062132
TCACCTAGGANCAGAGCACTTATGCCAGCACAGCCTGAGCCTCAGGCTGCAGCCCTGGCCCTGGCTGGTGGGTCACGCCG
TGGATGGCTGGGAGCAGCTCCTCTGGCTGGAGCTGCGATGGCTCAGGACAAAGGAAGACGAGTTGCTCTTTTTGCTGGCA
CTCGTCTCTCCCAGGCGTTTGCCCCTCAGGTAGCTGGCGGAGCAGCACAGGAAGCGCTGCACGAGTTCGTGGAAGAGGTG
GCCCGTGAACAGCATGTAGATCCAGGGGTTGCAGCAGCTGTTGAGGCTGGCCAGGAGCATGACGATGANGAAGNCNCGA
NNNNNNNNNNNN

>F18_OXTR_2_rs35062132
NNAGTAAATAGTTTNNCAGNNAATAGTTNGGNAAAGTTTGCGNNNTNNCACAGGCTTNGTGGTGGCACNGTNGTNGTT
TATATGGNTCATTCAATANGTGTTAGGTGAAATCCAGGCGAGTCCATGATTCNCCATGATGTGGAAAAAANGGGTGAGNTC
GTAGGGTCGTCCGANNGTNGTGAGAAGTAAGTTGACCGCAGNGTNATTCATGGTGGTNANNACANGAACGNATAAGTGT
ATNANNNNNATGACAANNGCAAGGTGATTTTNTTNGACCGGGGTGTCNTCANNCAAGGCATTGTGAGAATAGTGTAAGC
GGCGAAGAAGTTAAGANGNGAACACGTCAATANGGGANAATANTTGAAGAGATNGGAAAACNTGAAANGCGGTGAACA
TTGGAAAACCTTCTCCGGCCNGGAAACTNNTAAGGATCCCGACGCCGGCGAGATCCAGTCCCANCATNNCANTCGGAAA
GNGAAGGAATAAAGAGGATATTGNANTTTGACCAAAAGGTNTGGGTGATTNNNNNCANGAACGTAAAANTCCATNTCAA
AGAGAATAAGGAGGACTCGGAATTGCAGAATATCCAGCACAGCCTGAGCCTCAGGCTGCAGCCCTGGCCCTGGCTGGTGG
GTCACGCCGTGGATGGCTGGGAGCAGCTCCTCTGGCTGGAGCTGCGATGGCTCAGGACAAAGGAGGACGAGTTGCTCTTT
TTGCTGGCACTCGTCTCTCCCAGGCGTTTGCCCCTCAGGTAGCTGGCGGAGCAGCACAGGAAGCGCTGCACGAGTTCGTG
GAAGAGGTGGCCCGTGANCAGCATGTAGNNCCNNNNNNNCNNNNNNNNNNNC

>M15_OXTR_2_rs35062132
NNNNNNNNNNNNNNNNNTCNNGCTGCAGCCCTGGCCCTGGCTGGTGGGTCACGCCGTGGATGGCTGGGAGCAGCTCC
TCTGGCTGGAGCTGCGATGGCTCAGGACAAAGGAGGACGAGTTGCTCTTTTTGCTGGCACTCGTCTCTCCCAGGCGTTTGC
CCCTCAGGTAGCTGGCGGAGCAGCACAGGAAGCGCTGCACGAGTTCGTGGAAGAGGTGGCCCGTGAACAGCATGTAGAT
CCAGGGGTTGCAGCAGCTGTTGAGGCTGGCCAGGAGCATGACGATGATGAAGGCCGAGGCTGAGGGGGTGGGGGCAGG
AGAAAGGAGAAAAGGGCATA

>M12_OXTR_2_rs35062132
TCACCTAGGNNNGAGCACTTATGCCAGCACAGCCTGAGCCTCAGGCTGCAGCCCTGGCCCTGGCTGGTGGGTCACGCCGT
GGATGGCTGGGAGCAGCTCCTCTGGCTGGAGCTGCGATGGCTCAGGACAAAGGAGGACGAGTTGCTCTTTTTGCTGGCAC
TCGTCTCTCCCAGGCGTTTGCCCCTCAGGTAGCTGGCGGAGCAGCACAGGAAGCGCTGCACGAGTTCGTGGAAGAGGTGG
CCCGTGAACAGCATGTAGATCCAGGGGTTGCAGCAGCTGTTGAGGCTGGCCAGGAGCATGACGATGANGAAGACNNNNN
NNNNNNNNNNNN

>M18_OXTR_2_rs35062132
TCACCTAGGANCAGAGCACTTATGCCAGCACAGCCTGAGCCTCAGGCTGCAGCCCTGGCCCTGGCTGGTGGGTCACGCCG
TGGATGGCTGGGAGCAGCTCCTCTGGCTGGAGCTGCGATGGCTCAGGACAAAGGAGGACGAGTTGCTCTTTTTGCTGGCA
CTCGTCTCTCCCAGGCGTTTGCCCCTCAGGTAGCTGGCGGAGCAGCACAGGAAGCGCTGCACGAGTTCGTGGAAGAGGTG
GCCCGTGAACAGCATGTAGATCCAGGGGTTGCAGCAGCTGTTGAGGCTGGCCAGGAGCATGACGATGATGAAGNCCNNN
NNNNNNNNNNN

>F10_OXTR_2_rs35062132

NNNNNNNNNNNNNNNNNNNNNNNNNNNGCTNNNNNNNTGGCCCTGGCTGGTGGGTCACGCCGTGGATGGCTGGGA
GCAGCTCCTCTGGCTGGAGCTGCGATGGCTCAGGACAAAGGAGGACGAGTTGCTCTTTTTGCTGGCACTCGTCTCTCCCAG
GCGTTTGCCCCTCAGGTAGCTGGCGGAGCAGCACAGGAAGCGCTGCACGAGTTCGTGGAAGAGGTGGCCCGTGAACAGC
ATGTAGATCCAGGGGTTGCAGCAGCTGTTGAGGCTGGCCAGGAGCATGACGATGATGAAGGCCGAGGCTGAGGGGGTGG
GGGCAGGAGAAAGGAGAAAAGGGCAT

>M19_OXTR_2_rs35062132
NNNNNNNNNNNNNNNNCNNNNNGCTGCANCCCTGGCCCTGGCTGGTGGGTCACGCCGTGGATGGCTGGGAGCAGCTCC
TCTGGCTGGAGCTGCGATGGCTCAGGACAAAGGAGGACGAGTTGCTCTTTTTGCTGGCACTCGTCTCTCCCAGGCGTTTGC
CCCTCAGGTAGCTGGCGGAGCAGCACAGGAAGCGCTGCACGAGTTCGTGGAAGAGGTGGCCCGTGAACAGCATGTAGAT
CCAGGGGTTGCAGCAGCTGTTGAGGCTGGCCAGGAGCATGACGATGATGAAGGCCGAGGCTGAGGGGGTGGGGGCAGG
AGAAAGGAGAAAAGGGCAT

**OXTR_3_rs237900**

>HumanReferenceGenome_UCSC_OXTR_3_rs237900
CACCACCTCCTCCTAACCACGTGGGAAAGCAGATGGAGTCCCTTGAACCTGTTTCCACCACCACAGTAAACCATACAGCCCA
TATGCAGCCCAAGGACTGTGCTAAGGACTTTAACTGCATTTTCTCGTGGCATTCCCCTATGCCTGGTGACAGTTGGTACTTTT
ATCCTTATTGCCACCTTACAGAAGAGAAAGTGGAGGCTCAGGGAGGTTAATTGCCCTAAGTCACTTGGTCAAGGGATGACA
GAGCAGTGACTCTGTG

>BonoboReference_UCSC_OXTR_3_rs237900
CACCACCTCCTCCTAACCACGTGGGAAAGCAGATGGAGTCCCTTGAACCTGTTTCCACCACCACAGTAAACCATACAGCCCA
GATGCAGCCCAAGGACTGTGCTAAGGCCTTTAACTGCATTTTCTCGTGGCATTCCTCTATGCCTGGTGACAGTTGGTACTTTT
ATCCTTATTGCCGCCTTACAGAAGAGAAAGTGGAGGCTCAGGGAGGTTAATTGCCCTAAGTCACTTGGTCAAGGGATGACA
GAGCAGTGACTCTGTG

>F14_OXTR_3_rs237900
NNNNNNNNNNATGNNNCNNGGACTGTGCTAAGGACTTTAACTGCATTTTCTCGTGGCATTCCTCTATGCCTGGTGACAGT
TGGTACTTTTATCCTTATTGCCNCCTTACAGAAGAGAAAGTGGAGGCTCAGGGAGGTTAATTGCCCTAAGTCACTTGGTCAA
GGGATGACAGAGCAGTGACTCTGTGGGATTTCAAACCCGCTTATCCCCCAGGAAGTCCACAGACCCCTGGACATTCTGAGG
CAGCAAGATAAGGGCCTCCA

>F16_OXTR_3_rs237900
TTGAACCTGTTTCCACCACCACAGTAAACCATACAGCCCAGATGCAGCCCAAGGACTGTGCTAAGGACTTTAACTGCATTTT
CTCGTGGCATTCCTCTATGCCTGGTGACAGTTGGTACTTTTATCCTTATTGCCGCCTTACAGAAGAGAAAGTGGAGGCTCAG
GGAGGTTAATTGCCCTAAGTCACTTGGTCAAGGGATGACAGAGCAGTGACTCTGTGGGATTTCAAACCCGNNNTCNCCCA
GGAANTCCNNTNNNNNNNNNNNN

>M14_OXTR_3_rs237900
NNNNNNNCCNNNNTNNNNNNCNNGGACTGTGCTAAGGACTTTAACTGCATTTTCTCGTGGCATTCCTCTATGCCTGGNG
NCNGTTGGTACTTTTATCCTTATTGCCGCCTTACAGAAGAGAAAGTGGAGGCTCAGGGAGGTTAATTGCCCTAAGTCACTTG
GTCAAGGGATGACAGAGCAGTGACTCTGTGGGATTTCAAACCCGCTTATCCCCCAGGAAGTCCACAGACCCCTGGACATTC
TGAAGCAGCAAGATAAGGGCCTCCA

>F13_OXTR_3_rs237900
TTGAACCTGTTTTCCACCACCACAGTAAACCATACAGCCCAGATGCAGCCCAAGGACTGTGCTAAGGCCTTTAACTGCATTTT
CTCGTGGCATTCCTCTATGCCTGGTGACAGTTGGTACTTTTATCCTTATTGCCGCCTTACAGAAGAGAAAGTGGAGGCTCAG

GGAGGTTAATTGCCCTAAGTCACTTGGTCAAGGGATGACAGAGCAGTGACTCTGTGGGATTTCAAACCCGCNNNNNCCCA
GGAANTCCNCANNCNNNNNNNNN

>M10_OXTR_3_rs237900
TTGAACCTGTTTCCACCACCACAGTAAACCATACAGCCCAGATGCAGCCCAAGGACTGTGCTAAGGCCTTTAACTGCATTTT
CTCGTGGCATTCCTCTATGCCTGGTGACAGTTGGTACTTTTATCCTTATTGCCGCCTTACAGAAGAGAAAGTGGAGGCTCAG
GGAGGTTAATTGCCCTAAGTCACNTGTTCAAGGGATGACAGAGCAGTGACTCTGTGGGATTCAAACGCGATTNACCNNTN
CANNNNNNNNNNNNNNNNNA

>F05_OXTR_3_rs237900
NNNNNCNNNGNNNCNNNGACTGTGCTAAGGACTTTAACTGCATTTTCTCGTGGCATTCCTCTATGCCTGGTGACAGTTGG
TACTTTTATCCTTATTGCCACCTTACAGAAGAGAAAGTGGAGGCTCAGGGAGGTTAATTGCCCTAAGTCACTTGGTCAAGGG
ATGACAGAGCAGTGACTCTGTGGGATTTCAAACCCGCTTATCCCCCAGGAAGTCCACAGACCCCTGGACATTCTGAGGCAG
CAAGATAAGGGCCTCCACCAAN

>M13_OXTR_3_rs237900
TTGAACCTGTTTCCACCACCACAGTAAACCATACAGCCCAGATGCAGCCCAAGGACTGTGCTAAGGACTTTAACTGCATTTT
CTCGTGGCATTCCTCTATGCCTGGTGACAGTTGGTACTTTTATCCTTATTGCCGCCTTACAGAAGAGAAAGTGGAGGCTCAG
GGAGGTTAATTGCCCTAAGTCACTTGGTCAAGGGATGACAGAGCAGTGACTCTGTGGGATTTCAAACCCGCNNNNNCCCA
GGAANTCCACANNNNNNNNNNNNN

>F18_OXTR_3_rs237900
TTGAACCTGTTTCCACCACCACAGTAAACCATACAGCCCAGATGCAGCCCAAGGACTGTGCTAAGGCCTTTAACTGCATTTT
CTCGTGGCATTCCTCTATGCCTGGTGACAGTTGGTACTTTTATCCTTATTGCCGCCTTACAGAAGAGAAAGTGGAGGCTCAG
GGAGGTTAATTGCCCTAAGTCACTTGGTCAAGGGATGACAGAGCAGTGACTCTGTGGGATTTCAAACCCGCNNNNNCCCA
GGAAGTCCNCNNNCNNNNNNNNNNN

>M15_OXTR_3_rs237900
NNNNNNNNCNNNNGCNGCCNNNGACTGTGCTAAGGACTTTAACTGCATTTTCTCGTGGCATTCCTCTATGCCTGGTGACA
GTTGGTACTTTTATCCTTATTGCCNCCTTACAGAAGAGAAAGTGGAGGCTCAGGGAGGTTAATTGCCCTAAGTCACTTGGTC
AAGGGATGACAGAGCAGTGACTCTGTGGGATTTCAAACCCGCTTATCCCCCAGGAAGTCCACAGACCCCTGGACATTCTGA
AGCAGCAAGATAAGGGCCNNNNN

>M12_OXTR_3_rs237900
TTGAACCTGTTTCCACCACCACAGTAAACCATACAGCCCAGATGCAGCCCAAGGACTGTGCTAAGGACTTTAACTGCATTTT
CTCGTGGCATTCCTCTATGCCTGGTGACAGTTGGTACTTTTATCCTTATTGCCGCCTTACAGAAGAGAAAGTGGAGGCTCAG
GGAGGTTAATTGCCCTAAGTCACTTGGTCAAGGGATGACAGAGCAGTGACTCTGTGGGATTTCAAACCCGCTNNNNCCCA
NNAAGTCCNNNNNNNNNNNNNNNNN

>M18_OXTR_3_rs237900
TGAACCTGTTTCCACCACCACAGTAAACCATACAGCCCAGATGCAGCCCAAGGACTGTGCTAAGGCCTTTAACTGCATTTTCT
CGTGGCATTCCTCTATGCCTGGTGACAGTTGGTACTTTTATCCTTATTGCCGCCTTACAGAAGAGAAAGTGGAGGCTCAGGG
AGGTTAATTGCCCTAAGTCACTTGGTCAAGGGATGACAGAGCAGTGACTCTGTGGGATTTCAAACCCGCNNNNCCCCAGG
AANTCCNNNNNCNNNNNNNNNNN

>F10_OXTR_3_rs237900
NNNNNNNNNNATNNNNNNNGGACTGTGCTAAGGACTTTAACTGCATTTTCTCGTGGCATTCCTCTATGCCTGGTGACAGT
TGGTACTTTTATCCTTATTGCCGCCTTACAGAAGAGAAAGTGGAGGCTCAGGGAGGTTAATTGCCCTAAGTCACTTGGTCAA
GGGATGACAGAGCAGTGACTCTGTGGGATTTCAAACCCGCTTATCCCCCAGGAAGTCCACAGACCCCTGGACATTCTGAGG

CAGCAAGATAAGGGCCTCCCNNTAGGTCTCCTGGAGGGCCCCTTACAACTCAGCTGGTGTTAGAANAAATGCCTGGGTGTA
ACAGTGAGAAACAACCTTTCAAAAGTGTTGAACAATTTTTCAGGCAGAGGAATATGTATGCCAATCCAGGCAGCAAGATAA
GGGCCTCCAN

>M19_OXTR_3_rs237900
NNNNNNNNNNNNNNNANNNNNNNNNNNGNNTGTGCTAAGGACTTTAACTGCATTTTCTCGTGGCATTCCTCTATGCCTG
GTGACNGTTGGTACTTTTATCCTTATTGCCGCCTTACANAAGANAAAGTGGAGGCTCANGGAGGTTAATTGCCCTAAGTCA
CTTGGTCAAGGGATGACAGAGCAGTGACTCTGTGGGATTTCAAACCCGCTTATCCCCCACGAAGTCCGCNCACCCCTGGAC
ATTCTGACGCNNCAAGATAAGGGCCTCCACGAATCGGCCAACGCGCGGGGAGAGGCGGTTTGCGTATTGGGCGCTCTTCC
GCTTCCTCGCTCACTGACTCGCTGCGCTCGGTCGTTCGGCTGCGGCGAGCGGTATCAGCTCACTCAAAGGCGGTAATACGG
TTANCCACAGAAACAGGGGATAACGTAGGAAAGAACATGTGAGCAAAAGGNCAGCAAAAGGNCAGNAACCGTAAAAAG
GCCGCGTTGCTGGCGTTTTTCCATANGCTCCGCCCCCCTGACGATCATCACAAAAATCGACGCTCAANTCANAGGTGGCGA
AACCCGACAGGANTATAANNATACCATCGCGTTNCCCCNGGAAGCTCNCTCATGCGCTCTNCTGTTCTNACNCTCGCCGTT
TACCGGANACCTGTCCCGCCTNNCTCNCNNTCGGGAAGCGTGNCGCTNTTCTCATAGCTCACTCTGNANGNATCCTCAGT

**OXTR_rs2254295**

>HumanReferenceGenome_UCSC_OXTR_rs2254295
CAGAGGAAGAAGCCCCGCAAACTGGGAAAACAGGGATGGTTTCCTGAAAGGGGCACTGGATGAGGCTGCCATGTTCAGC
TGTTCAGGCTGTGCACTGCAAAGCTCCTAGGAACACCCTGTTCATAGACCATGAGGTAAAAGGCATCCCCTGGAGTTGTGCC
ACAAAGCAGCCCACACAATGGACTTGGGCTTCAAAGGATGGGAATCCGTGGAAGAAACTGGGGTGGGCGTTCCCATGGC
AGGA

>BonoboReferenceGenome_UCSC_OXTR_rs2254295
CAGAGGAAGAAGCCCCGCAAACTGGGAAAACAGGGATGGTTTCCTGAAAGGGGCACTGGATGAGGCTGCCATGTTCAGC
TGTTCAGGCTGTGCACTGCAAAGCTCCTAGGAACACCCTGTTCGTAGACCATGAGGTAAAAGGCATCCCCTGGAGTTGTGC
CACAAAGCAGCCCACACAATGGACTTGGGCTTCAAAGGATGGGAATCCGTGGAAGAAACTGGGGTGGGCGTTCCCATGGC
AGGA

>F14_OXTR_rs2254295
NTGCTCCANNNNACAGTAACCCAGCAGAAACTGGGGGTGTCCCTCCCAGAGGTCTGTGGGTGTACCCAGAACTCTGTGAT
CAACCTTGACCACACTGTCCCACATTTATGCATGTCAGCAGCTGGCCGCAGGAGGGCGAGGGCTTGGGAGGGCCCCTTGC
CAACTCTCTTCACTTGGGGGTTGACAGATGAAAGCAGAGGTTGTGTGGACAGGAGCCTGCAGAGGCATCAGTAACTGTTT
TTTGGAGTGAATGACTTAGCATTCAGAGGAAGAAGCCCCGCAAACTGGGAAAACAGGGATGGTTTCCTGAAAGGGGCACT
GGATGAGGCTGCCATGTTCAGCTGTTCAGGCTGTGCACTGCAAAGCTCCTAGGAACACCCTGTTCGTAGACCATGAGGTAA
AAGGCATCCCCTGGAGTTGTGCCACAAAGCAGCCCNNNNNNNNNNNN

>F16_OXTR_rs2254295
NNNNNNNNNNNNNNNNNNNNAGGTCTGTGGGTGTACCCAGAACTCTGTGATCAACCTTGACCACACTGTCCCACATTTA
TGCATGTCAGCAGCTGGCCGCAGGAGGGCGAGGGCTTGGGAGGGCCCCTTGCCAACTCTCTTCACTTGGGGGTTGACAG
ATGAAAGCAGAGGTTGTGTGGACAGGAGCCTGCAGAGGCATCAGTAACTGTTTTTTGGAGTGAATGACTTAGCATTCAGA
GGAAGAAGCCCCGCAAACTGGGAAAACAGGGATGGTTTCCTGAAAGGGGCACTGGATGAGGCTGCCATGTTCAGCTGTT
CAGGCTGTGCACTGCAAAGCTCCTAGGAACACCCTGTTCGTAGACCATGAGGTAAAAGGCATCCCCTGGAGTTGTGCCACA
AAGCAGCCCACACAATGGACTTGGGCTTCAAAGGATGGGAATCCGTGGAA

>M14_OXTR_rs2254295
NNNNNNNNNNNNNNACTNNNNNNNTGGAGTGATGACTTAGCATTCAGAGGAAGAAGCCCCGCAAACTGGGAAAACAGG
GATGGTTTCCTGAAAGGGGCACTGGATGAGGCTGCCATGTTCAGCTGTTCAGGCTGTGCACTGCAAAGCTCCTAGGAACAC

CCTGTTCGTAGACCATGAGGTAAAAGGCATCCCCTGGAGTTGTGCCACAAAGCAGCCCACACAATGGACTTGGGCTTCAAA
GGATGGGAATCCGTGGAA

>F13_OXTR_rs2254295
NNNNNNGNGNNGNCNNNNNNAGGTCTGTGGGTGTACCCAGAACTCTGTGATCAACCTTGACCACACTGTCCCACATTTA
TGCATGTCAGCAGCTGGCCGCAGGAGGGCGAGGGCTTGGGAGGGCCCCTTGCCAACTCTCTTCACTTGGGGGTTGACAG
ATGAAAGCAGAGGTTGTGTGGACAGGAGCCTGCAGAGGCATCAGTAACTGTTTTTTGGAGTGAATGACTTAGCATTCAGA
GGAAGAAGCCCCGCAAACTGGGAAAACAGGGATGGTTTCCTGAAAGGGGCACTGGATGAGGCTGCCATGTTCAGCTGTT
CAGGCTGTGCACTGCAAAGCTCCTAGGAACACCCTGTTCGTAGACCATGAGGTAAAAGGCATCCCCTGGAGTTGTGCCACA
AAGCAGCCCACACAATGGACTTGGGCTTCAAAGGATGGGAATCCGTGGAA

>M10_OXTR_rs2254295
NNNNNGGNNNCNCCNGAGGTCTGTGGGTGTACCCAGAACTCTGTGATCAACCTTGACCACACTGTCCCACATTTATGCATG
TCAGCAGCTGGCCGCAGGAGGGCGAGGGCTTGGGAGGGCCCCTTGCCAACTCTCTTCACTTGGGGGTTGACAGATGAAA
GCAGAGGTTGTGTGGACAGGAGCCTGCAGAGGCATCAGTAACTGTTTTTTGGAGTGAATGACTTAGCATTCAGAGGAAGA
AGCCCCGCAAACTGGGAAAACAGGGATGGTTTCCTGAAAGGGGCACTGGATGAGGCTGCCATGTTCAGCTGTTCAGGCTG
TGCACTGCAAAGCTCCTAGGAACACCCTGTTCGTAGACCATGAGGTAAAAGGCATCCCCTGGAGTTGTGCCACAAAGCAGC
CCACACAATGGACTTGGGCTTCAAAGGATGGAAATCCGTGGAA

>F05_OXTR_rs2254295
NTGCTCCACNNNACAGTAACCCAGCAGAAACTGGGGGTGTCCCTCCCAGAGGTCTGTGGGTGTACCCAGAACTCTGTGAT
CAACCTTGACCACACTGTCCCACATTTATGCATGTCAGCAGCTGGCCGCAGGAGGGCGAGGGCTTGGGAGGGCCCCTTGC
CAACTCTCTTCACTTGGGGGTTGACAGATGAAAGCAGAGGTTGTGTGGACAGGAGCCTGCAGAGGCATCAGTAACTGTTT
TTTGGAGTGAATGACTTAGCATTCAGAGGAAGAAGCCCCGCAAACTGGGAAAACAGGGATGGTTTCCTGAAAGGGGCACT
GGATGAGGCTGCCATGTTCAGCTGTTCAGGCTGTGCACTGCAAAGCTCCTAGGAACACCCTGTTCGTAGACCATGAGGTAA
AAGGCATCCCCTGGAGTTGTGCCACAAAGCAGNNNNNNNNNNNNNNNN

>M13_OXTR_rs2254295
NNNNNNNNNNNNCCNNNNNNGGTCTGTGGGTGTACCCAGAACTCTGTGATCAACCTTGACCACACTGTCCCACATTTAT
GCATGTCAGCAGCTGGCCGCAGGAGGGCGAGGGCTTGGGAGGGCCCCTTGCCAACTCTCTTCACTTGGGGGTTGACAGA
TGAAAGCAGAGGTTGTGTGGACAGGAGCCTGCAGAGGCATCAGTAACTGTTTTTTGGAGTGAATGACTTAGCATTCAGAG
GAAGAAGCCCCGCAAACTGGGAAAACAGGGATGGTTTCCTGAAAGGGGCACTGGATGAGGCTGCCATGTTCAGCTGTTCA
GGCTGTGCACTGCAAAGCTCCTAGGAACACCCTGTTCGTAGACCATGAGGTAAAAGGCATCCCCTGGAGTTGTGCCACAAA
GCAGCCCACACAATGGACTTGGGCTTCAAAGGATGGGAATCCGTGGAA

>F18_OXTR_rs2254295
NNNNNNNNNTNNNCNNGAGGTCTGTGGGTGTACCCAGAACTCTGTGATCAACCTTGACCACACTGTCCCACATTTATGCA
TGTCAGCAGCTGGCCGCAGGAGGGCGAGGGCTTGGGAGGGCCCCTTGCCAACTCTCTTCACTTGGGGGTTGACAGATGA
AAGCAGAGGTTGTGTGGACAGGAGCCTGCAGAGGCATCAGTAACTGTTTTTTGGAGTGAATGACTTAGCATTCAGAGGAA
GAAGCCCCGCAAACTGGGAAAACAGGGATGGTTTCCTGAAAGGGGCACTGGATGAGGCTGCCATGTTCAGCTGTTCAGGC
TGTGCACTGCAAAGCTCCTAGGAACACCCTGTTCGTAGACCATGAGGTAAAAGGCATCCCCTGGAGTTGTGCCACAAAGCA
GCCCACACAATGGACTTGGGCTTCAAAGGATGGGAATCCGTGGAA

>M15_OXTR_rs2254295
TTGCTCCACCTGACAGTAACCCAGCAGAAACTGGGGGTGTCCCTCCCAGAGGTCTGTGGGTGTACCCAGAACTCTGTGATC
AACCTTGACCACACTGTCCCACATTTATGCATGTCAGCAGCTGGCCGCAGGAGGGCGAGGGCTTGGGAGGGCCCCTTGCC
AACTCTCTTCACTTGGGGGTTGACAGATGAAAGCAGAGGTTGTGTGGACAGGAGCCTGCAGAGGCATCAGTAACTGTTTTT
TGGAGTGAATGACTTAGCATTCAGAGGAAGAAGCCCCGCAAACTGGGAAAACAGGGATGGTTTCCTGAAAGGGGCACTG

GATGAGGCTGCCATGTTCAGCTGTTCAGGCTGTGCACTGCAAAGCTCCTAGGAACACCCTGTTCGTAGACCATGAGGTAAA
AGGCATCCCCTGGAGTTGTGCCACAAAGCAGNNNNNNNNNNNNNNNN

>M12_OXTR_rs2254295
NNNNNNNNNNCNNNNNAGGTCTGTGGGTGTACCCAGAACTCTGTGATCAACCTTGACCACACTGTCCCACATTTATGCAT
GTCAGCAGCTGGCCGCAGGAGGGCGAGGGCTTGGGAGGGCCCCTTGCCAACTCTCTTCACTTGGGGGTTGACAGATGAA
AGCAGAGGTTGTGTGGACAGGAGCCTGCAGAGGCATCAGTAACTGTTTTTTGGAGTGAATGACTTAGCATTCAGAGGAAG
AAGCCCCGCAAACTGGGAAAACAGGGATGGTTTCCTGAAAGGGGCACTGGATGAGGCTGCCATGTTCAGCTGTTCAGGCT
GTGCACTGCAAAGCTCCTAGGAACACCCTGTTCGTAGACCATGAGGTAAAAGGCATCCCCTGGAGTTGTGCCACAAAGCAG
CCCACACAATGGACTTGGGCTTCAAAGGATGGNNATCCGTGGAA

>M18_OXTR_rs2254295
NNNNNGGGNNGNCCNNCNNAGGTCTGTGGGTGTACCCAGAACTCTGTGATCAACCTTGACCACACTGTCCCACATTTATG
CATGTCAGCAGCTGGCCGCAGGAGGGCGAGGGCTTGGGAGGGCCCCTTGCCAACTCTCTTCACTTGGGGGTTGACAGAT
GAAAGCAGAGGTTGTGTGGACAGGAGCCTGCAGAGGCATCAGTAACTGTTTTTTGGAGTGAATGACTTAGCATTCAGAGG
AAGAAGCCCCGCAAACTGGGAAAACAGGGATGGTTTCCTGAAAGGGGCACTGGATGAGGCTGCCATGTTCAGCTGTTCAG
GCTGTGCACTGCAAAGCTCCTAGGAACACCCTGTTCGTAGACCATGAGGTAAAAGGCATCCCCTGGAGTTGTGCCACAAAG
CAGCCCACACAATGGACTTGGGCTTCAAAGGANGGANNTCCGTGGAA

>F10_OXTR_rs2254295
TCAGAGGTTGTGTGGACAGGAGCCTGCAGAGGCATCAGTAACTGTTTTTTGGAGTGAATGACTTAGCATTCAGAGGAAGA
AGCCCCGCAAACTGGGAAAACAGGGATGGTTTCCTGAAAGGGGCACTGGATGAGGCTGCCATGTTCAGCTGTTCAGGCTG
TGCACTGCAAAGCTCCTAGGAACACCCTGTTCGTAGACCATGAGGTAAAAGGCATCCCCTGGAGTTGTGCCACAAAGCAGC
CCACANNNNNNNNNN

>M19_OXTR_rs2254295
TCAGAGGTTGTGTGGACAGGAGCCTGCAGAGGCATCAGTAACTGTTTTTTGGAGTGAATGACTTAGCATTCAGAGGAAGA
AGCCCCGCAAACTGGGAAAACAGGGATGGTTTCCTGAAAGGGGCACTGGATGAGGCTGCCATGTTCAGCTGTTCAGGCTG
TGCACTGCAAAGCTCCTAGGAACACCCTGTTCGTAGACCATGAGGTAAAAGGCATCCCCTGGAGTTGTGCCACAAAGCAGC
NCNCANNNNNNNNNNN

**OXTR_rs237878**

>HumanReferenceGenome_UCSC_OXTR_rs237878
CTATACTTTTGCAAAACGCAACAAAAATTAATCACTGGAAAAATGAAATGAAAACAGCCGTACAGTTACAAGCACCAATGTT
TATTGTTAGATTCAATAAACATAAAATTACTGCATCAAATTTCTATGAATATTTCTAAGCACTCTCAACTTCTATAATTACCTTGTT
GGGAGACAGGACACACTGATTGTGAACTGCCTTTTAAGTAGCAAGGTAATAGGGGAGGAAGCTGGGATCTGG

>BonoboReferenceGenome_UCSC_OXTR_rs237878
CTATACTTTTGCAAAATACAACAAAAATTAATCACTGGAAAAATGAAATGAAAACAGCCGTACAGTTACAAACACCAATGTTT
ATTGTTAGATTCAATAAACACAAAATTACTGCATCAAATTTCTATGAATATTTCTAAGCACTCTCAACTTCTATAATTACCTTGTT
GGGAGACAGGACACACTGATTGTGAACTGCCTTTTAAGTAGCAAGGTAATAGGGGAGGAAGCTGGGATCCAG

>F14_OXTR_rs237878
NNNNNNNNNNNNNNNTNGGCCNGTGCCTCTNNNACTGAGTGAAGGACCAGTTTTAATATTATTATTCTAACCCATTGCACA
GCTATACTTTTGCAAAATACAACAAAAATTAATCACTGGAAAAATGAAATGAAAACAGCCGTACAGTTACAAACACCAATGTT
TATTGTTAGATTCAATAAACACAAAATTACTGCATCAAATTTCTATGAATATTTCTAAGCACTCTCAACTTCTATAATTACCTTGT
TGGGAGACAGGACACACTGATTGTGAACTGCCTTTTAAGTAGCAAGGTAATAGGGGAGGAAGCTGGGAA

>F16_OXTR_rs237878
NNNNNNNNCNNNNNNNNNNNGCCAGTGCCTCTAANCTGAGTGAAGGACCAGTTTTAATATTATTATTCTAACCCATTGCAC
AGCTATACTTTTGCAAAATACAACAAAAATTAATCACTGGAAAAATGAAATGAAAACAGCCGTACAGTTACAAACACCAATG
TTTATTGTTAGATTCAATAAACACAAAATTACTGCATCAAATTTCTATGAATATTTCTAAGCACTCTCAACTTCTATAATTACCTT
GTTGGGAGACAGGACACACTGATTGTGAACTGCCTTTTAAGTAGCAAGGTAATAGGGGAGGAAGCTGGGAA

>M14_OXTR_rs237878
NNNNNNNNNNNNNNNNNTNNNNNGCNNNNGCCTCTAAAACTGAGTGAAGGACCAGTTTTAATATTATTATTCTAACCCAT
TGCACAGCTATACTTTTGCAAAATACAACAAAAATTAATCACTGGAAAAATGAAATGAAAACAGCCGTACAGTTACAAACAC
CAATGTTTATTGTTAGATTCAATAAACACAAAATTACTGCATCAAATTTCTATGAATATTTCTAAGCACTCTCAACTTCTATAATT
ACCTTGTTGGGAGACAGGACACACTGATTGTGAACTGCCTTTTAAGTAGCAAGGTAATAGGGGAGGAAGCTGGGAA

>F13_OXTR_rs237878
NNNNNNNNNNNNNNNNGGCCNGTGCCTCTNNACTGAGTGAAGGACCAGTTTTAATATTATTATTCTAACCCATTGCACAGC
TATACTTTTGCAAAATACAACAAAAATTAATCACTGGAAAAATGAAATGAAAACAGCCGTACAGTTACAAACACCAATGTTTA
TTGTTAGATTCAATAAACACAAAATTACTGCATCAAATTTCTATGAATATTTCTAAGCACTCTCAACTTCTATAATTACCTTGTTG
GGAGACAGGACACACTGATTGTGAACTGCCTTTTAAGTAGCAAGGTAATAGGGGAGGAAGCTGGGAA

>M10_OXTR_rs237878
NANNNNNNANNNNNTCNNNNNGGTGTTACCATGGTTATCCCAGATTGTACAGAATTCTATTCTTTTGTTGCTACTGTTTATA
ACAAAAGATGCAGAGCTTTGTCATTTTTACAGTTTTTGCAGCTATCACCTCATTTTACTTCCTACTTTATAGGCCAGTGCCTCTA
AAACTGAGTGAAGGACCAGTTTTAATATTATTATTCTAACCCATTGCACAGCTATACTTTTGCAAAATACAACAAAAATTAATC
ACTGGAAAAATGAAATGAAAACAGCCGTACAGTTACAAACACCAATGTTTATTGTTAGATTCAATAAACACAAAATTACTGCA
TCAAATTTCTATGAATATTTCTAAGCACTCTCAACTTCTATAATTACCTTGTTGGGAGACAGGACACACTGATTGTGAACTGCC
TTTTAAGTAGCAAGGTAATAGGGGAGGANNNTGGGATAA

>M13_OXTR_rs237878
NNNNNNNNNNNNNNCNNTGGGTGTTACCATGGTTATCCCAGATTGTACAGAATTCTATTCTTTTGTTGCTACTGTTTATAACA
AAAGATGCAGAGCTTTGTCATTTTTACAGTTTTTGCAGCTATCACCTCATTTTACTTCCTACTTTATAGGCCAGTGCCTCTAAA
ACTGAGTGAAGGACCAGTTTTAATATTATTATTCTAACCCATTGCACAGCTATACTTTTGCAAAATACAACAAAAATTAATCACT
GGAAAAATGAAATGAAAACAGCCGTACAGTTACAAACACCAATGTTTATTGTTAGATTCAATAAACACAAAATTACTGCATCA
AATTTCTATGAATATTTCTAAGCACTCTCAACTTCTATAATTACCTTGTTGGGAGACAGGACACACTGATTGTGAACTGCCTTT
TAAGTAGCAAGGTAATAGGGGAGGANNNTTGGGATAA

>F18_OXTR_rs237878
NNNNNNNNNACNNCGNGGGTGTTACCNTGGTTATCCCAGATTGTACAGAATTCTATTCTTTTGTTGCTACTGTTTATAACAA
AAGATGCAGAGCTTTGTCATTTTTACAGTTTTTGCAGCTATCACCTCATTTTACTTCCTACTTTATAGGCCAGTGCCTCTAAAAC
TGAGTGAAGGACCAGTTTTAATATTATTATTCTAACCCATTGCACAGCTATACTTTTGCAAAATACAACAAAAATTAATCACTG
GAAAAATGAAATGAAAACAGCCGTACAGTTACAAACACCAATGTTTATTGTTAGATTCAATAAACACAAAATTACTGCATCAA
ATTTCTATGAATATTTCTAAGCACTCTCAACTTCTATAATTACCTTGTTGGGAGACAGGACACACTGATTGTGAACTGCCTTTTA
AGTAGCAAGGTAATAGGGGAGGANNCTGGGATAN

>M15_OXTR_rs237878
NNNNNNNNNNNNNNNNNNNNNNNNNNNNNGCCTCTNNNCTGAGTGAAGGACCAGTTTTAATATTATTATTCTAACCCAT
TGCACAGCTATACTTTTGCAAAATACAACAAAAATTAATCACTGGAAAAATGAAATGAAAACAGCCGTACAGTTACAAACAC
CAATGTTTATTGTTAGATTCAATAAACACAAAATTACTGCATCAAATTTCTATGAATATTTCTAAGCACTCTCAACTTCTATAATT
ACCTTGTTGGGAGACAGGACACACTGATTGTGAACTGCCTTTTAAGTAGCAAGGTAATAGGGGAGGAAGCTGGGAA

>M12_OXTR_rs237878

NNNNNNNNNNNNNNNNNNNNNGCCNGNGNNNCTNNNCTGAGTGAAGGACCAGTTTTAATATTATTATTCTAACCCATT
GCACNGCNATACTTTTGCAAAATACAACAAAAATTAATCACTGGAAAAATGAAATGAAAACAGCCGTACAGTTACAAACACC
AATGTTTATTGTTAGATTCAATAAACACAAAATTACTGCATCAAATTTCTATGAATATTTCTAAGCACTCTCAACTTCTATAATTA
CCTTGTTGGGAGACAGGACACACTGATTGTGAACTGCCTTTTAAGTAGCAAGGTAATAGGGGANGAANCTGGGA

>M18_OXTR_rs237878
NNNNNNNNATANTCGNTGGGTGTTACCATGGTTATCCCAGATTGTACAGAATTCTATTCTTTTGTTGCTACTGTTTATAACAA
AAGATGCAGAGCTTTGTCATTTTTACAGTTTTTGCAGCTATCACCTCATTTTACTTCCTACTTTATAGGCCAGTGCCTCTAAAAC
TGAGTGAAGGACCAGTTTTAATATTATTATTCTAACCCATTGCACAGCTATACTTTTGCAAAATACAACAAAAATTAATCACTG
GAAAAATGAAATGAAAACAGCCGTACAGTTACAAACACCAATGTTTATTGTTAGATTCAATAAACACAAAATTACTGCATCAA
ATTTCTATGAATATTTCTAAGCACTCTCAACTTCTATAATTACCTTGTTGGGAGACAGGACACACTGATTGTGAACTGCCTTTTA
AGTAGCAAGGTAATAGGGGAGGANNTGGGATAA

>F10_OXTR_rs237878
NNNNNNNNNNNNNNNNNNNNNNNNNTNGCCTCTAAAACTGAGTGAAGGACCAGTTTTAATATTATTATTCTAACCCATT
GCACAGCTATACTTTTGCAAAATACAACAAAAATTAATCACTGGAAAAATGAAATGAAAACAGCCGTACAGTTACAAACACC
AATGTTTATTGTTAGATTCAATAAACACAAAATTACTGCATCAAATTTCTATGAATATTTCTAAGCACTCTCAACTTCTATAATTA
CCTTGTTGGGAGACAGGACACACTGATTGTGAACTGCCTTTTAAGTAGCAAGGTAATAGGGGAGGAAGCTGGGAA

>M19_OXTR_rs237878
NNNNNNNNNNANNGGCCAGTGCCTCTAAAACTGAGTGAAGGACCAGTTTTAATATTATTATTCTAACCCATTGCACAGCTA
TACTTTTGCAAAATACAACAAAAATTAATCACTGGAAAAATGAAATGAAAACAGCCGTACAGTTACAAACACCAATGTTTATT
GTTAGATTCAATAAACACAAAATTACTGCATCAAATTTCTATGAATATTTCTAAGCACTCTCAACTTCTATAATTACCTTGTTGG
GAGACAGGACACACTGATTGTGAACTGCCTTTTAAGTAGCAAGGTAATAGGGGAGGAAGCTGGGAA

**OXTR_rs237894**

>HumanReferenceGenome_UCSC__OXTR_rs237894
CATGATGTGGCCAGGCTGGTGGTGGCCAGAGTCCACGTCATCACCTCTGTGTTTGATGAGGTGCTCACATCTGGACTGGGC
TCCTGCTCCTGGCTGTGTGCTTGGGGCAGCGGGGAGGTCCCGGCTGCAGCAGCATCATGAATCCCTGCCGGTCTGTGTTTG
GATTCTTCTCAGTACCTGAGACAGAATGCTGAGGAGGACCAAGAGAAGCCCTGGGTTTTTCATGCAACACCCACCCTTCCC
TGTGAAT

>BonoboRefenceGenome_UCSC_OXTR_rs237894
CATGATGTGGCCAGGCTGGTGGTGGCCAGAGTCCACATCATCACCTCGGTGTTTGATGAGGTGCTCACATCTGGACTGGGC
TCCTGCTCCTGGCTGTGTGCTTGGGGCAGCGGGGAGGTCCCGGCTGCAGCAGCATCATGAATCCCTGCCGGTCTGTGTTTG
GATTCTTCTCAGTACCTGAGACAGAATGCTGAGGAGGACCAAGAGAAGCCCTGGGTTTTTCATGCAACACCCACCCTTCCC
TGTGAAT

>F14_OXTR_rs237894
NNNNNNNNNNNNNANNNNNTCNTCACCTCGGTGTTTGATGAGGTGCTCACATCTGGACTGGGCTCCTGCTCCTGGCTGTG
TGCTTGGGGCAGCGGGGAGGTCCCGGCTGCAGCAGCATCATGAATCCCTGCCGGTCTGTGTTTGGATTCTTCTCAGTACCT
GAGACAGAATGCTGAGGAGGACCAAGAGAAGCCCTGGGTTTTTCATGCAACACCCACCCTTCCCTGTGAATTCCTAAAGAA
GCAAAGGGAAGTAGGACCTGAAATTCTTAAAGTCCTCTTCTTTCCCTGCCCTTTGGGCTTCAGCAGGAGCCCGTCTCTACTG
AAGA

>F16_OXTR_rs237894
NNNNNNNNNNNNNNANTNNNNTNNTCACCTCGGTGTTTGATGAGGTGCTCACATCTGGACTGGGCTCCTGCTCCTGGCTG
TGTGCTTGGGGCAGCGGGGAGGTCCCGGCTGCAGCAGCATCATGAATCCCTGCCGGTCTGTGTTTGGATTCTTCTCAGTAC

CTGAGACAGAATGCTGAGGAGGACCAAGAGAAGCCCTGGGTTTTTCATGCAACACCCACCCTTCCCTGTGAATTCCTAAAG
AAGCANNACCAGTCGTGCCAGCTGCATTAATGAATCGGCCAACGCGCGGGGAGAGGCGGTTTGCGTATTGGGCGCTCTTC
CGCTTCCTCGCTCACTGACTCGCTGCGCTCGGTCGTTCGGCTGCGGCGAGCGGTATCAGCTCACTCAAAGGCGGTAATACG
GTTATCCACAGAATCAGGGGATAACGCAGGAAAGAACATGTGAGCAAAAGGCCAGCAAAAGGCCAGGAACCGTAAAAAG
GCCGCGTTGCTGGCGTTTTTCCATAGGCTCCGCCCCCCTGACGAGCATCACAAANNNNNCGCTNAGTCAGAGGTGGCGAA
ACCCGACAGGACTATAAAGATACCAGGCGTTTCCCCCTGGAAGCTCCCTCNNGCGCTCTCCTGTTCCGACCCTGCCGCTTAC
CGGATACCTGTCCGCCTTTCTCCCTTCGGGAAGCGTGGCGCTTTCTCATAGCTCACGCTGTAGGTATCTCAGTTCGGTGTANG
TCGTTCGCTCCAGCTGGGCTGNGTGCACGAANCCCCNNNNANCCCGANCGCTGCGCCTTATTCCGGTAACTATNCNNCN
TGAGNNCCACCCGGNNANAAANCNNNN

>M14_OXTR_rs237894
NNNNNNNNNNNNNTCNNNTCATCACCTCGGTGTTTGATGAGGTGCTCACATCTGGACTGGGCTCCTGCTCCTGGCTGTGT
GCTTGGGGCAGCGGGGAGGTCCCGGCTGCAGCAGCATCATGAATCCCTGCCGGTCTGTGTTTGGATTCTTCTCAGTACCTG
AGACAGAATGCTGAGGAGGACCAAGAGAAGCCCTGGGTTTTTCATGCAACACCCACCCTTCCCTGTGAATTCCTAAAGAAG
CAAAGGGAAGTAGGACCTGAAATTCTTAAAGTCCTCTTCTTTCCCTGCCCTTTGGGCTTCAGCAGGAGCCCGTCTCTACTGA
AGA

>F13_OXTR_rs237894
NNNNNNNNNNNNNNNNTCNNNTCATCACCTCGGTGTTTGATGAGGTGCTCACATCTGGACTGGGCTCCTGCTCCTGGCTG
TGTGCTTGGGGCAGCGGGGAGGTCCCGGCTGCAGCAGCATCATGAATCCCTGCCGGTCTGTGTTTGGATTCTTCTCAGTAC
CTGAGACAGAATGCTGAGGAGGACCAAGAGAAGCCCTGGGTTTTTCATGCAACACCCACCCTTCCCTGTGAATTCCTAAAG
A

>M10_OXTR_rs237894
NNNNNNNNNNNNNNNNNTNNNNTCATCACCTCGGTGTTTGATGAGGTGCTCACATCTGGACTGGGCTCCTGCTCCTGGCTG
TGTGCTTGGGGCAGCGGGGAGGTCCCGGCTGCAGCAGCATCATGAATCCCTGCCGGTCTGTGTTTGGATTCTTCTCAGTAC
CTGAGACAGAATGCTGAGGAGGACCAAGAGAAGCCCTGGGTTTTTCATGCAACACCCACCCTTCCCTGTGAATTCCTAAAG
AA

>F05_OXTR_rs237894
TCGAGGCCTNGGCATGATGTGGCCAGGCTGGTGGTGGCCAGAGTCCACATCATCACCTCGGTGTTTGATGAGGTGCTCACA
TCTGGACTGGGCTCCTGCTCCTGGCTGTGTGCTTGGGGCAGCGGGGAGGTCCCGGCTGCAGCAGCATCATGAATCCCTGCC
GGTCTGTGTTTGGATTCTTCTCAGTACCTGAGACAGAATGCTGAGGAGGACCAAGAGAAGCCCTGGGTTTTTCATGCAACA
CCCACCCTTCCCTGTGAATTCCTAAAGAAGCAAAGGGAAGTAGGACCTGAAATTCTTAAAGTCCTNNNTNNNNNCNNNNN
NNN

>M13_OXTR_rs237894
NNNNNNNNNNNNNANTNNNTCATCACCTCGGTGTTTGATGAGGTGCTCACATCTGGACTGGGCTCCTGCTCCTGGCTGTGT
GCTTGGGGCAGCGGGGAGGTCCCGGCTGCAGCAGCATCATGAATCCCTGCCGGTCTGTGTTTGGATTCTTCTCAGTACCTG
AGACAGAATGCTGAGGAGGACCAAGAGAAGCCCTGGGTTTTTCATGCAACACCCACCCTTCCCTGTGAATTCCTAAAGAAG
CAAAGGGAAGTAGGACCTGAAATTCTTAAAGTCCTCTTCTTTCCCTGCCCTTTGGGCTTCAGCAGGAGCCCGTCTCTACTGA
AGA

>F18_OXTR_rs237894
NNNNNNNNNNNNNAGTCNNNTCATCACCTCGGTGTTTGATGAGGTGCTCACATCTGGACTGGGCTCCTGCTCCTGGCTGTG
TGCTTGGGGCAGCGGGGAGGTCCCGGCTGCAGCAGCATCATGAATCCCTGCCGGTCTGTGTTTGGATTCTTCTCAGTACCT
GAGACAGAATGCTGAGGAGGACCAAGAGAAGCCCTGGGTTTTTCATGCAACACCCACCCTTCCCTGTGAATTCCTAAAGAA
GCAAAGGGAAGTAGGACCTGAAATTCTTAAAGTCCTCTTCTTTCCCTGCCCTTTGGGCTTCAGCAGGAGCCCGTCTCTACTG
AAGA

>M15_OXTR_rs237894
NNNNNNNNNNNNNNNCNNNTCATCACCTCGGTGTTTGATGAGGTGCTCACATCTGGACTGGGCTCCTGCTCCTGGCTGT
GTGCTTGGGGCAGCGGGGAGGTCCCGGCTGCAGCAGCATCATGAATCCCTGCCGGTCTGTGTTTGGATTCTTCTCAGTACC
TGAGACAGAATGCTGAGGAGGACCAAGAGAAGCCCTGGGTTTTTCATGCAACACCCACCCTTCCCTGTGAATTCCTAAAGA
AGCAAAGGGAAGTAGGACCTGAAATTCTTAAAGTCCTCTTCTTTCCCTGCCCTTTGGGCTTCAGCAGGAGCCCGTCTCTACT
GAAGA

>M12_OXTR_rs237894
NNNNNNNNNNNANTCNNNTCATCACCTCGGTGTTTGATGAGGTGCTCACATCTGGACTGGGCTCCTGCTCCTGGCTGTGT
GCTTGGGGCAGCGGGGAGGTCCCGGCTGCAGCAGCATCATGAATCCCTGCCGGTCTGTGTTTGGATTCTTCTCAGTACCTG
AGACAGAATGCTGAGGAGGACCAAGAGAAGCCCTGGGTTTTTCATGCAACACCCACCCTTCCCTGTGAATTCCNNNAGAA
GCAAAGGGAAGTAGGACCTGAAATTCTTAAAGTCCTCTTCTTTCCCTGCCCTTTGGGCTTCAGCAGGAGCCCGTCTCTACTG
AAGA

>M18_OXTR_rs237894
GNNNNNNNNNNAGTCNNTCATCACCTCGGTGTTTGATGAGGTGCTCACATCTGGACTGGGCTCCTGCTCCTGGCTGTGTG
CTTGGGGCAGCGGGGAGGTCCCGGCTGCAGCAGCATCATGAATCCCTGCCGGTCTGTGTTTGGATTCTTCTCAGTACCTGA
GACAGAATGCTGAGGAGGACCAAGAGAAGCCCTGGGTTTTTCATGCAACACCCACCCTTCCCTGTGAATTCCTAAAGAAGC
AAAGGGAAGTAGGACCTGAAATTCTTAAAGTCCTCTTCTTTCCCTGCCCTTTGGGCTTCAGCAGGAGCCCGTCTCTACTGAA
GA

>F10_OXTR_rs237894
NNNNNNNNNNNNNNNNNCNNTCATCACCTCNGTGTTTGATGAGGTGCTCACATCTGGACTGGGCTCCTGCTCCTGGCTGT
GTGCTTGGGGCAGCGGGGAGGTCCCGGCTGCAGCAGCATCATGAATCCCTGCCGGTCTGTGTTTGGATTCTTCTCAGTACC
TGAGACAGAATGCTGAGGAGGACCAAGAGAAGCCCTGGGTTTTTCATGCAACACCCACCCTTCCCTGTGAATTCCTAAAGA
AGCAAAGGGAAGTAGGACCTGAAATTCTTAAAGTCCTCTTCTTTCCCTGCCCTTTGGGCTTCAGCAGGAGCCCGTCTCTACT
GAAGA

>M19_OXTR_rs237894
NNNNNNNNNNNNNNNNNNNNTCATCNCCTCTTTGTTTGATGACGGGCTCACATCTGGACTGGGCTCCTGCTCCTGGCTGT
GTGCTTGGGGCAGCGGGGAGGTCCCGGCTGCAGCAGCATCATGAATCCCTGCCGGTCTGTGTTTGGATTCTTCTCAGTACC
TGAGACAGAATGCTGAGGAGGACCAAGAGAAGCCCTGGGTTTTTCATGCAACACCCACCCTTCCCTGTGAATTCCTAAAGA
AGCAAAGGGAAGTAGGACCTGAAATTCTTAAAGTCCTCTTCTTTCCCTGCCCTTTGGGCTTCAGCAGGAGCCCGTCTCTACT
GAAGA

**OXTR_rs237895**

>HumanReferenceGenome_UCSC_OXTR_rs237895
CTCCTCAAGACCTGGATCTGGTTTCTTCAGATCTGCCCAGGAAACAGGTGAGGCCTAAGGCCTTACTTAGAGACTTCCACTA
CTGGCTTTTCCCCAACCCCCCTCCCTCATCATGCGTCCTTCATAAGAACCCTGGCACCGGAAACGTCAGCTCTGCCTCTGACT
TGCTGTGTGACCTGGTGCAAGTGCCTTCCCCTCTCTGGGCCCTAGAACCCTCTCTGTAAGTTGTTACCAGGGG

>BonoboReferenceGenome_UCSC_OXTR_rs237895
CTCCTCAAGACCTGGATCTGGTTTCTTCAGATCTGCCCAGGAAACAGGTGAGGCCTAAGGCCTTACTTAGAGACTTCCACTA
CTGGCTTTTCCCCAACCCCCCTCCCTCATCATGCGTCCTTCATAAGAACCCTGGCACCGGAAACGTCAGCTCTGCCTCTGACT
TGCTGTGTGACCTGGTGCAAGTGCCTTCCCCTCTCTGGGCCCTAGAACCCTCTCTGTAAGTTGTTACCAGGGGCCA

>F14_OXTR_rs237895

NNNNNNNNNNNNNNNNNNNNNGNTCNAGCTCCTCAGACCTGGATCTGGTTTCTTCAGATCTGCCCAGGAAACAGGTGA
GGCCTAAGGCCTTACTTAGAGACTTCCACTACTGGCTTTTCCCCAACCCCCCTCCCTCATCATGCGTCCTTCATAAGAACCCTG
GCACCGGAAACGTCAGCTCTGCCTCTGACTTGCTGTGTGACCTGGTGCAAGTGCCTTCCCCTCTCTGGGCCCTAGAACCCTC
TCTGTAAGTTGTTACCAGGGGCCACAGGTGTTCAGTAAATATTGAATAACCAGTGTCTTGGCCACGAAAGCAAACACTGAGC
TAACTGGGAGGGTTCCATAAAGCCTGGAGN

>F16_OXTR_rs237895
NNNNNNNNNNNNNNNNNNNNNNTNNNANCNNNNANANCTGGATCTGGTTTCTTCTGATCTGCCCAGGAAACAGGTG
AGGCCNTNNGNCTTACTTAGAGACTTCCACTACTGGCTTTTCCCCAACCCCCCTCCCTCATCATGCGTCCTTCATAAGAACCC
TGGCACCGGAAACGTCAGCTCTGCCTCTGACTTGCTGTGTGACCTGGTGCAAGTGCCTTCCCCTCTCTGGGCCCTANAACCC
TCTCTGTAANTTGTTACCAGGGGCCACAGGTGTTCAGTCAATATTGAATAACCACNGTCTTGGNCACGAAAGCAAACACTG
ACCTAACTGGGAGGGTTCCNTTNNNNTNNNTCANACCNCNCNNN

>M14_OXTR_rs237895
TCTACTCCAAAGGGTGCTGGTTTCACTTTCTTGATAGTTGGTGTTCGAGCTCCTCAAGACCTGGATCTGGTTTCTTCAGATCT
GCCCAGGAAACAGGTGAGGCCTAAGGCCTTACTTAGAGACTTCCACTACTGGCTTTTCCCCAACCCCCCTCCCTCATCATGC
GTCCTTCATAAGAACCCTGGCACCGGAAACGTCAGCTCTGCCTCTGACTTGCTGTGTGACCTGGTGCAAGTGCCTTCCCCTC
TCTGGGCCCTAGAACCCTCTCTGTAAGTTGTTACCAGGGGCCACAGGTGTTCAGTAAATATTGAATAACCAGTGTCTGGCCA
CGAAAGNNNNCNNNNNNNNNN

>F13_OXTR_rs237895
NNNNNNNNNNNNNNNNNGNNNNCNAGCTCNCAGACCTGGATCTGGTTTCTTCAGATCTGCCCAGGAAACAGGTGAGGC
CTANGGCCTTACTTAGAGACTTCCACTACTGGCTTTTCCCCAACCCCCCTCCCTCATCATGCGTCCTTCATAAGAACCCTGGCA
CCGGAAACGTCAGCTCTGCCTCTGACTTGCTGTGTGACCTGGTGCAAGTGCCTTCCCCTCTCTGGGCCCTAGAACCCTCTCT
GTAAGTTGTTACCAGGGGCCACAGGTGTTCAGTAAATATTGAATAACCAGTGTCTTGGCCACGAAAGCAAACACTGAGCTA
ACTGGGAGGGTTCCATAAAGCCTGGAGTAAA

>M10_OXTR_rs237895
TTCGAGCTCCTCAAGACCTGGATCTGGTTTCTTCAGATCTGCCCAGGAAACAGGTGAGGCCTAAGGCCTTACTTAGAGACTT
CCACTACTGGCTTTTCCCCAACCCCCCTCCCTCATCATGCGTCCTTCATAAGAACCCTGGCACCGGAAACGTCAGCTCTGCCT
CTGACTTGCTGTGTGACCTGGTGCAAGTGCCTTCCCCTCTCTGGGCCCTAGAACCCTCTCTGTAAGTTGTTACCAGGGGCCA
CAGNNNCAGTANNNNNNNNNN

>F05_OXTR_rs237895
NNNNNNNNNNNGNNGTNCNNNCTCCTCAGACCTGGATCTGGTTTCTTCAGATCTGCCCAGGAAACAGGTGAGGCCTAA
GGCCTTACTTAGAGACTTCCACTACTGGCTTTTCCCCAACCCCCCTCCCTCATCATGCGTCCTTCATAAGAACCCTGGCACCG
GAAACGTCAGCTCTGCCTCTGACTTGCTGTGTGACCTGGTGCAAGTGCCTTCCCCTCTCTGGGCCCTAGAACCCTCTCTGTA
AGTTGTTACCAGGGGCCACAGGTGTTCAGTAAATATTGAATAACCAGTGTCTTGGCCACGAAAGCAAACACTGAGCTAACT
GGGAGGGTTCCATAAAGCCTGGAG

>M13_OXTR_rs237895
TNNNNNNNNNGGTGTTCGAGCTCCTCAGACCTGGATCTGGTTTCTTCAGATCTGCCCAGGAAACAGGTGAGGCCTAAGGC
CTTACTTAGAGACTTCCACTACTGGCTTTTCCCCAACCCCCCTCCCTCATCATGCGTCCTTCATAAGAACCCTGGCACCGGAA
ACGTCAGCTCTGCCTCTGACTTGCTGTGTGACCTGGTGCAAGTGCCTTCCCCTCTCTGGGCCCTAGAACCCTCTCTGTAAGTT
GTTACCAGGGGCCACAGGTGTTCAGTAAATATTGAATAACCAGTGTCTTGGCCACGAAAGCAAACACTGAGCTAACTGGGA
GGGTTCCATAAAGCCCCTGGAGTA

>F18_OXTR_rs237895

CTTCCACTACTGGCTTTTCCCCAACCCCCCTCCCTCATCATGCGTCCTTCATAAGAACCCTGGCACCGGAAACGTCAGCTCTG
CCTCTGACTTGCTGTGTGACCTGGTGCAAGTGCCTTCCCCTCTCTGGGCCCTAGAACCCTCTCTGTAAGTTGTTACCAGGGG
CCACAGNNNNCAGTNNNNNNNNNNNN

>M15_OXTR_rs237895
NNNNNNNNNNNNNNNNNNNNNNNNNNNNNAGCTCCTCAAGACCTGGATCTGGTTTCTTCAGATCTGCCCAGGAAACA
GGTGAGGCCTAAGGCCTTACTTAGAGACTTCCACTACTGGCTTTTCCCCAACCCCCCTCCCTCATCATGCGTCCTTCATAAGA
ACCCTGGCACCGGAAACGTCAGCTCTGCCTCTGACTTGCTGTGTGACCTGGTGCAAGTGCCTTCCCCTCTCTGGGCCCTAGA
ACCCTCTCTGTAAGTTGTTACCAGGGGCCACAGGTGTTCAGTAAATATTGAATAACCAGTGTCTTGGCCACGAAAGCAAACA
CTGAGCTAACTGGGAGGGTTCCATAAAGCCTGGAGTA

>M12_OXTR_rs237895
NNNNNNNNNNNNNNGGNGNNNNAGCTCCTCANGACCTGGATCTGGTTTCTTCAGATCTGCCCAGGAAACAGGTGAGGC
CTAAGGCCTTACTTAGAGACTTCCACTACTGGCTTTTCCCCAACCCCCCTCCCTCATCATGCGTCCTTCATAAGAACCCTGGCA
CCGGAAACGTCAGCTCTGCCTCTGACTTGCTGTGTGACCTGGTGCAAGTGCCTTCCCCTCTCTGGGCCCTAGAACCCTCTCT
GTAAGTTGTTACCAGGGGCCACAGGTGTTCAGTAAATATTGAATAACCAGTGTCTTGGCCACGAAAGCAAACACTGAGCTA
ACTGGGAGGGTTCCATAAAGCCTGGAGTA

>M18_OXTR_rs237895
CTGCCCAGGAAACAGGTGAGGCCTAAGGCCTTACTTAGAGACTTCCACTACTGGCTTTTCCCCAACCCCCCTCCCTCATCAT
GCGTCCTTCATAAGAACCCTGGCACCGGAAACGTCAGCTCTGCCTCTGACTTGCTGTGTGACCTGGTGCAAGTGCCTTCCCC
TCTCTGGGCCCTAGAACCCTCTCTGTAAGTTGTTACCAGGGGCCACAGGNNNCAGTAANNNNNNNNNNNN

>F10_OXTR_rs237895
NNNNNNNNNNNNNNNNNNNNNNNNNNNNNNNNCTCTCNGACCTGGATCTGGTTTCTTCAGATCTGCCCAGGAAACAGG
TGAGGCCTAAGGCCTTACTTAGAGACTTCCACTACTGGCTTTTCCCCAACCCCCCTCCCTCATCATGCGTCCTTCATAAGAACC
CTGGCACCGGAAACGTCAGCTCTGCCTCTGACTTGCTGTGTGACCTGGTGCAAGTGCCTTCCCCTCTCTGGGCCCTAGAACC
CTCTCTGTAAGTTGTTACCAGGGGCCACAGGTGTTCAGTAAATATTGAATAACCAGTGTCTTGGCCACGAAAGCAAACACTG
AGCTAACTGGGAGGGTTCCATAAAGCCTGGAGTA

>M19_OXTR_rs237895
NNNNNNNNNNNNNNNNNNNNNNNNNNNANNTCCTCANACCTGGATCTGGTTTCTTCAGATCTGCCCAGGAAACAGGTG
AGGCCTAAGGCCTTACTTAGAGACTTCCACTACTGGCTTTTCCCCAACCCCCCTCCCTCATCATGCGTCCTTCATAAGAACCCT
GGCACCGGAAACGTCAGCTCTGCCTCTGACTTGCTGTGTGACCTGGTGCAAGTGCCTTCCCCTCTCTGGGCCCTAGAACCCT
CTCTGTAAGTTGTTACCAGGGGCCACAGGTGTTCAGTAAATATTGAATAACCAGTGTCTTGGCCACGAAAGCAAACACTGAG
CTAACTGGGAGGGTTCCATAAAGCCTGGAGTA

**OXTR_rs2270463**

>HumanReferenceGenome_UCSC_OXTR_rs2270463
AGGCACTGGCAATCCAACGAGAAGCCCCACTACATCGCATTCCAGCTCCTGAAGCCATTCCCTGGGTAAGAGAGCCCTGATT
GTTATTATCGTCTCTCCATTTTACAGATGGGGACAGAGTGGCCTAGAGAGGTGAAGTGTGCCAAAGGTCACACAGCCAGCC
AGAGGCAGAGGAGGGACTAACCCCACTTCCCTCCATGGGCAGTGCGCCCCAAAGGAGCCAAAATGCTGACAGCAGTGGG
GCAAATTCCAGGG

>BonoboReferenceGenome_UCSC_OXTR_rs2270463
AGGCACTGGCAATCCAACGAGAAGCCCCACTACATCGCATTCCAGCTCCTGAAGCCATTCCCTGGGTAAGAGAGCCCTGATT
GTTATTATCGTCTCTCCATTTTACAGATGGGGACAGAGTGGCCTAGAGAGGTGAAGTGTGCCAAAGGTCACACAGCCAGCC

AGAGGCAGAGGAGGGACTAACCCCACTTCCCTCCATGGGCAGTGCGCCCCAAAGGAGCCAAAATGCTGACAGCAGTGGG
GCAAATTCCAGGG

>F14_OXTR_rs2270463
GNNNNNNNNNTNNNNNNNNANANAGCCCTGATTGTTATTATCGTCTCTCCATTTTACAGATGGGGACAGAGTGGCCTAG
AGAGGTGAAGTGTGCCAAAGGTCACACAGCCAGCCAGAGGCAGAGGAGGGACTAACCCCACTTCCCTCCATGGGCAGTG
CGCCCCAAAGGAGCCAAAATGCTGACAGCAGTGGGGCAAATTCCAGGGGCCCCGGAGCGACAGCCGAGTTGGCCACAGC
CACACGATGCATAGGAAA

>F16_OXTR_rs2270463
NNNNNNNNNNCNNNNNAGANAGCCCTGATTGTTATTATCGTCTCTCCATTTTACAGATGGGGACAGAGTGGCCTAGAGA
GGTGAAGTGTGCCAAAGGTCACACAGCCAGCCAGAGGCAGAGGAGGGACTAACCCCACTTCCCTCCATGGGCAGTGCGC
CCCAAAGGAGCCAAAATGCTGACAGCAGTGGGGCAAATTCCAGGGGCCCCGGAGCGACAGCCGAGTTGGCCACAGCCAC
ACGATGCATAGGAAA

>M14_OXTR_rs2270463
NNNNNNNNNNCNNGGNANAGAGCCCTGATTGTTATTATCGTCTCTCCATTTTACAGATGGGGACAGAGTGGCCTAGAGA
GGTGAAGTGTGCCAAAGGTCACACAGCCAGCCAGAGGCAGAGGAGGGACTAACCCCACTTCCCTCCATGGGCAGTGCGC
CCCAAAGGAGCCAAAATGCTGACAGCAGTGGGGCAAATTCCAGGGGCCCCGGAGCGACAGCCGAGTTGGCCACAGCCAC
ACGATGCATAGGAAA

>F13_OXTR_rs2270463
NNNNNNANNNCNNNNNAGAGAGCCCTGATTGTTATTATCGTCTCTCCATTTTACAGATGGGGACAGAGTGGCCTAGAGA
GGTGAAGTGTGCCAAAGGTCACACAGCCAGCCAGAGGCAGAGGAGGGACTAACCCCACTTCCCTCCATGGGCAGTGCGC
CCCAAAGGAGCCAAAATGCTGACAGCAGTGGGGCAAATTCCAGGGGCCCCGGAGCGACAGCCGAGTTGGCCACAGCCAC
ACGATGCATAGGAAA

>M10_OXTR_rs2270463
TCCAGCCTGCCAAGCGTAGTTGAAAANTAACAACTCCCAGTAGCACTAGGGCGGATGCAGCTGTTGAGTGAAAACGCGGG
AGTTATCACGAAGTTAGGGCGAAAGGAAGGGTGGCACTCCNTAGTGCGTCATAAGCTAGCTTGCATGCNTGAGAATTTCAG
AATGTAATGATACCTTTGTTAGATAAGGCTGCTTACNTTTGCTTGCCAGCTCNTGAAGCCATTCCCTGGGTAAGAGAGCCATG
ATTGTTATTATNGTCTCTCCATTTTACAGATGGGGACAGAGTGGCCTAGAGAGGTGAAGTGTGCCAAAGGTCACACAGCCA
GCCAGAGGCAGAGGAGGGACTAACCCCACTTCCCTCCATGGGCAGTGCGCCCCAAAGGAGCCAAAATGCTGACAGCAGT
GGGGCAAANCCAGNNCCNCGANNNNNNNNNNNNNNNN

>F05_OXTR_rs2270463
NNNNNTNNNNGGNAGAGAGCCCTGATTGTTATTATCGTCTCTCCATTTTACAGATGGGGACAGAGTGGCCTAGAGAGGTG
AAGTGTGCCAAAGGTCACACAGCCAGCCAGAGGCAGAGGAGGGACTAACCCCACTTCCCTCCATGGGCAGTGCGCCCCAA
AGGAGCCAAAATGCTGACAGCAGTGGGGCAAATTCCAGGGGCCCCGGAGCGACAGCCGAGTTGGCCACAGCCACACGAT
GCATAGGAAA

>M13_OXTR_rs2270463
NNNNNNNNNNNNNNNNNNNNNNNNAGANAGCCCTGATTGTTATTATCGTCTCTCCATTTTACAGATGGGGACAGAGTGGC
CTAGAGAGGTGAAGTGTGCCAAAGGTCACACAGCCAGCCAGAGGCAGAGGAGGGACTAACCCCACTTCCCTCCATGGGC
AGTGCGCCCCAAAGGAGCCAAAATGCTGACAGCAGTGGGGCAAATTCCAGGGGCCCCGGAGCGACAGCCGAGTTGGCCA
CAGCCACACGATGCATAGGAAA

>F18_OXTR_rs2270463

NNNNNNNNNNNNNNNNNTCNNNNNNAGAGAGCCCTGATTGTTATTATCGTCTCTCCATTTTACAGATGGGGACAGAGTGG
CCTAGAGAGGTGAAGTGTGCCAAAGGTCACACAGCCAGCCAGAGGCAGAGGAGGGACTAACCCCACTTCCCTCCATGGG
CAGTGCGCCCCAAAGGAGCCAAAATGCTGACAGCAGTGGGGCAAATTCCAGGGGCCCCGGAGCGACAGCCGAGTTGGCC
ACAGCCACACGATGCATAGGAAA

>M15_OXTR_rs2270463
NNNNNNNNNNNNNCNNGNNNNAGANNAGCCCTGATTGTTATTATCGTCTCTCCATTTTACAGATGGGGACAGAGTGGCCT
AGAGAGGTGAAGTGTGCCAAAGGTCACACAGCCAGCCAGAGGCAGAGGAGGGACTAACCCCACTTCCCTCCATGGGCAG
TGCGCCCCAAAGGAGCCAAAATGCTGACAGCAGTGGGGCAAATTCCAGGGGCCCCGGAGCGACAGCCGAGTTGGCCACA
GCCACACGATGCATAGGAAA

>M12_OXTR_rs2270463
NNNNNNNNNNCNNGNNAGAGAGCCCTGATTGNTATTATCGTCTCTCCATTTTACAGATGGGGACAGAGTGGCCTAGAGA
GGTGAAGTGTGCCAAAGGTCACACAGCCAGCCAGAGGCAGAGGAGGGACTAACCCCACTTCCCTCCATGGGCAGTGCGC
CCCAAAGGAGCCAAAATGCTGACAGCAGTGGGGCAAATTCCAGGGGCCCCGGAGCGACAGCCGAGTTGGCCACAGCCAC
ACGATGCATAGGAAA

>M18_OXTR_rs2270463
NNNNNNANNNCNNNNNAGAGAGCCCTGATTGNTATTATCGTCTCTCCATTTTACAGATGGGGACAGAGTGGCCTAGAGA
GGTGAAGTGTGCCAAAGGTCACACAGCCAGCCAGAGGCAGAGGAGGGACTAACCCCACTTCCCTCCATGGGCAGTGCGC
CCCAAAGGAGCCAAAATGCTGACAGCAGTGGGGCAAATTCCAGGGGCCCCGGAGCGACAGCCGAGTTGGCCACAGCCAC
ACGATGCATAGGAAA

>F10_OXTR_rs2270463
NNNNNNNNNNNNNNNNNNAGANAGCCCTGATTGTTATTATCGTCTCTCCATTTTACAGATGGGGACAGAGTGGCCTAGAG
AGGTGAAGTGTGCCAAAGGTCACACAGCCAGCCAGAGGCAGAGGAGGGACTAACCCCACTTCCCTCCATGGGCAGTGCG
CCCCAAAGGAGCCAAAATGCTGACAGCAGTGGGGCAAATTCCAGGGGCCCCGGAGCGACAGCCGAGTTGGCCACAGCCA
CACGATGCATAGGAAA

>M19_OXTR_rs2270463
NNNNNNNNNCNNNNNAGAGAGCCCTGATTGTTATTATCGTCTCTCCATTTTACAGATGGGGACAGAGTGGCCTAGAGAG
GTGAAGTGTGCCAAAGGTCACACAGCCAGCCAGAGGCAGAGGAGGGACTAACCCCACTTCCCTCCATGGGCAGTGCGCCC
CAAAGGAGCCAAAATGCTGACAGCAGTGGGGCAAATTCCAGGGGCCCCGGAGCGACAGCCGAGTTGGCCACAGCCACAC
GATGCATAGGAAA

**AVPR1A_ rs3803107**

>HumanReferenceGenome_UCSC_AVPR1A_rs3803107
ATTAATTTTAATGGATGAAAACATAATTTCTCTGAAGTTTTCTCTGAATATTAAGACTGATGATGAGCTCTCTTTGTTCAGAAAT
AGTGCCGCATTTTATGTGACTTTTAAACCAATCAAGTCTTACTTATGTTAGAAATGAAAATAAAAGAAACTAACAACAAAATAT
AAAGCTAGGGTGGTTATGATTTTTCCTTTAAAAATATAACTTCTGTTGTATTTCTGGGAATCAGACCTGCATATTTTAGTA

>BonoboReferenceGenome_UCSC_AVPR1A_rs3803107
ACCATCCCATCTCCTGGACACTGTTTAAGGCTGCATTTTCTGATGCACAAATTAATTTTAATGGATGAAAACATAATTTCTCTG
AAGTTTTCTCTGAATATTAAGACTGATGATGAGCTCTCTTTGTTCAGAAATAGTGCCACATTTTATGTGACTTTTAAACCAATC
AAGTCTTACTTATGTTAGAAATGAAAATAAAAGAAACCAACAACAAAATATAAAGCTAGGGTGGTTATGAT

>F14_AVPR1A

NNNNNNNNNNTCTCTGGNNNTGTTNNNGCTGCATTTTCTGATGCACAAATTAATTTTAATGGATGAAAACATAATTTCTCT
GAAGTTTTCTCTGAATATTAAGACTGATGATGAGCTCTCTTTGTTCAGAAATAGTGCCACATTTTATGTGACTTTTAAACCAAT
CAAGTCTTACTTATGTTAGAAATGAAAATAAAAGAAACCAACAACAAAATATAAAGCTAGGGTGGTTATGATTTTTCCTTTAA
AAATATAACTTCTGTTGTATTTCTGGGAATCNACCTGCANNGN

>F16_AVPR1A_rs3803107
NNNNNNNNCCNTANNNNNNNNNTGNTTTAGGCTGCATTTTCTGATGCACAAATTAATTTTAATGGATGAAAACATAATTTCT
CTGAAGTTTTCTCTGAATATTAAGACTGATGATGAGCTCTCTTTGTTCAGAAATAGTGCCACATTTTATGTGACTTTTAAACCA
ATCAAGTCTTACTTATGTTAGAAATGAAAATAAAAGAAACCAACAACAAAATATAAAGCTAGGGTGGTTATGATTTTTCCTTTA
AAAATATAACTTCTGTTGTATTTCTGGGAATCNNNCCTGCATA

>M14_AVPR1A_rs3803107
NNNNNNNNNNNNNNNNTCNGGANNTGTTTAAGGCTGCATTTTCTGATGCACAAATTAATTTTAATGGATGAAAACATAATT
TCTCTGAAGTTTTCTCTGAATATTAAGACTGATGATGAGCTCTCTTTGTTCAGAAATAGTGCCACATTTTATGTGACTTTTAAAC
CAATCAAGTCTTACTTATGTTAGAAATGAAAATAAAAGAAACCAACAACAAAATATAAAGCTAGGGTGGTTATGATTTTTCCT
TTAAAAATATAACTTCTGTTGTATTTCTGGGAATCAGACCTGCAN

>F13_AVPR1A_rs3803107
NNNNNNNNNNNNTNATNNTGNNNNTGTTTAAGGCTGCATTTTCTGATGCACAAATTAATTTTAATGGATGAAAACATAATT
TCTCTGAAGTTTTCTCTGAATATTAAGACTGATGATGAGCTCTCTTTGTTCAGAAATAGTGCCACATTTTATGTGACTTTTAAAC
CAATCAAGTCTTACTTATGTTAGAAATGAAAATAAAAGAAACCAACAACAAAATATAAAGCTAGGGTGGTTATGATTTTTCCT
TTAAAAATATAACTTCTGTTGTATTTCTGG

>M10_AVPR1A_rs3803107
NNNNNNNNNNNNTCTCNGNANNTGTNNAGGCTGCATTTTCTGATGCACAAATTAATTTTAATGGATGAAAACATAATTTCTC
TGAAGTTTTCTCTGAATATTAAGACTGATGATGAGCTCTCTTTGTTCAGAAATAGTGCCACATTTTATGTGACTTTTAAACCAA
TCAAGTCTTACTTATGTTAGAAATGAAAATAAAAGAAACCAACAACAAAATATAAAGCTAGGGTGGTTATGATTTTTCCTTTA
AAAATATAACTTCTGTTGTATTTCTGGGAATCAGACCTGCATA

>F05_AVPR1A_rs3803107
ATNTCTTTGCGGGGNGGGGAAAAGAATAAGGTGTTCCCCNNGCCAACAGNTAAAANCCATGTTCACTTATCCCANACANT
TTGCGCTTNATATGTTCTCCCCGCNAGCGGGGGTAACCATNNTATGNTATTCTATNNCCCTCTCACAGTGCTCCCCGCNCCAC
CGAGNNATTCCGCTCNGAAATACAGCNNTNACCAGGGTNCACNAGNNCCCCNGGCCTACACCCCAGGCACTTGTACTCCT
AGGAGGTACCATCCCATCTCCTGGACACTGTTTAAGGCTGCATTTTCTGATGCACAAATTAATTTTAATGGATGAAAACATAAT
TTCTCTGAAGTTTTCTCTGAATATTAAGACTGATGATGAGCTCTCTTTGTTCAGAAATAGTGCCACATTTTATGTGACTTTTAAA
CCAATCAAGTCTTACTTATGTTAGAAATGAAAATAAAAGAAACCAACAACAAAATATAAAGCTAGGGTGGTTANGATTNCCT
TNANNNNNNNNNN

>M13_AVPR1A_rs3803107
NNNNNNNNNNNNNNNNNAANNNNNNNNNNNNGNNTTAGGCTGCATTTTCTGATGCACAAATTAATTTTAATGGATGAAAA
CATAATTTCTCTGAAGTTTTCTCTGAATATTAAGACTGATGATGAGCTCTCTTTGTTCAGAAATAGTGCCACATTTTATGTGACT
TTTAAACCAATCAAGTCTTACTTATGTTAGAAATGAAAATAAAAGAAACCAACAACAAAATATAAAGCTAGGGTGGTTATGAT
TTTTCCTTTAAAAATATAACTTCTGTTGTATTTCTGGGA

>F18_AVPR1A_rs3803107
NNNNGNNNNNCNNNNNCCTGGANNTGTTTAAGGCTGCATTTTCTGATGCACAAATTAATTTTAATGGATGAAAACATAAT
TTCTCTGAAGTTTTCTCTGAATATTAAGACTGATGATGAGCTCTCTTTGTTCAGAAATAGTGCCACATTTTATGTGACTTTTAAA
CCAATCAAGTCTTACTTATGTTAGAAATGAAAATAAAAGAAACCAACAACAAAATATAAAGCTAGGGTGGTTATGATTTTTCC
TTTAAAAATATAACTTCTGTTGTATTTCTGGGA

>M15_AVPR1A_rs3803107
NNNNNNNNCCNTCTCNGGANCTGTTNNNGCTGCATTTTCTGATGCACAAATTAATTTTAATGGATGAAAACATAATTTCTCT
GAAGTTTTCTCTGAATATTAAGACTGATGATGAGCTCTCTTTGTTCAGAAATAGTGCCACATTTTATGTGACTTTTAAACCAAT
CAAGTCTTACTTATGTTAGAAATGAAAATAAAAGAAACCAACAACAAAATATAAAGCTAGGGTGGTTATGATTTTTCCTTTAA
AAATATAACTTCTGTTGTATTTCTGGGAATCAGACCTGCAN

>M12_AVPR1A_rs3803107
NNNNNNNNNNNNNNNNTCCTGGNNCTGTTTAGGCTGCATTTTCTGATGCACAAATTAATTTTAATGGATGAAAACATAATTT
CTCTGAAGTTTTCTCTGAATATTAAGACTGATGATGAGCTCTCTTTGTTCAGAAATAGTGCCACATTTTATGTGACTTTTAAAC
CAATCAAGTCTTACTTATGTTAGAAATGAAAATAAAAGAAACCAACAACAAAATATAAAGCTAGGGTGGTTATGATTTTTCCT
TTAAAAATATAACTTCTGTTGTATTTCTGGGA

>M18_AVPR1A_rs3803107
NNNNNNNNNNNNTCTCTGNNNCTGTNNNNGNTGCATTTTCTGATGCACAAATTAATTTTAATGGATGAAAACATAATTTCTC
TGAAGTTTTCTCTGAATATTAAGACTGATGATGAGCTCTCTTTGTTCAGAAATAGTGCCACATTTTATGTGACTTTTAAACCAA
TCAAGTCTTACTTATGTTAGAAATGAAAATAAAAGAAACCAACAACAAAATATAAAGCTAGGGTGGTTATGATTTTTCCTTTA
AAAATATAACTTCTGTTGTATTTCTGGGAATCAGACCTGCAT

>F10_AVPR1A_rs3803107
NNNNNNNNNNCNTCTCNGNANNTGTTTAGGCTGCATTTTCTGATGCACAAATTAATTTTAATGGATGAAAACATAATTTCT
CTGAAGTTTTCTCTGAATATTAAGACTGATGATGAGCTCTCTTTGTTCAGAAATAGTGCCACATTTTATGTGACTTTTAAACCA
ATCAAGTCTTACTTATGTTAGAAATGAAAATAAAAGAAACCAACAACAAAATATAAAGCTAGGGTGGTTATGATTTTTCCTTTA
AAAATATAACTTCTGTTGTATTTCTGGGAATCAGACCTGCAN

>M19_AVPR1A_rs3803107
NNNNNNNNCNNNTCTNNTGGANNTGTTTAAGGCTGCATTTTCTGATGCACAAATTAATTTTAATGGATGAAAACATAATTT
CTCTGAAGTTTTCTCTGAATATTAAGACTGATGATGAGCTCTCTTTGTTCAGAAATAGTGCCACATTTTATGTGACTTTTAAAC
CAATCAAGTCTTACTTATGTTAGAAATGAAAATAAAAGAAACCAACAACAAAATATAAAGCTAGGGTGGTTATGATTTTTCCT
TTAAAAATATAACTTCTGTTGTATTTCTGGGAATCAGACCTGCAN
