## Supplementary Material 5 for "Linked *OXTR* Variants Are Associated With Social Behavior Differences in Bonobos (*Pan paniscus*)"

| Bonobo ID | F14 | F16 | M14 | F13 | M10 | F05 | M13 | F18 | M15 | M12 | M18 | F10 | M19 |
|---|---|---|---|---|---|---|---|---|---|---|---|---|---|
| **F14** | 1 | 0 | 0 | 0.125 | 0 | 0 | 0.125 | 0.031 | 0 | 0.063 | 0.094 | 0 | 0.25 |
| **F16** | 0 | 1 | 0 | 0 | 0.031 | 0 | 0 | 0.016 | 0 | 0 | 0 | 0 | 0 |
| **M14** | 0 | 0 | 1 | 0 | 0 | 0 | 0 | 0 | 0.125 | 0 | 0 | 0 | 0.25 |
| **F13** | 0.125 | 0 | 0 | 1 | 0 | 0 | 0.25 | 0.031 | 0 | 0.188 | 0.344 | 0 | 0.063 |
| **M10** | 0 | 0.031 | 0 | 0 | 1 | 0 | 0 | 0.063 | 0 | 0.063 | 0.031 | 0 | 0 |
| **F05** | 0 | 0 | 0 | 0 | 0 | 1 | 0 | 0 | 0.25 | 0 | 0 | 0 | 0 |
| **M13** | 0.125 | 0 | 0 | 0.25 | 0 | 0 | 1 | 0.031 | 0 | 0.188 | 0.219 | 0 | 0.063 |
| **F18** | 0.031 | 0.016 | 0 | 0.031 | 0.063 | 0 | 0.031 | 1 | 0 | 0.031 | 0.031 | 0 | 0.016 |
| **M15** | 0 | 0 | 0.125 | 0 | 0 | 0.25 | 0 | 0 | 1 | 0 | 0 | 0 | 0.063 |
| **M12** | 0.063 | 0 | 0 | 0.188 | 0.063 | 0 | 0.188 | 0.031 | 0 | 1 | 0.344 | 0 | 0.031 |
| **M18** | 0.094 | 0 | 0 | 0.344 | 0.031 | 0 | 0.219 | 0.031 | 0 | 0.344 | 1 | 0 | 0.047 |
| **F10** | 0 | 0 | 0 | 0 | 0 | 0 | 0 | 0 | 0 | 0 | 0 | 1 | 0 |
| **M19** | 0.25 | 0 | 0.25 | 0.063 | 0 | 0 | 0.063 | 0.016 | 0.063 | 0.031 | 0.047 | 0 | 1 |

**Supplementary Material 5**: Cell value is the kinship coefficient between the corresponding individuals. Each individual has a kinship coefficient of 1 with themselves.
